# Brain-Resident CD8^+^ T Cells Regulate Neuronal Activity and Behavior via Interferon-Gamma

**DOI:** 10.64898/2026.08.24.746768

**Authors:** Kyungsoo Park, Juneil Jang, Shin Jeon, Sunsook Hwang, Kyuhyun Choi, Timothy O. Cox, Shin Foong Ngiow, Julia N. Flores, Camille F. Harrison, Shuaitong Liu, F. Chris Bennett, Michael A. Silverman, John E. Wherry, Christoph A. Thaiss, Marc V. Fuccillo, Yeong Shin Yim

## Abstract

Maintaining brain homeostasis is crucial for proper function of the central nervous system and has traditionally been attributed to neuronal and glial interactions. However, recent research highlights the essential role of brain-resident immune cells in this process. Our study characterizes brain-specific CD8^+^ T cells and elucidates their significant contribution to brain homeostasis and behavior. We identified a distinct population of CD8^+^ T cells that infiltrates the brain during early development, undergoes clonal expansion, and acquires effector memory-like characteristics through interactions with microglia. Notably, the absence of these cells results in hyperactivation of neuronal activity and abnormal behaviors, due to loss of regulation of interferon-gamma (IFN-γ) secreted by CD8^+^ T cells. Our findings demonstrate that IFN-γ secreting brain-specific CD8^+^ T cells are crucial for maintaining the physiological level of neuronal excitability and normal behavioral patterns. This study provides novel insights into neuroimmune interactions, emphasizing the critical role of CD8^+^ T cells in sustaining brain function and behavior.

## Introduction

Maintaining homeostasis in the brain is essential for the proper functioning of the central nervous system (CNS). This balance is achieved through dynamic interactions among various cell types, including neurons, glia, and immune cells that reside in the brain and at the brain border. Homeostatic plasticity, an intrinsic property of neural networks, enables the brain to maintain stable activity levels despite fluctuations in external and internal conditions^1–3^. Recent research has highlighted the significant role of immune cells, particularly T cells, in maintaining brain homeostasis^4–10^. Traditionally, T cells, especially CD8^+^ T cells, have been widely recognized as potent antiviral effectors due to their ability to eradicate viral reservoirs. They accomplish this by secreting cytotoxic agents such as perforin, which forms pores in the membranes of infected cells, and granzyme B, which enters these cells to trigger apoptosis^11,12^. This cytotoxic capability is crucial for controlling viral infections like Human Immunodeficiency Virus (HIV), where CD8^+^ T cells target and destroy infected cells, thereby reducing the viral load^13,14^. In the brain, increased infiltration of CD8^+^ T cells has been observed in neurodegenerative diseases like multiple sclerosis (MS), Alzheimer’s disease (AD), and Parkinson’s disease (PD), indicating a contribution of CD8^+^ T cell to CNS damage and neuronal death^15–20^. For instance, in MS, CD8^+^ T cells invade the CNS, attacking myelin sheaths and contributing to the demyelination process^17–19^. Similarly, in AD, CD8^+^ T cells have been found around amyloid plaques, suggesting a potential role in exacerbating neuronal death^15,16^. Beside their roles in neurodegenerative disease, CD8+ T cells may also participate in regulating brain function under non-pathological but physiologically challenging conditions. For example, exposure to an enriched environment (EE)-a well-established paradigm that induces neural plasticity and cognitive enhancement-has been shown to require CD8^+^ T cells for its beneficial effects on hippocampal function^21^. Brain-resident CD8^+^ T cells in the healthy brain exhibit distinct phenotypic characteristics compared to their peripheral counterparts and when these are depleted, EE-induced improvements in hippocampus-dependent behavior, neurogenesis in the dentate gyrus, and functional synaptic plasticity were impaired. These findings suggest that, even in the absence of overt inflammation, CD8^+^ T cells can modulate neural circuit remodeling in response to environmental stimuli, highlighting their broader role in CNS adaptation.

However, contrary to the conventional view of CD8^+^ T cells as effector cells active under immune or environmental stimulation, recent studies have identified their presence in both humans and mice under homeostatic conditions, suggesting that they may play alternative roles. These CD8^+^ T cells do not exhibit the typical cytotoxic profile^4–10^. For example, brain-specific CD8^+^ T cells in human white matter show increased expression of the tissue resident marker CD69 and low expression of cytolytic enzymes, while maintaining polyfunctionality upon activation. These unique characteristics suggest that CD8^+^ T cells in the human brain may provide immune surveillance, protecting against neurotropic virus reactivation while being tightly regulated by key immune checkpoint molecules^4^. Similarly, CD8^+^ T cells have also been detected in the brains of mice under steady-state conditions^5–10^. Notably, these CD8^+^ T cells appear to be memory T cells, as indicated by their low expression of CD62L and high expression of CD44^5^. However, it is not yet well understood when CD8^+^ T cells come to the brain, why CD8^+^ T cells are present in the brain under non-pathological conditions, or what role these cells play in the brain’s immune environment.

Our study specifically investigates the role of brain-resident CD8^+^ T cells in maintaining physiological brain function. We characterized CD45^hi^ immune cells across the lifespan of mice and observed that CD8^+^ T cells infiltrate the brain early in development, undergo clonal expansion, and acquire effector memory-like characteristics. Importantly, the absence of these cells leads to increased neuronal activity and abnormal behaviors, due to loss of regulatory mechanisms mediated by Interferon-gamma (IFN-γ). This work provides new insights into how brain-specific T cells function both in maintaining brain health, emphasizing their dual role in neuroimmune interactions.

## Results

### The brain possesses its own immune system, similar to other non-lymphoid organs

Recent research has suggested the presence of immune cells, including lymphocytes, within the brain^4–6,9,10,22^. A remaining question is whether these immune cells reside within the brain parenchyma and/or at brain borders. To address this, we adopted a protocol to isolate brain-specific CD45^hi^ immune cells, distinct from those in circulation and surrounding brain regions. Given the abundance of small blood vessels spread out in the brain, we segregated immune cells within the brain from those derived from blood circulation^23^. Circulating immune cells were labeled with an anti-CD45 (clone 30-F11) antibody via tail vein injection before brain tissue extraction from wild-type (WT) mice, facilitating the discrimination of brain-resident immune cells from circulating ones. After blood collection, the mice were perfused with cold PBS, and the brain and spleen were collected. The dural mater and choroid plexus (ChP) were isolated from the skull cap and brain, respectively. We then enriched immune cells from each brain compartment through Percoll gradient centrifugation following tissue digestion with Collagenase and DNase. Isolated immune cells from the blood and spleen, as well as enriched immune cells from the brain compartments, were then analyzed via flow cytometry (Figure 1A, and Figure S1-2).

**Figure 1.**
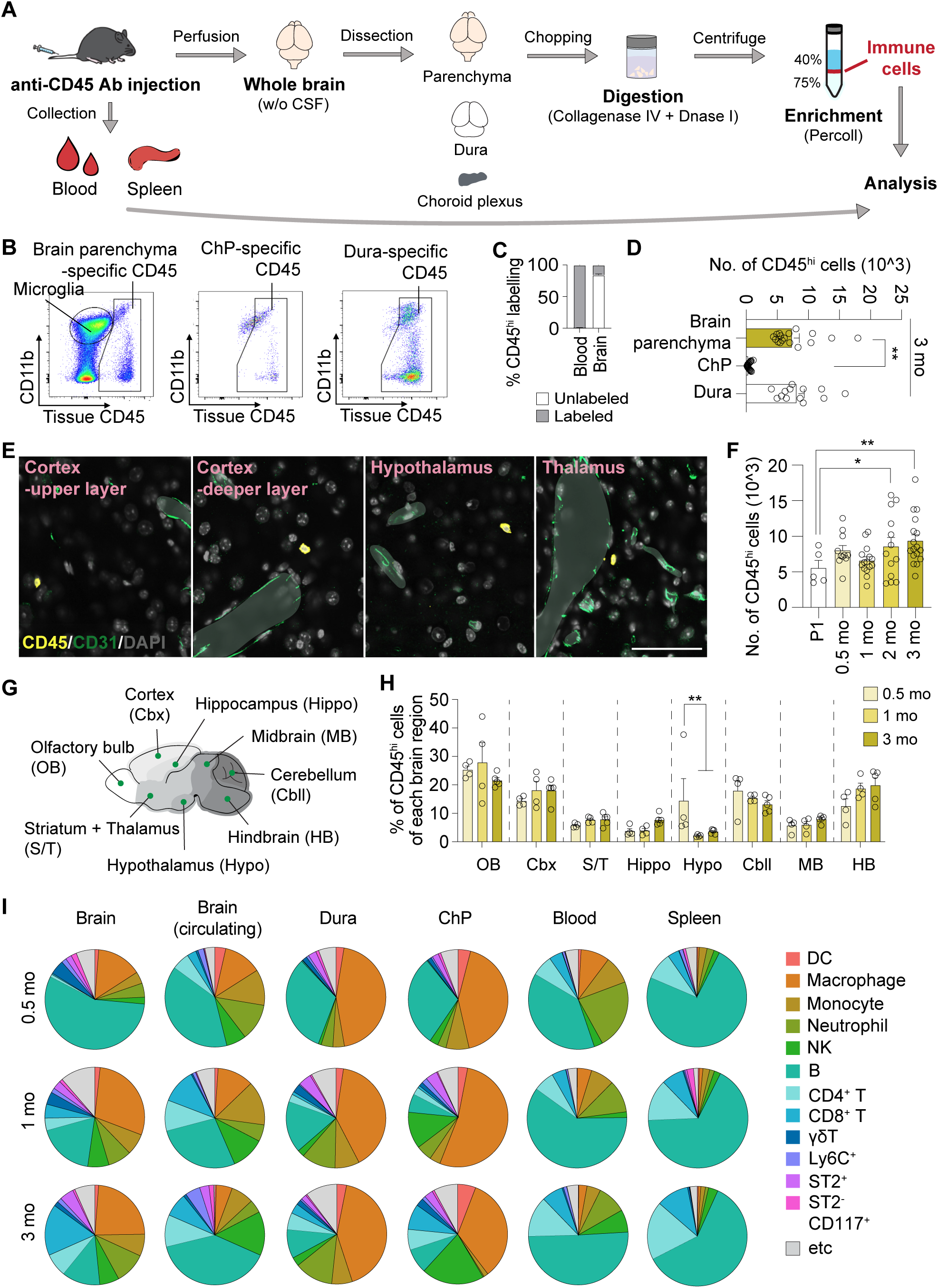
Resident immune cells were observed in the brain, similar to those in other non-lymphoid organs. (**A**) Schematic illustrating the experimental workflow for isolating immune cells from the blood, spleen, brain parenchyma, dura, and choroid plexus (ChP). (**B**) Representative FACS plots showing the gating strategy used to isolate CD45^hi^ cells in the brain parenchyma, ChP, and dura. (**C**) Quantification of CD45^hi^ immune cells in the brain parenchyma, ChP, and dura of 3-month (mo)-old mice. (**D**) Representative images showing CD45^hi^ immune cells located beyond the glia limitans in the upper and deeper layers of the cerebral cortex, hypothalamus, and thalamus. (**E**) Absolute numbers of CD45^hi^ immune cells in the brain at postnatal day 1 (P1), 0.5 mo, 1 mo, 2 mo, and 3 mo, as measured by flow cytometry. (**F**) Schematic showing the sub-regions of the mouse brain. (**G**) Relative distribution of CD45^hi^ immune cells across mouse brain regions, determined by flow cytometry. The relative distribution was calculated as the number of CD45^hi^ immune cells in each region of brain divided by the total number of CD45^hi^ immune cells in whole brain. (**H**) Pie chart depicting the distribution of tissue-resident and circulating blood immune cells in the brain parenchyma, along with immune cells in the dura, ChP, blood and spleen at 0.5 mo, 1 mo, and 3 mo. *P<0.05, **P<0.01, calculated by One-way ANOVA with Dunnett’s multiple comparisons test (C and E) and Two-way ANOVA with Tukey’s multiple comparisons test (G). Data are shown as the mean ±SEM.

We identified CD45^hi^, non-microglia immune cells, lacking vascular labeling in the brain parenchyma, dura, and ChP (Figure 1B and C, and Figure S1A). The brain parenchyma of 3-month-old adult mice (3 mo) showed a similar number of CD45^hi^ immune cells compared to those in the dura (Parenchyma: 7692 ± 890.6 cells, Dura: 8429 ± 940.6 cells, mean ± S.E.), while the ChP had a lower number of immune cells (ChP: 509.9 ± 52.21 cells). We didn’t observe any differences between males and females in the number of total CD45^hi^ immune cells in the brain (Figure S1B). To ensure that the presence of CD45^hi^ immune cells in the brain is not due to any pathology, we analyzed the proportion of naive/effector cells among CD4^+^ and CD8^+^ T cells, as well as the proportion of NK cells in the spleen. The mice we used showed a high naive portion of CD4^+^ and CD8^+^ T cells in the spleen and low levels of NK cells, suggesting non-pathological conditions in the experimental mice (Figure S2B-D). Next, we conducted confocal imaging of the mouse brain to confirm the presence of CD45^hi^ immune cells in the brain parenchyma (Figure 1D and Figure S3). First, we co-stained anti-CD45 antibody (clone 30-F11) with anti-IBA1 to see whether anti-CD45 staining labels CD45^hi^ immune cells, not IBA1^+^ microglia. We observed segregated populations stained with anti-CD45 (CD45^hi^ immune cells) and IBA1 (IBA1^+^ microglia) (Figure S3A). Next, using an anti-CD31 antibody to label blood vessels, we confirmed that CD45^hi^ immune cells are present in the brain parenchyma and are localized away from the vessels, including in the cerebral cortex, hypothalamus, and thalamus (Figure 1D). Next, we investigated when CD45^hi^ immune cells first appeared in the brain. Immune cells from the brain parenchyma were isolated at five different time points: postnatal day 1 (P1), 2 weeks (0.5 month; mo), 1 mo, 2 mo, and 3 mo. Surprisingly, CD45^hi^ immune cells were detectable as early as P1, with a trend of increasing numbers starting from 0.5 mo (Figure 1E). Additionally, we examined where immune cells localize within specific brain regions. The brain was divided into eight subregions: Olfactory Bulb (OB), Cortex (Cbx), Hippocampus (Hippo), Hypothalamus (Hypo), Striatum/Thalamus (S/T), Midbrain (MB), Hindbrain (HB), and Cerebellum (Cbll), with dura immune cells included as a control (Figure 1F and Figure S3B and C). Using flow cytometry, we quantified CD45^hi^ immune cells in each subregion, and the relative distribution was calculated as the number of CD45^hi^ immune cells in each region of brain divided by the total number of CD45^hi^ immune cells in whole brain (Figure 1G). Relatively, CD45^hi^ immune cells appeared more in OB, Cbx, Cbll, and HB (Figure 1G). We also confirmed their presence in the brain parenchyma with immunohistochemistry, although this method may include some CD45^hi^ immune cells from the leptomeninges or vasculature (Figure S3B and C). However, we observed no regional specificity of total CD45^hi^ immune cell number within the brain at different developmental stages (0.5 mo, 1 mo, or 3 mo), with the exception of a slight reduction observed in the Hypo in aged mice (Figure 1G).

We then assessed the identity of CD45^hi^ immune cells. By utilizing a panel of antibodies designed to characterize immune cell composition, we identified nine subtypes of CD45^hi^ immune cells, which accounted for approximately 90-95 % of total CD45^hi^ immune cells in the brain. These included Dendritic cells (DC), Macrophage, Monocyte, Neutrophil, Natural killer cells (NK), B cells (B), CD4^+^ T cells (CD4^+^ T), CD8 ^+^ T cells (CD8^+^ T), γδT cells (γδT), Ly6C^+^ monocytes, ST2^+^ ILC2, ST2^-^CD117 (c-kit)^+^ILC precursors (Figure 1H). We confirmed that no male/female differences were observed in any major immune cell population at 3 mo (Figure S1C). We isolated CD45^hi^ immune cells from the dura, blood, and spleen at 0.5 mo, 1 mo, and 3 mo, and compared the proportion of CD45^hi^ immune cells in the brain parenchyma with circulating CD45^hi^ immune cells from brain samples, which were labeled with anti-CD45 antibody before perfusion. During early postnatal stages, B cells constituted the primary immune cell population in the brain (approximately 56.11% at 0.5 mo). However, with age, the proportion of B cells dramatically declined (R^2^=0.432) and was replaced by Macrophage, CD4^+^ T cells (R^2^=0.539), and CD8^+^ T cells (R^2^=0.672) (Figure 1H and Figure S4A). This transition was more pronounced when comparing CD45^hi^ cell populations in the brain parenchyma with those in the dura, blood, and spleen (Figure 1H and Figure S4). Additionally, a large number of macrophage and B cells were detected in the dura, while B cells were the major cell population in blood and spleen (Figure 1H and Figure S4).

As an independent method to identify and quantify CD45^hi^ immune cells in the brain, we employed single-cell RNA sequencing (scRNA-seq). CD45^hi^ immune cells from the brain parenchyma and blood of 3 mo male, as well as brain parenchyma of 3 mo female mice, underwent a droplet-based scRNA-seq system, and the results were analyzed with the Seurat package (Figure S5)^24^. After removing low-quality scRNA-seq cells, we obtain 11,028 male-blood, 5,417 male-brain, and 3,138 female-brain CD45^hi^immune cells from 5 pooled samples. By using specific cell markers, we classified the nine immune cell groups; B cells (*Ms4a1*, *Cd79a*), CD4^+^ T cells (*Cd3e*, *Cd4*), CD8^+^ T cells (*Cd3e*, *Cd8a*), γδT cells (*Trdc*, *Trdv4*), NK cells (*Ncr1*, *Klrb1c*), Neutrophils (*S100a8*, *S100a9*), DCs (*Itgax*, *Mrc1*), and Macrophages/Monocyte (*Itgam*, *Fn1*) (Figure S5A, B, F, and G)^25^. Consistent with FACS analysis, CD8^+^ T cells constituted the major cell population in the brain, whereas B cells predominated in the blood by scRNA-seq (Figure S5C), while male and female brain samples didn’t show differences in cell proportions (Figure S5H). Moreover, the proportions and transcriptional profiles of each immune cell differed between the blood and brain, with effector-like genes (e.g. *Cd44* for T cells and *Cd86* for myeloid Cells) enriched in brain immune cells (Figure S5D and E)^26–28^. Consistently, we observed marginal differences on transcriptional profiles between male and female brain immune cells (Figure S5I and J). In summary, our findings suggest that the brain parenchyma has its own immune system, akin to other non-lymphoid organs, and can be distinguished from the dura and choroid plexus. Furthermore, our data highlight the increased abundance of T lymphocytes, particularly CD8^+^ T cells, within the brain parenchyma with age.

### Brain-specific CD8^+^ T cells migrate into the brain in a Naive state in early life

Despite the absence of prior infection, one possible explanation for the presence of CD45^hi^ immune cells in non-lymphoid organs like the brain parenchyma and dura is continuous immune surveillance, which is necessary for CNS integrity^29^. However, while CD8^+^ T cells are traditionally known as the immune system’s effector cells that “kill” virus-infected cells or tumor cells^30^, their role in the brain under non-pathological conditions, particularly whether it is restricted to immune surveillance, remains underexplored. We aimed to investigate unexplored functions of brain-specific CD8^+^ T cells and their unique characteristics in non-pathological conditions. We observed an age-dependent increase in both the absolute number and proportion of CD8^+^ T cells within total CD45^hi^ immune cells in the brain, contrasting with their levels in the blood (Figure 2A and B, and Figure S6A). Confocal imaging revealed scattered CD8^+^ T cells throughout the brain parenchyma, located away from CD31-positive blood vessels (Figure 2C). Notably, the brain parenchyma harbors significantly more CD8^+^ T cells compared to ChP and dura, suggesting potential active participation in brain functions followed by clonal expansion between 1 mo and 3 mo (Figure 2D). This expansion was more dramatic when we calculated the CD4/CD8 ratio in the brain, dura, blood, and spleen (Figure S6B), and linear regression between cell numbers and age (Figure S6C). In the peripheral immune system, CD4^+^ T cells are typically more abundant than CD8^+^T cells, resulting in a CD4/CD8 ratio greater than 1, typically around 2^31^. Consistent with this, we were able to observe approximately a two-fold difference between the number of CD4^+^ T cells and CD8^+^ T cells in blood and spleen at 0.5 mo, 1 mo, and 2 mo. Interestingly, the CD4/CD8 ratio in the brain is around 2.3 at 0.5 mo and dramatically decreases with age (1.09 at 1 mo and 0.5 at 2 mo) (R^2^=0.7429), while a high CD4/CD8 ratio (more than 1) is maintained in the dura (4.05 at 0.5 mo, 2.22 at 1 mo, and 1.99 at 2 mo) and ChP (2.2 at 0.5 mo, 2.3 at 1 mo, and 1.07 at 2 mo) (Figure S6B). A significant linear regression was observed between the logarithm of CD8^+^ T cell numbers in the brain and dura versus the logarithm of age (p-value < 2.2^e-16^, adjusted R^2^= 0.7013). The exponents were significantly different, with a value of 1.85 for the brain and 1.02 for the dura (Figure S6C). Except OB and Cbll, we observed that CD8^+^ T cells are globally distributed throughout the entire brain at 0.5 mo and that their numbers increase uniformly across almost entire brain regions with age, which indicate that age-dependent expansion of CD8^+^ T cells in the brain is not due to any localized infections (Figure S6D).

**Figure 2.**
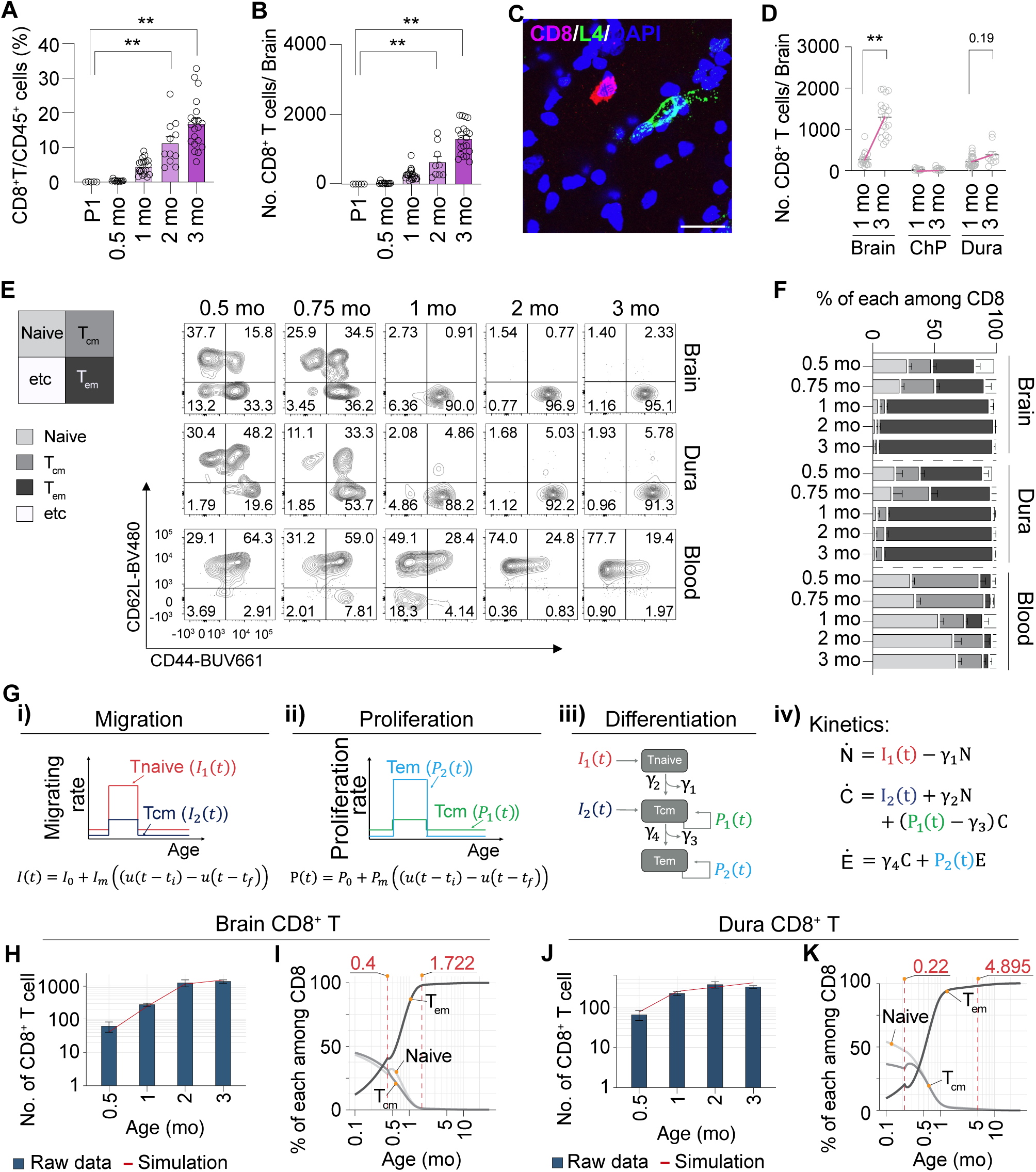
Brain-specific CD8^+^ T cells migrate into the brain in a Naive state during early life. (**A** and **B**) Relative and absolute distribution of CD8^+^ T cells in the brain at different ages (P1, 0.5 mo, 1 mo, 2 mo, and 3 mo), as analysed by flow cytometry. (**C**) Representative images showing CD8^+^ T cells located beyond the vessels (stained with CD31) in the brain. (**D**) Changes in the absolute number of CD8^+^ T cells in the brain, ChP, and dura from 1 mo to 3 mo, as analysed by flow cytometry. (**E** and **F**) Representative FACS plots of Naive, Tcm, and Tem CD8^+^ T cells (**E**), and their quantifications (**F**). The composition of CD8^+^ T cells in the brain, dura, and blood was analysed using flow cytometry. All FACS plots were gated on live-dead^-^, CD45^hi^, TCRβ^+^, CD4^-^, and CD8^+^ cells. (**G**) Formula showing an ordinary differential equation model (iv) encapsulating the processes of migration (i), proliferation (ii), and differentiation (iii) of brain- and dura- CD8^+^ T cells. (**H**) Number of brain- CD8^+^ T cells across different mouse ages. Bar plots represent real data, the red line represents the model’s solution. (**I**) Subtype proportions of brain CD8^+^ T cells across different mouse ages, as calculated from the model. The dotted red vertical lines indicate: (left) the onset of rapid migration and proliferation, and (right) when the proportion of Tem cells exceeds 98% of the total CD8^+^ T cell population. (**J**) Number of dura CD8^+^ T cells across different mouse ages. Bar plots represent real data, the red line represents the model’s solution. (**K**) Subtype proportions of dura CD8^+^ T cells across different mouse ages, as calculated from the model. The dotted red vertical lines indicate: (left) the onset of rapid migration and proliferation, and (right) when the proportion of Tem cells exceeds 98% of the total CD8^+^ T cell population. *P<0.05, **P<0.01, calculated by an One-way ANOVA with Dunnett’s multiple comparisons test (A and B), and Two-way ANOVA with Sidak’s multiple comparisons test (D). Data are shown as the mean ±SEM.

In the context of the conventional immune response, effective immune reaction progression encompasses several phases, including (i) activation, clonal expansion, and differentiation from naive to effector cells, (ii) migration to relevant tissues, cytokines synthesis, and clearance of target cells. To understand the phases of immune response, we employed flow cytometry to characterize T cell subpopulations using CD44 and CD62L markers at five different time points, including 0.5 mo, 0.75 mo. 1 mo, 2 mo, and 3 mo. CD62L^high^/CD44^low^ indicates naive CD8^+^ T cells (Tnaive), CD62L^high^/CD44^high^ indicates central memory CD8^+^ T cells (Tcm), and CD62L^low^/CD44^high^ means effector memory CD8^+^ T cells (Tem), respectively (Figure 2E). While blood CD8^+^ T cells remain as non-effector cells, surprisingly, CD8^+^ T cells in both brain parenchyma and dura are mostly Tnaive at 0.5 mo of age, shifting to an effector memory-like phenotype post-weaning, with mild differences on their differentiation kinetics (Figure 2E and F, Figure S6E). These results suggest that CD8^+^ T cells enter the brain as naive cells and subsequently differentiate within the brain. To rule out the possibility of blood contamination or mis-gated cells in our standard strategy, we first gated brain CD45^hi^ cells. After gating TCRβ^+^ and CD8^+^ T cells, we separated CD8^+^ T cells from the blood based on circulating CD45 labeling (Figure S6F). As expected, the new gating strategy (#2) did not alter the calculation of the proportion of naïve CD8^+^ T cells in the 0.5-month sample, indicating that these naive cells purely reside in the brain parenchyma (Figure S6G).

The next question is why we are able to detect naive cells only before weaning, not after weaning. Does this indicate that CD8^+^ T cell migration requires a critical time window? To explore this, we utilized T cell transplantation experiments. CD8^+^ T cells were isolated from spleen and lymph nodes from CD45.1 donor mice and cells were transferred via tail vein injection into 0.5 mo or 3 mo-old-recipient mice (Figure S6H). Interestingly, only the T cells transferred into 0.5-mo-old CD8 KO mice showed repopulation of CD8^+^ T cells in the brains (Figure S6I and J). In contrast, we did not observe high repopulation of CD8^+^ T cells in the dura when cells were transferred to 0.5-0.75-month-old mice, indicating that the migration kinetics of CD8^+^ T cells differ between the brain parenchyma and the dura (Figure S6I). Additionally, cells transferredd into 3 mo-old recipient mice did not repopulate not only the dura but also the brain efficiently (Figure S6J). These data suggest that existence of a specific developmental window that facilitates CD8^+^ T cell migration into the brain. However, how can we explain the rapid increase in CD8^+^ T cells from 1 month to 3 months (Figure 2D)? We next tested whether these cells were proliferating within the brain by analyzing the proportion of Ki67^+^CD8^+^ T cells. As we expected, CD8^+^ T cells from 0.5 mo-old and 1 mo-old mice showed an increased proportion of Ki67^+^ cells, whereas those from 3 mo-old mice did not (Figure S6K). These data suggest that brain CD8^+^ T cells enter the brain early as Tnaive or Tcm and proliferate within the brain, while cells in the dura exhibit slightly different kinetics. Based on these results, we formulated the ordinary differential equations (ODEs) model that encapsulates the processes of migration, differentiation, and proliferation of CD8^+^ T cells in the brain and dura (Figure 2G, Figure S6L and M). We denoted N for naive, C for central memory (Tcm), and E for effector memory (Tem) CD8^+^ T cells in the formula. To describe the steep increase in CD8^+^ T cells in 1 mo-old mice, we modeled the proliferation and migration of CD8^+^ T cells using an impulse function (Figure 2G, Figure S6L and M). We assumed that the differentiation rates from Tnaive to Tcm, and from Tcm to Tem are time-invariant. Parameters of the model were identified through a genetic algorithm with a fitness function that minimizes the difference between the model solution and the number of the CD8^+^ T cells based on FACS. The model proposed that the rapid proliferation of the memory CD8^+^ T cells and increased migration of naive CD8^+^ T cells, occurring before weaning, are essential to explain the explosive expansion of the Tem CD8^+^ T cell population in the brain and dura (Figure 2H-K). Expansion of the CD8^+^ T cells mainly came from the proliferation rate of Tem cells, which increased 14.4-fold in the brain and 22.1-fold in the dura during a rapid expansion period, suggesting the clonal expansion of CD8^+^ T cells. The number of brain-CD8^+^ T cells increases more rapidly than that of dura CD8^+^ T cells, and the proportion of Tem cells also rises quickly within a short period. In the dura, Tem cells make up more than 98% of total CD8^+^ T cells by about 5 mo, whereas in the brain, this threshold is reached at approximately 1.7 mo (right red dotted line in Figure 2I and K). The model indicates an earlier and shorter duration of the impulse function for migration and proliferation of CD8^+^ T cells in the brain, compared to cells in the dura, with the brain’s period spanning from 0.4 to 1.4 mo, and the dura from 0.22 to 0.82 mo (left red dotted line in Figure 2I and K). These results suggest that CD8^+^ T cells enter the brain in early life, specifically before weaning as naïve cells and becoming Tem CD8^+^ T cells, with kinetics different from those of cells in the dura. Then, the following question is how brain-specific CD8^+^ T cells acquire TCR signal to differentiate into Tem CD8^+^ T cells in the brain.

### CD8^+^ T cells differentiate into Tem in the brain through interaction with microglia

T cell differentiation from naive to effector is initiated by T-cell receptor (TCR) recognition and binding to cognate antigens presented on major histocompatibility complex (MHC) class I molecules by Antigen Presenting Cells (APCs). Beta-2 macroglobulin (β-2m) is a key compartment of MHC I that is essential for antigen presentation (Figure 3A)^32,33^. To investigate the mechanisms underlying the differentiate into Tem cells during later developmental stages, we explored whether clonal expansion occurs within this population in adulthood. Clonal expansion, characterized by the proliferation of T cells with identical TCR configurations upon antigen encounter, is a pivotal process in T cell activation^34^. Consistent with FACS analysis (Figure 2E and F), brain samples from 2 mo-old mice showed clonal expansion, evidenced by increased clonotype counts in rare clonal proportions, while blood from the same mice and samples from 0.5 mo-old mice didn’t (Figure 3B). Importantly, although we didn’t observe any clonal expansion in the 0.5 mo brain, they still showed decreased TCR diversity compared to blood CD8^+^ T cells, indicating that CD8^+^ T cell migration into the brain still requires TCR specificity. (Figure 3C). We further compared our TCR repertoire to the public database, VDJDB, which stores several pathogen-related repertoires from various studies^35^. The results showed the most abundant TCR sequences from blood and brain samples at 0.5 mo and 2 mo did not overlap with those observed in disease, such as influenza infection from the analysis of the repertoire similarity (Infection Similarity Index) using the Jaccard index (Figure S7A). We also compared TCR diversity of CD8^+^ T cells with other tissues, including the mesenteric lymph node (mLN), lung, liver, and large intestine (L.I.). As expected, brain CD8^+^ T cells exhibited the highest degree of clonally expansion (Figure S7B).

**Figure 3.**
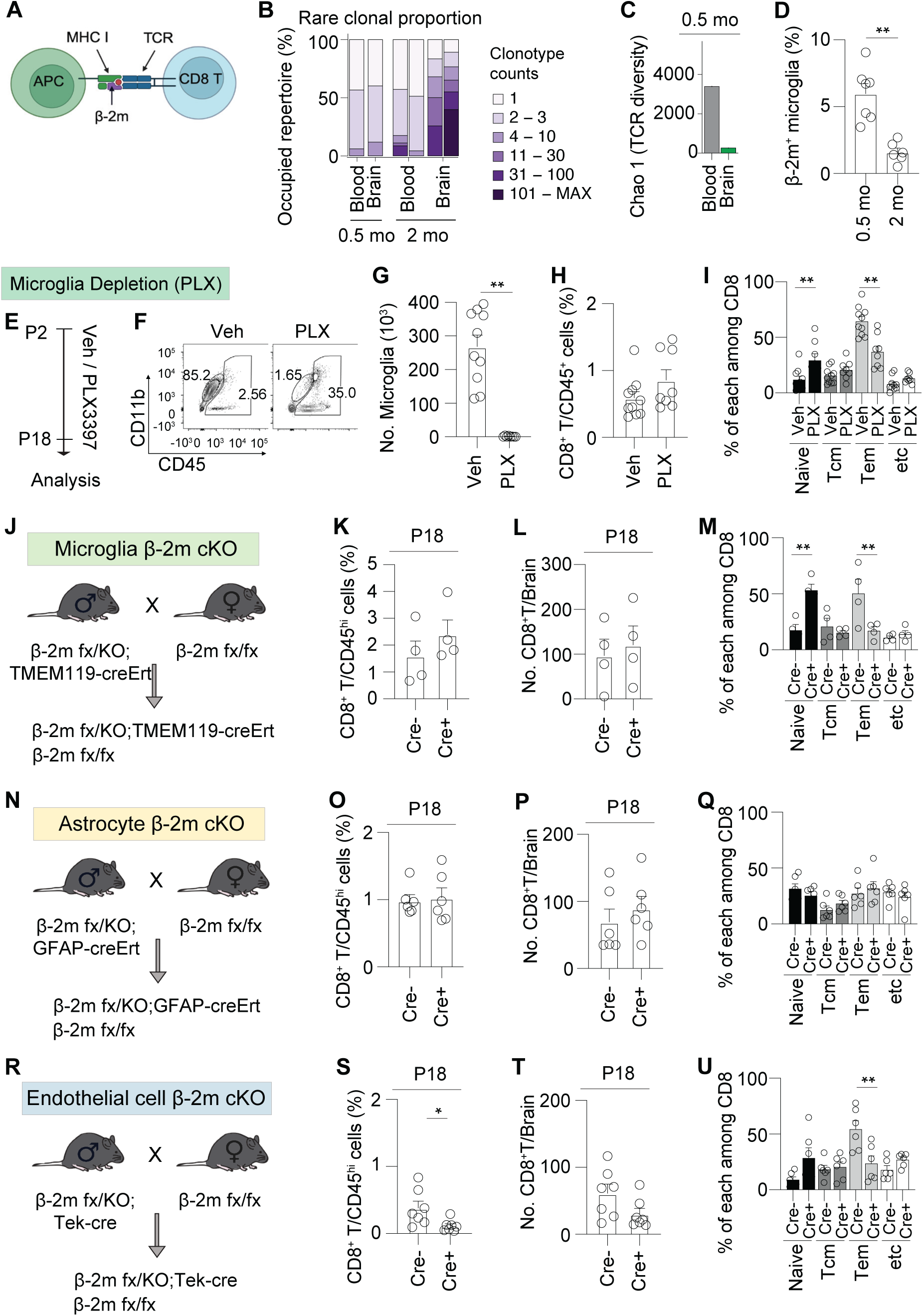
CD8^+^ T cells acquire an effector-memory characteristic in the brain through interaction with microglia. (**A**) Schematic diagram showing canonical interaction between antigen-presenting cells and CD8^+^ T cells. APC; antigen-presenting cells, MHC I; TCR; T-cell receptor. (**B**) The proportion of rare clonal CD8^+^ T cells was determined from TCR sequencing results. Blood and brain CD8^+^ T cells were isolated from 0.5 mo and 2 mo old male mice (0.5 mo, n=1, pooled with 10 samples, from 1 independent experiment; 2 mo, n=2, pooled with 10 samples, from 2 independent experiments). (**C**) TCR diversity was analysed using bulk TCR sequencing. Blood and brain CD8^+^ T cells were isolated from 10 mice at 0.5 mo of age and pooled for sequencing (n=1, pooled with 10 samples, from 1 independent experiment). (**D**) The proportion of β-2m^+^ microglia over total microglia at 0.5 mo and 2 mo mouse brains, as analysed by flow cytometry. (**E**) A schematic diagram of PLX injection strategy. Microglia cells in the brain were targeted by subcutaneously injecting 30 mg/kg of PLX (PLX3397, CSF1R inhibitor) from postnatal day 2 to 21st. (**F**) Microglia depletion of vehicle (Veh) or PLX-injected (PLX) mice was analysed in the brain-resident immune cells using flow cytometry, and representative FACS plots were shown. (**G**) The number of microglia in the brain of each mouse was counted based on FACS recording of whole-brain CD45^hi^ immune cells. (**H**) Relative distribution of CD8^+^ T cells among CD45^hi^ immune cells in the brain with Veh and PLX. (**I and J**) Quantifications to show the composition of CD8^+^ T cells in the brain with Veh and PLX (n=8, from 3 independent experiments). (**K**) Mating strategy to generate TMEM119-creERT2 dependent β2m conditional knock-out mice. (**L** and **M**) Relative distribution of CD8^+^ T cells among CD45^hi^ immune cells in the brain of WT and microglia-specific β2m cKO. (**N and O**) Quantifications to show the composition of CD8^+^ T cells in the brain of WT and microglia-specific β2m cKO (n=4, from 4 independent experiments). (**P**) Mating strategy to generate GFAP-creERT2 dependent β2m conditional knock-out mice. (**Q** and **R**) Relative distribution of CD8^+^ T cells among CD45^hi^ cells in the brain of WT and astrocyte-specific β2m cKO. (**S and T**) Quantifications to show the composition of CD8^+^ T cells in the brain of WT and astrocyte-specific β2m cKO (n=6, from 3 independent experiments). (**U**) Mating strategy to generate Tek-cre (Tie2 cre) dependent β2m conditional knock-out mice. (**V and W)** Relative distribution of CD8^+^ T cells among CD45^hi^ cells in the brain of WT and brain endothelial cell-specific β2m cKO (**X and Y**) Quantifications to show the composition of CD8^+^ T cells in the brain of WT and brain endothelial cell-specific β2m cKO (n=6, from 4 independent experiments). **P<0.01, calculated by student t-test (E, H, I, L, and O), and Two-way ANOVA with Sidak’s multiple comparisons tests (J, M, and P). Data are shown as the mean ±SEM.

Previously the potential role as APCs of microglia was suggested due to their expression of MHC I and II, as well as co-stimulatory molecules necessary for T cell activation. We found that *H2-D1*, an MHC I gene in mice, was upregulated in microglia, oligodendrocyte, and endothelial cells before weaning in a healthy brain (Figure S7C)^36, 37^. Therefore, we planned to remove β-2m from each cell population to see which cell populations act as APC to naive brain-CD8^+^ T cells. We confirmed increased β-2m^+^ microglia prior to weaning (0.5 mo) compared to β-2m^+^ cells from 2 mo-old mice in its protein level (Figure 3D). Then, we pursued pharmacological microglia depletion experiments using PLX-3397, initiated in mice at postnatal day (P) 3 for 2 weeks. Consistent with previous findings, PLX-3397-mediated microglia depletion resulted in arrested differentiation of CD8^+^ T cells within the brain (Figure 3E-I). However, PLX-3397-mediated microglia depletion also generates effects on CD8^+^ T cells in the spleen (Figure S7D-H). It is also important to acknowledge that PLX-3397 not only inhibits CSF1R but also targets c-Kit. This dual activity could contribute to off-target effects that extend beyond microglial depletion^38^. Furthermore, emerging evidence suggests that newer, more selective analogs of PLX-3397 can modulate brain endothelial cholesterol metabolism independently of microglia^39^. These findings underscore the need for careful interpretation of results derived from PLX-3397-based experiments and highlight the evolving complexity of pharmacological tools used to probe microglial function. Therefore, we generated microglia-specific MHC I conditional KO (cKO) by crossing β-2m floxed mice with TMEM119-CreERT2 (Figure 3J). Tamoxifen was injected into nursing dams 5 days after delivery, and it was delivered by breast milk to pups. Littermates with no cre (Cre-) were used as control samples. As we expected, depletion of β-2m on microglia (Cre+) led to an arrest in the differentiation of CD8^+^ T cells within the brain (Figure 3K-M, and Figure S7I-K), while generating no effects on peripheral CD8^+^ T cells (Figure S7L and M). This collective evidence implicates microglia as acting as APC, triggering the differentiation of CD8^+^ T cells, respectively. Since other cell populations in the brain, including astrocyte and endothelial cell also expressed increased β-2m before weaning, we attained through the generation of astrocyte-specific, and endothelial cell-specific MHC I conditional KO to show whether astrocytes, oligodendrocyte, and endothelial potentially can act as APC (Figure 3N-U and Figure S7N-W). Interestingly, astrocyte--specific cKO didn’t affect the proportion of CD8^+^ T cells in the brain and spleen, while endothelial cell-specific cKO showed the effect on Tem CD8^+^ T cell proportions in the brain and also naive and effector memory cells in the spleen, which might be important for the global Tem cell migration. Collectively, our findings implicate microglia as key APCs triggering the differentiation of CD8^+^ T cells within the brain microenvironment. Taken together, we propose that migrated naive CD8^+^ T cells into the brain undergo transformation into Tem CD8^+^ T cells through interactions with microglia.

### Microbiome-dependent cues during weaning promote CD8^+^ T cell expansion

One caveat from our APC experiments (Figure 3) is that, while we were able to identify the APCs responsible for inducing the transformation of CD8^+^ T cells from a naive to an Tem state, we did not observe any reduction in CD8^+^ T cell numbers in the brains of cKO mice. This finding suggests that two distinct mechanisms may underlie CD8^+^ T cell differentiation and clonal expansion. A recent publication has highlighted the interaction between the gut microbiome and T cells in the brain^10^. This study demonstrated that gut-licensed T cells undergo clonal expansion in the brain after weaning. During birth, various body surfaces are colonized by microbes, which play a crucial role in shaping immune system development. Notably the weaning-associated response to microbiota is essential for immune ontogeny^40,41^. Based on these findings, we focused on investigating whether changes in the microbiota of pups influence immune cell entry into the brain. We conducted a comparative analysis of mice housed under specific pathogen-free (SPF) and germ-free (GF) conditions. Intriguingly, mice maintained in the GF environment exhibited no significant changes in their CD8^+^ T cell proportions before and after weaning (Figure 4A), whereas cells in the dura showed a less robust but still meaningful increase (Figure 4B). Interestingly, a similar but weaker pattern was observed for CD4^+^ T cells (Figure 4C and D).

**Figure 4.**
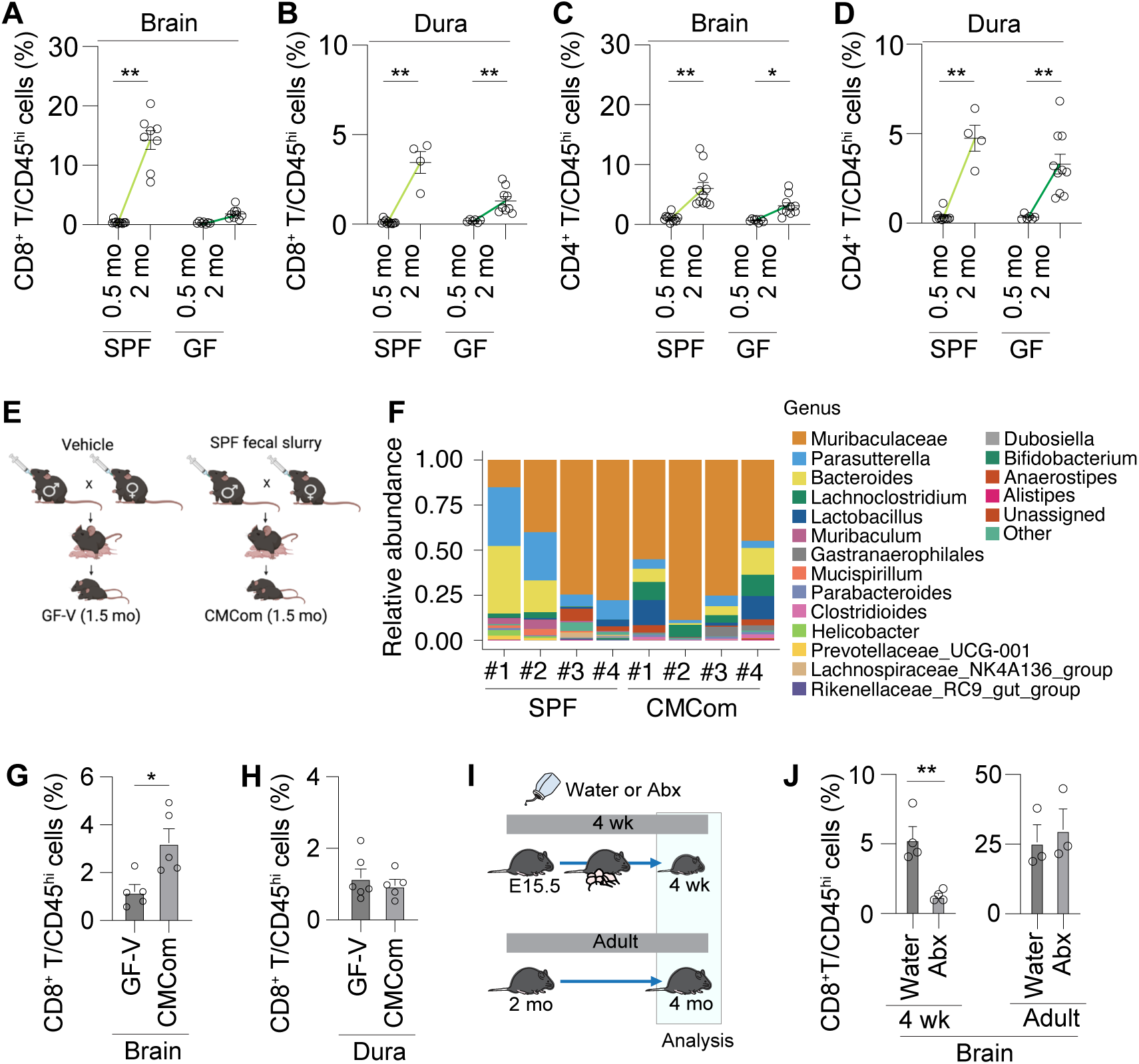
Microbiome-dependent cues during weaning promote CD8^+^ T cell expansion. (**A-D)** Changes in the relative distribution of CD8^+^ T cells and the CD4^+^ T cells among CD45^hi^ cells in the brain, and dura from 0.5 mo to 2 mo. (**E**) Schematic view of making gnotobiotic mouse model. Germ-free mouse was gavaged fecal slurry collected from SPF mice. (**F** and **G**) Relative distribution of CD8^+^ T cells among CD45^hi^ immune cells in bran and dura of Germ-free mice (GF-V) and CMCom gnotobiotic mice (CMCom) (n=5, from 2 independent experiments). (**H**) Schematic view of antibiotics treatment strategy. The mix of antibiotics or water was administered through drinking water from E15.5 (through mom) to 4 weeks of their age (4wk) or from 2 mo to 4 mo (Adult) of their age (**I**) Relative distribution of CD8^+^ T cells among CD45^hi^ immune cells in the brain.

Next, we colonized GF mice with SPF fecal slurry (CMCom) and confirmed that introducing the SPF gut microbiome was sufficient to increase CD8^+^ T cell numbers in the brain after weaning, but not in the dura (Figure 4E-G). We then investigated whether the weaning period represents a unique time window for CD8^+^ T cell clonal expansion, similar to their migration into the brain. To test this, we depleted the gut microbiome using antibiotic cocktails either before weaning or after two months of age (Figure 4H). The 4-week group received antibiotics from embryonic day 15.5 (E15.5) until one month of age (right after weaning), while the adult group received antibiotics from two months of age for two months. Each group was examined at the end of their respective antibiotic treatments (Figure 4H). Interestingly, we observed that the presence of the microbiome was required to induce CD8^+^ T cell expansion during the weaning period (Figure 4I). In summary, our findings reveal that CD8^+^ T cell expansion depends on microbiome-derived signals during the weaning stage.

### The absence of brain-specific CD8^+^ T cells is associated with abnormal behaviors through altering neuronal activities

To investigate the role of CD8^+^ T cells in the brain, we used CD8 knockout (KO) mice, which lack functional cytotoxic T cells but have no defects in CD4 helper T cells. CD8 KO mice were confirmed to have no CD8^+^ T cells and no difference in the proportion of CD4^+^ T cells in the brain, although there was a small increase in CD4^+^ T cells in the spleen (Figure S8A-E). Next, we conducted a series of behavioral tests to assess developmental milestones, including eye opening, Tactile startle reflex, grasp reflex, cliff avoidance, negative geotaxis, and surface righting (Figure S8A-K). The absence of CD8 didn’t affect the mouse’s developmental milestones compared to age-matched WT mice (Figure S8A-K). We then tested whether the absence of CD8^+^ T cells has any impact on adult behaviors related to social interaction, emotion, and cognition using a battery of behavior tests, including assessments of social behavior, anxiety, and recognition in adulthood. Anxiety was tested with two different assays; open field test (OFT) and elevated zero maze test (EZM). Each test allows us to analyze the level of anxiety by measuring the time spent in the center zone (OFT) or open arm (EZM). Surprisingly, CD8 KO mice displayed decreased time spent in the center zone (OFT) and open arm (EZM), indicating increased anxiety, with slightly decreased velocity in the OFT (Figure S8L and M). Sociability was tested using the three-chamber social interaction test. While CD8 KO mice didn’t display abnormal interaction time with an empty cup or a cup with a social target mouse, nor did they show differences in total distance moved, they exhibited decreased sociability, indicating deficits in social interaction (Figure S8N). Additionally, these mice showed impaired novel object recognition (Figure S8O). In parallel, we examined whether microglia-specific MHC I cKO (Figure 3K) mice and 4 wk Abx-treated mice (Figure 4H) also exhibited behavioral deficits. Consistent with our CD8^+^ T cell finding (Figure 3 and 4), both groups of mice showed significantly reduced social interaction and center time (Figure S8P and Q).

Next, we sought to determine whether a specific time window is critical for CD8^+^ T cells to be involved in maintaining brain homeostasis. Given the substantial expansion of CD8^+^ T cells during juvenile stages, we investigated whether the presence of CD8^+^ T cells in the brain during this period is necessary for normal behavior. To selectively control CD8^+^ T cell numbers during a specific time window, we used a blocking antibody to deplete CD8^+^ T cells at two different time points, starting at postnatal day (P) 21 (3 weeks) and 9 wk. P21 pups or 9 wk-old mice were treated with an anti-mouse CD8a antibody (Clone #2.43, BioXCells) or an IgG2b isotype control via intraperitoneal injection. The first three injections were administered over three days, followed by injections every five days until the week of behavior tests. At 8 or 13 wk old, mice underwent social interaction and anxiety tests, alongside FACS analysis to confirm T cell depletion efficiency after the behavior tests (Figure 5A and Figure S9Q).

**Figure 5.**
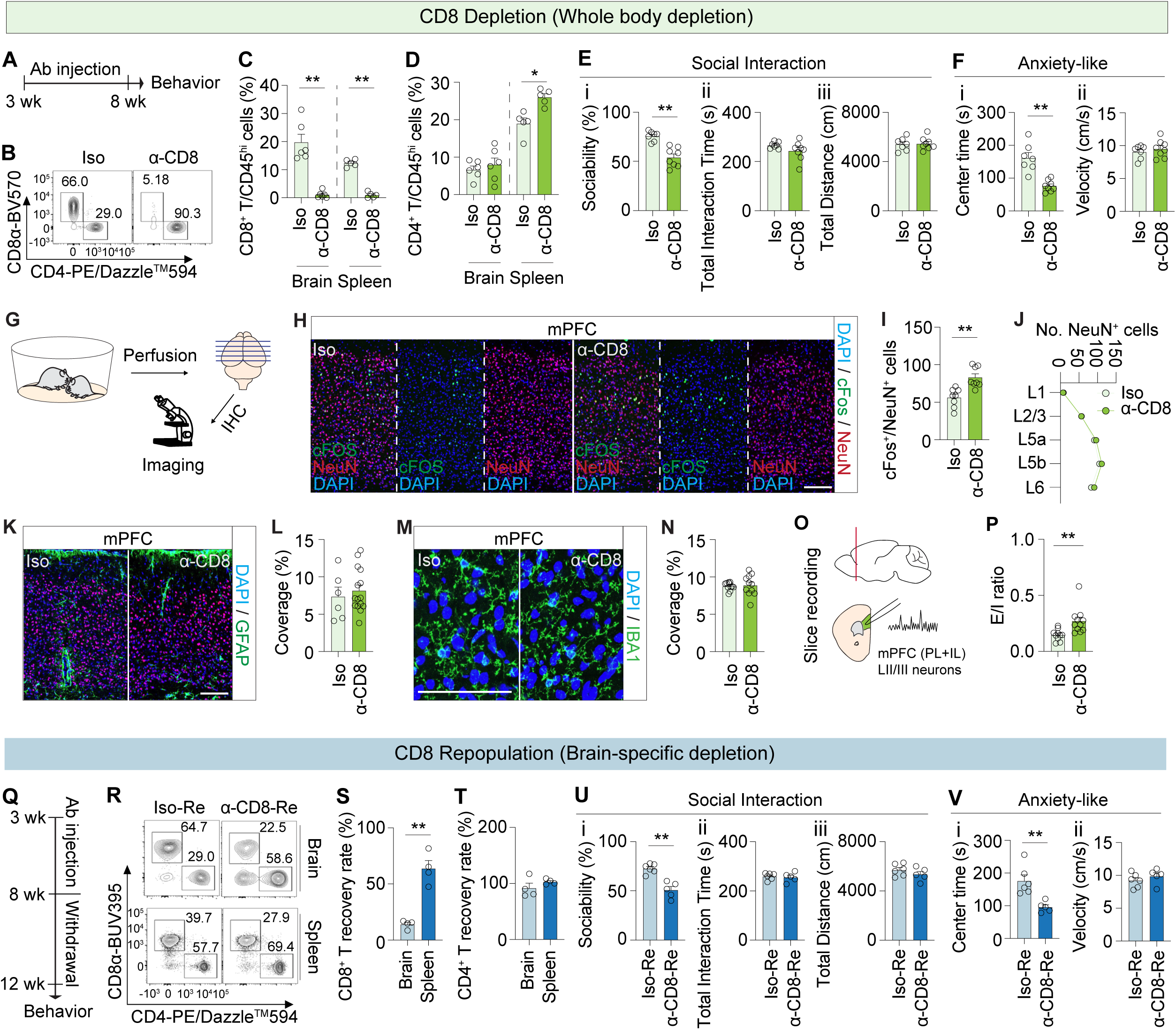
The absence of brain-specific CD8^+^ T cells is associated with abnormal behaviors through altering neuronal activities. (**A**) A CD8 depletion model was generated by i.p injecting an isotype control antibody or anti-CD8 antibody(clone 2.43) to WT C57BL/6 mice from 3 wks to 8 wks according to the injection protocol every 3-5 days. Social interaction and anxiety were tested at 8 wks (Iso and α-CD8). (**B**-**D**) The composition of brain and spleen CD4^+^ T and CD8^+^ T cells in both male mice groups, which were injected with isotype control antibody (Iso) or anti-CD8 depleting antibody (α-CD8), was analysed by flow cytometry. Clone 53-6.7 CD8a-detecting FACS antibody was used to avoid epitope competition with anti-CD8 depleting antibody (clone 2.43). Representative FACS plots of T cells (Livedead^-^, CD45^hi^TCRβ^+^) (**B**) were shown. Proportions of CD8^+^ T cells (**C**) and CD4^+^ T cells (**D**) in the CD8-depleted mice brain and spleen are shown (Iso n=6; α-CD8, n=6 from 3 independent experiments). (**E** and **F**) Social interaction (**E**) and anxiety-like (**F**) tests were conducted on Iso and α-CD8 mice (ISO, n=7, α-CD8, n=8, from 3 independent experiments). (**G**) Schematic view of experimental flow for measuring cFos expression in the brain. (**H**) Representative images illustrating cFos (green) and NeuN (red) expression in the mPFC of Iso (left) and α-CD8 (right). Scare bar: 200um. (**I and J**) Quantification of cFos^+^NeuN^+^ cells (I) and total NeuN^+^ cells per layer (J) in the mPFC (Iso, n=8, α-CD8, n=8, from 4 independent experiments). cFos^+^NeuN^+^ cells were counted within the rectangle ROI (415 μm width X 830 μm height). (**K**) Representative images illustrating GFAP^+^ astrocyte (green) in the mPFC of Iso (left) and α-CD8 (right). Scare bar: 200um. (**L**) The coverage of GFAP^+^ cells in the mPFC (Iso, n=6, α-CD8, n=15, from 3 independent experiments). (**M**) Representative images illustrating IBA^+^ microglia (green) in the mPFC of Iso (left) and α-CD8 (right). Scare bar: 200um. (**N**) Quantification of IBA1^+^ microglia coverage in the mPFC (Iso, n=11, α-CD8, n=11, from 3 independent experiments). (**O and P**) Schematics to show the recording area (mPFC layer II/III neurons) (**O**) and E/I ratio measured in L2/3 excitatory neurons of mPFC (**P**) (Iso, n=10 from 3 mice, α-CD8, n=11 from 3 mice, from 3 independent experiments). (**Q**) A CD8 repopulation model was generated by injecting depleting antibody for 5 wks from 3wks of mice age followed by 4 wks of antibody withdrawal (Iso-Re and α-CD8-Re). (**R**) Representative FACS plots of T cells (Livedead^-^, CD45^hi^TCRβ^+^) were shown. (**S and T**) The repopulation rate was calculated as the percentage of CD8^+^ T cells or CD4^+^ T cells in α- CD8-Re over Iso-Re in both the brain and spleen. (**U** and **V**) Social interaction (**U**) and anxiety-like (**V**) tests were conducted with Iso-Re and α-CD8-Re mice. *P<0.05, **P<0.01, calculated by a student t-test (C-F, I-N, P, and S-V). Data are shown as the mean ±SEM.

In the juvenile depletion group, mice treated with the IgG2b isotype control (Iso) had approximately 15% brain-resident CD8^+^ T cells among total brain CD45^hi^ cells, whereas treatment with the anti-mouse CD8a antibody (α-CD8) reduced this to less than 2% (Figure 5B and C). Similar depletion efficiency was observed in the spleen (Figure 5C). Conversely, other immune cells, including CD4^+^ T cell levels remained unchanged in the brains of ISO and α-CD8 mice, with a slight increase observed in the neutrophil of α-CD8 mice (Figure 5D and Figure S9A-G). The effective depletion of CD8^+^ T cells was also confirmed by their absolute numbers in the brain (Fig S9H). In the periphery, only CD4^+^ T cells showed a slight increase observed in the spleens of α-CD8 mice (Figure 5D and Figure S9A-G). Consistent with the behavioral deficits observed in CD8 KO mice, α-CD8 mice displayed abnormal sociability and increased anxiety, while their exploratory behaviors remained intact (Figure 5E and F). Next, we explored how the absence of CD8^+^ T cells in the brain induces abnormal behaviors. Using cFos immunoreactivity as a proxy for neural activity, we investigated whether neurons in specific brain areas are more susceptible to the absence of CD8^+^ T cells. After habituating the mice in the procedure room, they were sacrificed to measure neuronal activity via cFos expression in both Iso and α-CD8 mice. Using microscopy, we quantified cFos^+^ cell counts across multiple brain regions that have been previously implicated in social deficits and/or anxiety-related behaviors (Figure 5G). These regions included the medial prefrontal cortex (mPFC), basolateral amygdala (BLA), hippocampus, primary motor cortex (M1), primary somatosensory cortex (S1), and ectorhinal cortex (ECT) (Figure 5H and I, Figure S10A)^42,43^. Surprisingly, mice with reduced CD8^+^ T cell populations exhibited significantly increased cFos^+^ staining in the mPFC and a minor, though not statistically significant, increase in the BLA. Separate quantification of the cFos^+^ and cFos^-^ populations among NeuN^+^ neurons revealed a significant increase only in cFos^+^ cells in the α-CD8 group (Figure 5H and I, Figure S10B). However, we did not observe any group differences in the numbers of neurons, astrocytes, or microglia (Figure 5J-N). We further analyzed microglial reactivity in more detail. Using the function “Analyze skeleton” and the plugin “fractal analysis”, we assessed 10 different morphological characteristics of microglia: cell body size, number of branches, average brain length, number of junctions, length of the longest-shortest path, endpoints per microglia, circularity, fractal dimension, density, and span ratio (Figure S10C-L). Although overall IBA1^+^ microglial coverage (based on signal intensity) did not differ between groups (Fig 5M and N), subtle alterations in microglial morphology were detected (Figure S10C-L). Next, we isolated microglia from Iso and α-CD8 mice and performed bulk RNA sequencing to determine whether the absence of CD8^+^ T cells affects microglial transcriptional profiles (Figure S10M). While several genes were upregulated in α-CD8 mice, there were no specific enrichments associated with homeostatic (HOM), M1-, M2-, or disease-associated microglia (DAM) gene signatures (Figure S10N and O, Table S5). These histological examinations suggest that the absence of CD8^+^ T cells leads to increased neuronal excitability, particularly in the mPFC, with minimal effects on microglial morphology and function.

Dysregulation of neural activity in the mPFC has previously been associated with emotional processing, decision-making, memory, self-perception, and social behaviors in general^44^. To further investigate how CD8^+^ T cell depletion alters neuronal circuit function, we conducted brain slice recording to measure the neuronal Excitation/Inhibition ratio (E/I ratio) of neurons in the mPFC. Whole-cell patch-clamp recordings from pyramidal neurons in layer II/III revealed a significantly increased E/I ratio in the mPFC of CD8 depleted mice (Figure 5O and P), compared to controls, aligning with the elevated cFos expression (Figure 5H and I). This suggests heightened net excitatory drive within the mPFC in the absence of CD8^+^ T cells. Interestingly, despite the increased E/I ratio, the number of action potentials evoked by current injection was significantly reduced in α-CD8 mice, indicating reduced intrinsic excitability of mPFC neurons (Figure S10P). This was further supported by a decrease in input resistance (Ri), suggesting that the neuronal membrane properties were altered in a way that reduced responsiveness to depolarizing input (Figure S10Q). To determine whether changes in inhibitory synaptic architecture underlie these physiological alterations, we examined inhibitory synapse markers using immunostaining for vGAT (a presynaptic GABAergic marker) and gephyrin (a postsynaptic scaffolding protein). While the overall coverage of cGAT and gephyrin signals remained unchanged between groups, we observed a significant increase in the number of inhibitory synapses in α-CD8 mice, defined by co-localization of vGAT and gephyrin puncta (Figure S10R-U). Altogether, our findings indicate that CD8^+^ T cells regulate cortical circuit balance through dysregulation of inhibitory synaptic organization and neuronal excitability, rather than through a simple loss of synaptic elements.

To further determine whether the observed phenotypic consequences specifically stemmed from brain-resident CD8^+^ T cells rather than peripheral CD8^+^ T cells, we utilized a CD8^+^ T cell repopulation mouse model (α-CD8-Re). Following one month of antibody withdrawal (Figure 5Q), notable repopulation of CD8^+^ T cells occurred in the spleen, whereas the brain recovered to approximately 20% of typical CD8^+^ T cell numbers (Figure 5R-T and Figure S9P). We also confirmed this repopulation model didn’t affect other immune cells in the brain and spleen (Figure S9I-O). Importantly, even with this repopulation, persistent behavioral deficits were evident, demonstrating that the observed phenotypic alterations in adult behaviors, as seen in both CD8 KO and α-CD8 mice, are attributable to the absence of CD8^+^ T cells within the brain (Figure 5U and V). Furthermore, this consistency in phenotypic manifestations suggests that the presence of CD8^+^ T cells in the brain before adulthood (in a fully mature brain) is crucial for circuit formation.

Separately, we depleted CD8^+^ T cells in adult mice (starting at 9 wk old for 4 wk; α-CD8-AD) to investigate whether there is a critical time window during which CD8^+^ T cells influence brain function (Figure S9Q). Unfortunately, we achieved only around 50% depletion efficiency of CD8^+^ T cells in the brain, while the periphery (Spleen) showed almost complete depletion (Figure S9R and S). More interestingly, α-CD8-AD mice didn’t display any abnormal behaviors (Figure S9T and U). These results potentially indicate two things: 1) CD8^+^ T cells are required for normal behavior during a specific time-window, and 2) the absence of peripheral CD8^+^ T cells doesn’t impact these types of behaviors, respectively.

### Brain-specific CD8^+^ T cells show unique tissue-resident characteristics

To gain a deeper understanding of how brain-CD8^+^ T cells affect neuronal activity and behavior, we aimed to identify the characteristics of brain-CD8^+^ T cells. We integrated dura CD8^+^ T cells from public data with our CD8^+^ T cell cluster (Figure 6)^45^. This comparison revealed heterogeneity within CD8^+^ T cell lineage from blood, brain, and dura (3 mo and 20 mo) (Figure 6A-E). In more detail, while blood-CD8^+^ T cells show naive characteristics with high *Sell*, *Ccr7*, and *Tcf7*, brain and dura CD8^+^ T cells displayed higher expression levels of genes associated with effector functions, including *Ccl5*, *Cxcr3*, and *CD44*. Also, both brain and 3 mo dura (D-3m) CD8^+^ T cells showed increased tissue-resident markers, such as *Itga1*, and decreased circulating markers like *S1pr1* (Figure 6B and Figure S11A). However, surprisingly, while both brain and M-3m CD8^+^ T cells have low expression of cytotoxic molecules, *Gzma*, with a slight increase in *Gzmb* expression in brain CD8^+^ T cells, only brain CD8^+^ T cells showed increased *Tnfa* and *Ifng* (Figure 6B)^46–48^. To investigate the differential transcriptomic states of CD8^+^ T cells in the brain, blood, and dura, we utilized Monocle2 to identify and order cells along a pseudotime trajectory (Figure 6C and D)^49^. The pseudotime analysis revealed two distinct trajectories (from blood to brain and from blood to dura), underscoring the unique characteristics of CD8^+^ T cells in the brain compared to those in the blood and dura (Figure 6C). To pinpoint the genes involved in the progression of CD8^+^ T cells along each trajectory, we conducted hierarchical clustering of genes whose expression varied with pseudotime. Interestingly, by mapping gene expression along the bifurcated paths, we discovered that CD8^+^ T cells in the brain tended to upregulate genes associated with tissue residency and effector functions (Figure 6D). Moreover, gene ontology (GO) analysis revealed that genes associated with ‘Naive’ cells were upregulated as pseudotime progressed in blood. In contrast, genes related to ‘Effector or Memory CD8^+^ T cells’ were upregulated along both the brain’s and D-3m’s trajectories as pseudotime advanced (Figures 6D and Table S2-3). These findings support the idea that CD8^+^ T cell identity is organ specific. Cell identity is regulated by transcription factors (TFs) that control the expression of its target genes. The network of interacting TFs that govern gene expression, and thereby determine cell identity, is known as a Gene Regulatory Network (GRN). The target genes regulated by a specific TF are grouped into units called the regulons. To identify organ-specific regulons, we used SCENIC (single-cell regulatory network inference and clustering) (Figure 6E)^50,51^. This analysis allowed us to identify organ-specific regulons active in CD8^+^ T cells. While naive-related TFs, such as *Tcf7* and *Lef1*, were enriched in blood CD8^+^ T cells, effector-related TFs, such as Eomes and *Tbx21*, were more prominent in brain CD8^+^ T cells. Consistent with the pseudotime analysis (Figure 6C and D), dura CD8^+^ T cells also exhibited elevated levels of Tem markers (Table S4).

**Figure 6.**
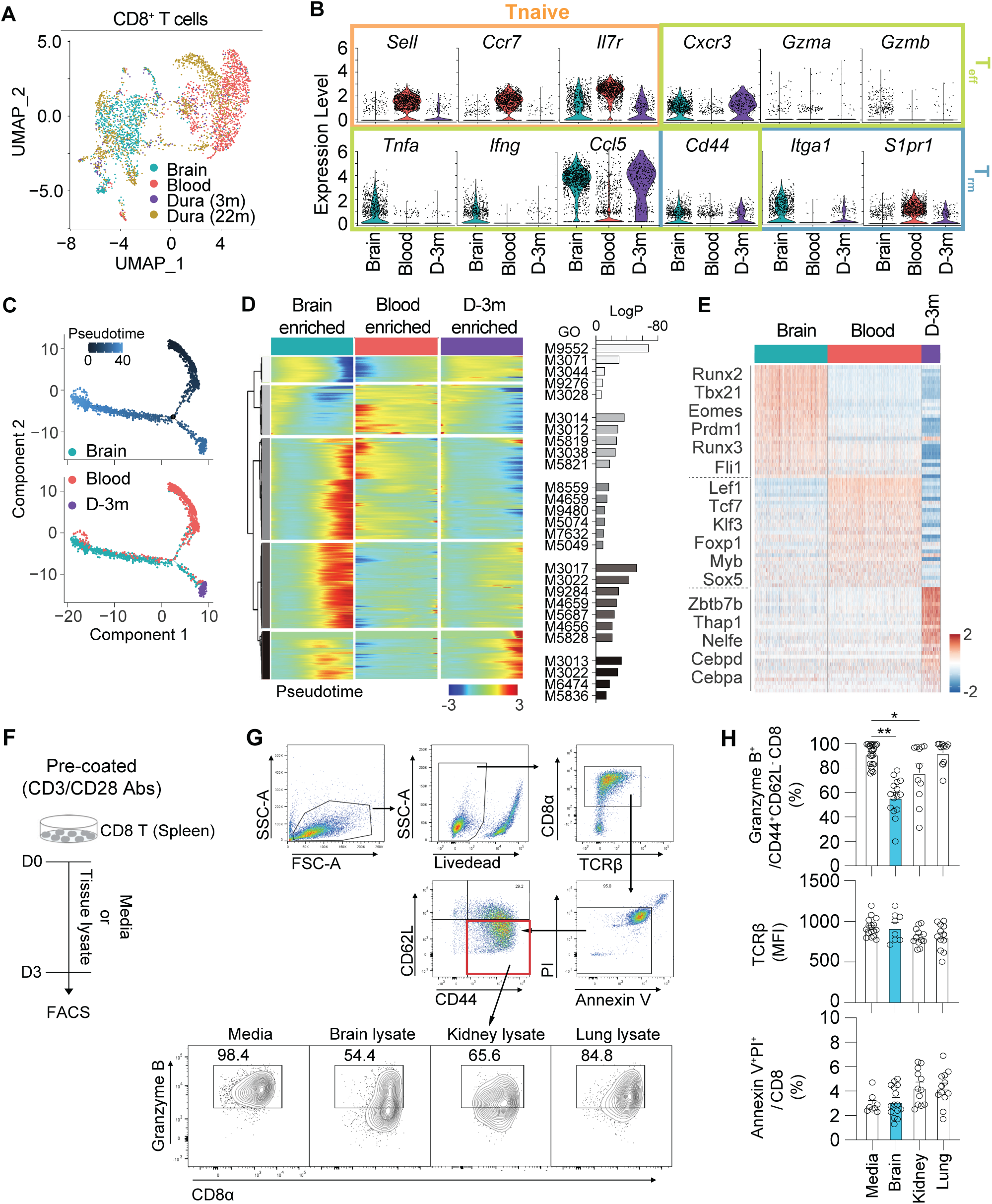
Brain-specific CD8^+^ T cells show unique tissue-resident characteristics. (**A**) UMAP embedding in CD8^+^ T cells for brain, blood, 3 mo dura (D-3m), and 22 mo dura (D-22m). Colors represent origin tissues. (**B**) Violin plots showing indicated marker gene expressions for Tnaive (*Sell, Ccr7, Il7r*), Teff (*Cxcr3, Gzma, Gzmb, Tnfa, Ifng, Ccl5, Cd44*), and Trm (*Cd44, Itga1, S1pr1*) from CD8^+^ T cells in the brain (blue), blood (red), D-3m (purple). (**C**) Pseudotime analysis of CD8^+^ T cells for brain, blood and D-3m. (**D**) Heatmap of differentially expressed genes in different sub-population of CD8^+^ T cells over pseudotime and selected GO terms to represent each cluster. Each column represents different origin tissues (brain, blood and D-3m). (**E**) z-scaled TF regulon enrichment score of the CD8^+^ T cells in brain, blood, and D-3m. Specific regulons are depicted. (**F**) Schematic diagram of CD8^+^ T -tissue lysate co-culture experiment. Isolated naive CD8^+^ T cells from the spleen and lymph nodes were seeded on the CD3ε/CD28 antibody pre-coated plate. Cells were incubated in the media only or tissue lysate for 3 days. (**G** and **H**) Representative FACS plots (**G**) and quantification graph (**H**) to show Granzyme B^+^ Tem CD8^+^ T cells, TCRb (MFI), and AnnexinV^+^PI^+^ cells from cells incubated with media or brain, kidney, or lung tissue lysate (Media, n=23, Brain, n=15, Kidney, n=11, Lung, n=12, from 5 independent experiments). *P<0.05, **P<0.01, calculated by an One-way ANOVA with Dunnett’s multiple comparisons test (H). Data are shown as the mean ±SEM.

As an independent approach to identify the characteristics of brain-CD8^+^ T cells, we used high dimensional flow cytometry. Specifically, to confirm the level of granzyme B protein, we isolated brain CD8^+^ T cells from naive mice and LCMV Amstrong-infected mice as a positive control and compared their granzyme B and perforin secretion by flow cytometry (Figure S11B-E)^52^. While CD8^+^ T cells from brain and spleen of LCMV-infected mice showed almost 90% granzyme B^+^ CD8^+^ T cells, we observed a near absence of granzyme B secretion in brain-CD8^+^ T cells (Figure S11B and C). However, around 30% of CD8^+^ T cells were still perforin^+^ (Figure S11B and D). Additionally, we used CD107a, as a marker of degranulation and cytotoxic potential^53^ and found no significant differences between brain and spleen CD8^+^ T cells (Fig S11F and G). Next, we measured the proportion of IFN-γ^+^ and TNFα^+^ CD8^+^ T cells. After PMA/Ionomycin stimulation, we observed high levels of IFN-γ^+^ or TNFα^+^ single-positive cells, as well as a substantial number of IFN-γ^+^ and TNFα^+^ double-positive cells, compared to those from the blood and spleen (Figure S11H-K). We also confirmed that among all CD45^hi^ immune cells in multiple organs, including brain, dura, spleen, blood, kidney, and liver, brain CD8^+^ T cells secrete IFNγ more robustly (Figure S11L). However, in mice without any pathology, achieving strong TCR stimulation, like that induced by PMA/Ionomycin, remains challenging. To further investigate IFNγ production under physiological conditions, we employed two complementary approaches: immunohistochemistry using an anti-IFNγ antibody and ELISA to quantify IFNγ levels in brain lysates. Immunostaining of brain tissues with CD8 and IFNγ antibodies revealed that approximately 10% of brain CD8^+^ T cells produce IFNγ under basal conditions (Fig S11M and N). Consistently, ELISA results showed not robust, but detectable IFNγ in the brain, which was significantly reduced in the CD8 KO mouse brain (Figure S11O). Additionally, the reduced level of IFNγ in CD8 KO mice was restored following transfer of CD8^+^ T cells from WT mice, but not after transfer of CD8^+^ T cells from IFNg KO mice (Figure S11O). These results suggest that brain-CD8^+^ T cells are a notable source of basal IFNγ in the healthy brain.

Moreover, consistent with scRNA seq results, we confirmed that brain CD8^+^ T cells are CD69^hi^ tissue resident cells (Figure S11P and Q). We next asked whether the reduced cytotoxicity observed in brain CD8^+^ T cells could be attributed to an exhausted phenotype. To investigate this, we examined multiple markers associated with CD8^+^ T cell exhaustion, including PD-1, 2B4, TIGIT, CD160, Tim3, Lag3 and Tox. We consistently observed that approximately 20% of brain CD8^+^ T cells expressed elevated levels of exhaustion markers, with the exception of Tim3 and Lag3, compared to their expressions in spleen cells. However, the overall expression levels of these markers were much lower than those observed under chronic infection (C13-infected condition). To further characterize brain CD8^+^ T cells, we compared our scRNA-seq data with a publicly available transcriptomic atlas of CD8^+^ T cells, which includes populations spanning from naive to terminally exhausted states (Figure S11V-X)^54^. Integration of these datasets revealed that brain CD8^+^ T cells are predominantly enriched in the memory-like progenitor (MP) cluster, characterized by low expression of *Gzma* and *Gzmb*, moderate levels of *Prf1* and *Pdcd1*, and high expression of *Tnfa* and *Ifng*. Taken together, our observations suggest that brain-specific CD8^+^ T cells exhibit distinct tissue-resident and effector memory-like characteristics compared to blood and dura CD8^+^ T cells, with unique gene expression profiles and functional responses shaped by brain-derived factors.

To understand how brain-specific CD8^+^ T cells acquire unique characteristics in the brain, we performed co-culture experiments with CD8^+^ T cells and tissue lysate from the brain, kidney, and lung. After isolating naive CD8^+^ T cells from the spleen and mLNs of WT mice, cells were seeded onto plates coated with anti-CD3 and anti-CD28 antibodies. Simultaneously, protease inhibitor-treated crude tissue lysate was added to the naive CD8^+^ T cell culture. After 3 days of incubation, the cells were examined for their effector functions by flow cytometry (Figure 6F). We next assessed how the brain microenvironment influences the cytotoxic potential of effector memory-like CD8^+^ T cells. To this end, we gated on CD44^+^CD62L^-^ CD8^+^ T cells and evaluated the proportion of granzyme B^+^ cells following incubation with lysates from different tissues. We observed a marked reduction in granzyme B^+^CD8^+^ T cells in cultures treated with brain lysate, whereas kidney lysate caused only a mild decrease and lung lysate had no effect (Figure 6G and H). Importantly, TCRb mean fluorescence intensity (MFI) and the proportions of Annexin V^+^ and PI^+^ CD8^+^ T cells were comparable across all conditions, indicating that the suppression of granzyme B expression was not due to T cell receptor downregulation or cell death (Figure 6H). These findings demonstrate that the brain uniquely contains soluble factor(s) capable of specifically dampening CD8^+^ T cell cytotoxic activity, supporting the notion that brain-derived cues shape a distinct functional state of tissue-resident CD8^+^ T cells.

### IFN-γ secreted by brain-CD8^+^ T cells restores behavioral abnormalities

IFN-γ is a pivotal cytokine primarily associated with immune responses against infections and tumors. Most studies regarding the role of IFN-γ in the brain focus on pathological conditions. However, emerging evidence suggests that IFN-γ also plays essential roles in modulating brain function under non-pathological conditions, underscoring its significance beyond immune responses^55,56^. While significant progress has been made in elucidating the effects of microglia and astrocyte-derived IFN-γ under pathological conditions, the precise mechanisms by which brain-specific CD8^+^ T cells utilize this cytokine to maintain brain homeostasis remain a compelling area for further exploration. Given that brain-CD8^+^ T cells secrete IFN-γ (Figure 6 and Figure S11), we investigated whether the absence of IFN-γ due to CD8^+^ T cell depletion contributes to the induction of neuronal excitatory/inhibitory ratio imbalance and behavioral abnormalities. To test this, we introduced CD8^+^ T cells isolated from WT or IFN-γ KO mice into CD8 KO mice by adaptive transfer and measured their neuronal excitability and behaviors (Figure 7A). CD8^+^ T cells were isolated from the spleen of IFN-γ KO mice and transferred to 2-wk-old CD8 KO mice. PBS was injected as a control. Six weeks later, mice underwent sociability and open field tests, as well as brain slice recordings. Surprisingly, while CD8^+^T cells isolated from WT mice restored behavioral and electrophysiological abnormalities in CD8 KO mice, compared to PBS injected CD8 KO mice, CD8^+^ T cells isolated from IFN- γ KO mice exhibited deficits (Figure 7B-F). In parallel, we transferred CD8^+^ T cells to the Severe Combined Immune Deficiency mutation (SCID) mice, which previously showed IFN-γ-dependent social deficits^56^. Consistent with CD8^+^ T cell transplantation experiments with CD8 KO mice, the transfer of CD8^+^ T cells from WT into SCID mice rescued behavioral deficits (Figure S12A-E).

**Figure 7.**
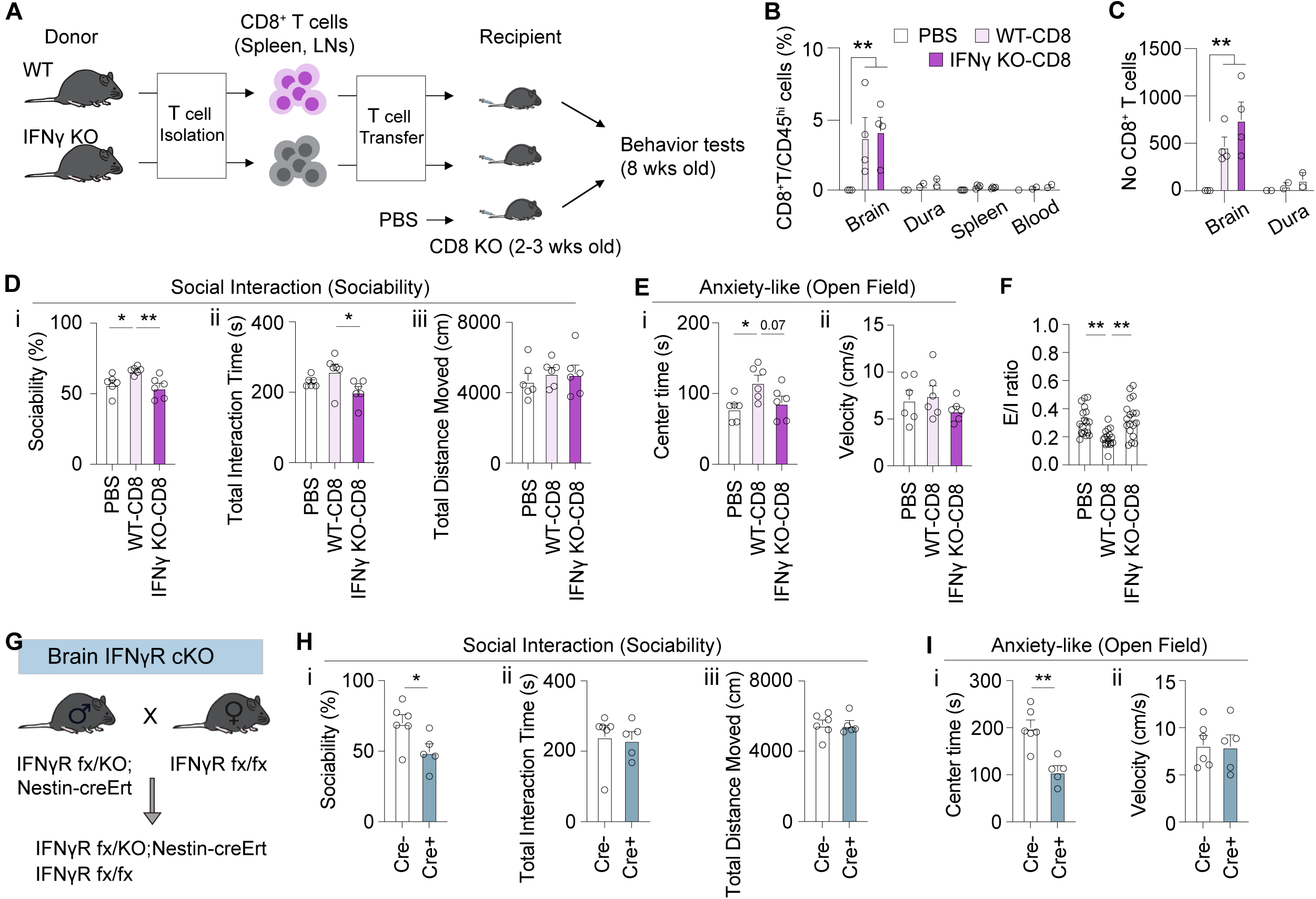
IFN-γ secreted by brain- CD8^+^ T cells restores behavioral abnormalities. (**A**) Schematic diagram of CD8^+^ T cell transfer experiment to CD8 KO recipient mice. CD8^+^ T cells were isolated from spleen and LNs of WT or Ifnγ KO donor mice. 6 weeks after T cell transfer, recipient mice were used for behavioral examinations. (**B and C**) Relative distribution (B) and absolute number (C) of repopulated CD8^+^ T cells among CD45^hi^ cells in the brain, dura, spleen, and blood of PBS (CD8 KO group), WT-CD8 (CD8 KO mice adoptively transferred with WT CD8^+^ T cells), and Ifng KO-CD8 (CD8KO mice adoptively transferred with CD8^+^ T cells from Ifnγ KO mice). (**D** and **E**) Social interaction (**D**) and anxiety-like (**E**) tests were conducted on PBS, WT-CD8, and Ifng KO-CD8 mice. (**F**) E/I ratio measured in L2/3 excitatory neurons of mPFC of each group. (**G**) Mating strategy to generate Nestin-creERT dependent IFNγR1 conditional knock-out mice. (**H and I**) Social interaction (**H**) and anxiety-like (**I**) tests were conducted on control and KO mice. (Cre-, n=6, Cre+, n=5, from 2 independent experiments). *P<0.05, **P<0.01, calculated by an One-way ANOVA (B-I). Data are shown as the mean ±SEM.

To identify the target cell population responsible for mediating IFN-γ signaling in the brain, we generated cell-type-specific cKO mice for the IFN-γ receptor (Ifngr1) in neurons (Nestin-creERT), microglia (Tmem119-creERT), and astrocytes (GFAP-creERT). Behavioral testing revealed that neuron-specific Ifngr1 cKO mice exhibited pronounced abnormalities in social interaction and anxiety-related behaviors, closely resembling the phenotypes observed in CD8-deficient mice (Figure 7G-I). In contrast, microglia-specific Ifngr1 cKO mice displayed only mild anxiety-like behaviors without social deficits, and astrocyte-specific deletion of Ifngr1 had no measurable behavioral effect (Figure S12F-K).

Taken together, these results highlight the critical role of IFN-γ produced by brain-specific CD8^+^ T cells in maintaining neural homeostasis and promoting normal behavioral outcomes through interactions with IFNGR1-expressing neurons.

## Discussion

In this study, we identified resident CD45^hi^ lymphocytes in the brain parenchyma. With diverse techniques in both neuroscience and immunology, we elucidated a previously underappreciated role of brain-specific CD8^+^ T cells in maintaining neuronal function and behavioral homeostasis, expanding the understanding of neuroimmune interactions within the CNS. Traditionally, CD8^+^ T cells are known for their cytotoxic role in peripheral immune responses, targeting and eliminating infected or malignant cells. However, our study demonstrates that within the brain, these cells contribute to neurodevelopment and behavioral regulation through non-cytotoxic mechanisms, challenging the conventional view of CD8^+^ T cell function.

Cell-to-cell communication in the brain is fundamental for the maintenance of neural networks and overall brain health. This communication involves not only neurons but also glial cells and immune cells, which together create a complex, dynamic environment. Emerging evidence highlights the crucial role of neuroimmune interactions in modulating brain function, suggesting that immune cells, such as microglia and T cells, play significant roles in synaptic pruning, neurogenesis, and the response to injury^4–10,57–60^. Understanding the diverse functions of these understudied immune cells, especially CD8^+^ T cells, is essential for gaining a comprehensive view of brain physiology and pathology.

We identified that brain-specific CD8^+^ T cells infiltrate the brain early in life. This early infiltration is followed by clonal expansion and acquisition of an effector-memory phenotype via interactions with microglia, the brain’s resident immune cells. The importance of these interactions was confirmed through genetic and pharmacological interventions that impaired microglial function, leading to disrupted CD8^+^ T cell maturation and function. These findings align with previous research indicating the critical role of microglia in supporting T-cell differentiation and function^61^ More specifically, microglia express MHCI and II molecules, enabling them to present antigens to infiltrating or resident T cells. Through this antigen presentation capacity, microglia help orchestrate adaptive immune responses in the brain and influence T cell activation, tolerance, and cytokine secretion^61^. Our genetic data provide direct evidence supporting this antigen-presenting role. Microglia-specific B2m cKO mice exhibited a pronounced arrest in CD8^+^ T cell differentiation restricted to the brain, accompanied by behavioral deficits, indicating that microglia MHC I mediated signaling is essential for local CD8^+^ T cell maturation and function. Although the specific antigens recognized by these cells remain to be defined, microglial cytosolic peptides are plausible candidates. Our data also suggest that microbial exposure and antigenic experience are key determinants of CNS CD8^+^ T cell development. In GF mice, brain CD8^+^ T cells failed to expand postnatally, in contrast to the robust expansion observed in SPF mice. Remarkably, microbial reconstitution of GF dams restored this phenotype in their offspring, while early-life Abx treatment abolished it, revealing a critical developmental window for microbiota-dependent regulation. Together, these findings highlight a model in which early-life microbial exposure provides essential antigenic or metabolic cues that shape a CD8^+^ T cell pool capable of entering and adapting to the brain, where microglial MHC-1 mediated interactions guide their local differentiation and functional maturation.

In addition to these immune-cell interactions, our data demonstrated that CD8^+^ T cell depletion disrupts mPFC circuit homeostasis, characterized by an increased E/I ratio reflecting heightened excitatory drive, reduced neuronal excitability likely due to compensatory intrinsic adaptations such as decreased input resistance, and an increase in inhibitory synapse number without changes in vGAT or gephyrin coverage, suggesting enhanced stabilization of inhibitory contact. Collectively, these findings reveal a complex interplay between immune regulation and synaptic physiology, where the absence of CD8^+^ T cells induces compensatory yet maladaptive changes in neuronal function. These results are consistent with earlier reports linking neuronal excitability to behavioral changes^62–64^.

Our mechanistic studies reveal that brain-specific CD8^+^ T cells secrete IFN-γ, which plays a critical role in maintaining neuronal excitability and synaptic function. In the absence of IFN-γ signaling, either through CD8^+^ T cell depletion or IFN-γ blockade, we observed significant synaptic dysregulation and behavioral deficits. Conversely, the reintroduction of IFN-γ in CD8^+^ T cell-depleted mice diminished these deficits in neuronal excitability and behavior, highlighting IFN-γ as a pivotal mediator in neuroimmune communication within the brain^56,65,66^. Our study further identifies neuronal IFN-γ receptor signaling as the key mediator linking brain CD8^+^ T cell activity to behavioral regulation. Cytokines are increasingly recognized for their neuromodulatory functions in the non-pathological brain. For example, cytokines such as IL-4 and IL-13, released by dura T cells, have been shown to promote cognitive function by modulating astrocyte activity^67,68^. While most studies have focused on cytokines released from dura immune cells, our research emphasizes the importance of understanding cytokine production by brain-resident immune cells in maintaining normal brain function. Our findings extend this concept by demonstrating that brain-specific CD8^+^ T cells and their cytokine, IFN-γ, are crucial for regulating neuronal excitability and behavior in the non-pathological brain. The relatively small number of CD8^+^ T cells in the brain raises intriguing questions about their role and regulation. One possibility is that the brain must balance the beneficial effects of CD8^+^ T cells with the potential risks of overactivation. An excessive immune response within the brain could lead to inflammation and neuronal damage, conditions that are detrimental to brain function. Therefore, maintaining a small population of CD8^+^ T cells could be a strategy to ensure their beneficial effects without risking overactivation. Additionally, high levels of IFN-γ secreted by a larger population of CD8^+^ T cells could disrupt neural homeostasis, leading to unintended consequences such as excitotoxicity and neuroinflammation. This delicate balance highlights the complexity of immune regulation in the brain and underscores the need for precise control of immune cell activity and cytokine levels to maintain optimal brain function. Indeed, research indicates that the effects of IFN-γ are dose-dependent, with both beneficial and detrimental outcomes based on its concentration and context. At physiological levels, IFN-γ supports synaptic plasticity and cognitive functions, as demonstrated in studies where IFN-γ enhances social behavior and cognitive performance^56,66^. However, excessive levels of IFN-γ can lead to neuroinflammation and neuronal damage. For instance, chronic overexpression of IFN-γ has been associated with neurodegenerative conditions and impaired cognitive function^69^. This dose-dependent effect underscores the necessity of maintaining IFN-γ within a narrow physiological range to harness its beneficial effects while avoiding potential neurotoxic consequences.

Our research suggests that resident immune cells in the brain, particularly CD8^+^ T cells, might play a direct role in neurodevelopmental processes. This is of particular interest given the observation that children with autoimmune diseases, such as juvenile idiopathic arthritis, celiac disease, and type 1 diabetes have an increased risk of developing neurodevelopmental and psychiatric disorders^70,71^. Similarly, children with primary immunodeficiency diseases (PIDD), including Wiskott-Aldrich syndrome and common variable immunodeficiency (CVID), also face a high risk of various psychiatric and neurodevelopmental disorders^72–74^. This association highlights a significant link between immune dysregulation and neurodevelopmental issues, such as intellectual delay, autism spectrum disorders (ASD), schizophrenia, attention-deficit/hyperactivity disorder (ADHD), and anxiety disorders. These conditions often involve disruptions in neuronal excitability and synaptic function^75–77^, which we propose may be influenced by aberrant CD8^+^ T cell activity and IFN-γ signaling. Although combined with technical difficulties, it is not possible to isolate CD8^+^ T cells in the human brain under a non-pathological condition, computational analyses can simulate the proportion of CD8^+^ T cells in the brain across different ages from bulk-RNA sequencing results. Our CIBERSORT analysis indicated the presence of CD8^+^T cells in the brain, with their numbers increasing with age (Figure S12L)^78–80^. Surprisingly, the proportion of CD8^+^ T cells was found to be reduced in patients with neurodevelopmental disorders, including ASD, schizophrenia, and bipolar disorder (Figure S12M)^81–83^. These findings suggest that dysregulation of brain-specific CD8^+^ T cells may impact resident brain cells, leading to disruptions in neuronal activities and contributing to abnormal behavior. Immune dysregulation, which affects peripheral cytokine levels, could influence brain-resident immune cells triggering a cascade of neuroimmune responses that contribute to synaptic dysfunction and behavioral abnormalities observed in these disorders. Future studies should focus on elucidating the specific molecular pathways through which CD8^+^ T cells and IFN-γ affect neuronal function and how these pathways are altered in disease states. Understanding the dual impact of peripheral and brain-resident immune cells on neurodevelopment could reveal novel therapeutic targets for preventing and treating psychiatric disorders linked to immune dysregulation.

While our NK cell profiling and TCR repertoire analyses support the absence of overt infection in SPF mice, these approaches cannot completely exclude the influence of the microbiome or nonpathogenic SPF flora on peripheral immune activation. Microbial-derived antigens and metabolites may drive low-level CD8^+^ T cell activation even in the absence of deliberate infection. Additionally, current TCR databases are limited in their coverage of antigen specificities, making it difficult to fully rule out activation by commensal or environmental antigens. We therefore acknowledge these methodological constraints as caveats when interpreting our findings related to the activation state and origin of brain-resident CD8^+^ T cells.

In conclusion, our study uncovers a critical and novel role for brain-specific CD8^+^ T cells in maintaining neuronal homeostasis and behavior. These findings emphasize the importance of neuroimmune interactions in brain function, opening new avenues for research into immune-based therapeutic strategies for neurological and psychiatric disorders. The potential to modulate CD8^+^ T cell activity and IFN-γ signaling offers promising prospects for innovative treatments aimed at restoring normal brain function and behavior in affected individuals.

## Methods

### Animals

All experiments and breeding were conducted in accordance with the guidelines of the Institutional Animal Care and Use Committee (IACUC) of the University of Pennsylvania. C57BL/6J, CD45.1 (#002014), *Ifng* KO (#002287), B6 SCID (#001913), *CD8α* KO (#002665), *β2m*^fl/fl^ (#034858), *IFN-γR1*^fl/fl^ (#025394), Tmem119-creER^T2^ (#031820), and GFAP-creER^T2^ (#012849), TEK-Cre (#008863), Nestin-CreER^T2^ (#016261) mice were purchased from Jackson Laboratory and inbred. *β2m*^fl/fl^ mice were crossed with GFAP-creER^T2^, and Tmem119-creER^T2^, TEK-creER^T2^ mice to remove β2m from astrocytes, and microglia in the brain, and endothelial cells respectively. *IFN-γR1*^fl/fl^ mice were crossed with Nestin-CreER^T2^, GFAP-creER^T2^, and Tmem119-creER^T2^ mice to remove Interferon-γ receptor specifically from neurons, astrocytes, and microglia.

Tamoxifen (Sigma, T5648) was dissolved in corn oil at 37°C overnight to a concentration of 40mg/mL. Lactating dams were administered Tamoxifen via intraperitoneal (i.p.) injection at a dose of 150mg/kg of body weight once daily. Injection was performed for five consecutive days, starting on postnatal day 5 (P5) of the pups. The pups were monitored daily for signs of distress or adverse effects following maternal injection.

For making antibiotics-treated mice, C57BL/6 pregnant female or 2-month-old male mice were treated with antibiotics mix via drinking water. The antibiotics cocktail consisted of 1g/L Vancomycin, 1g/L Amphicillin, 1g/L Neomycin, and 0.25g/L Metronidazole dissolved in autoclaved ddH_2_O. The drinking water was changed twice weekly.

Both sexes were used for Pre-weaning experiments, while only male mice were used for adult behavior experiments. Mice were housed at 22.2 °C and 52.1% humidity. Mice were given access to autoclaved food and water ad libitum and were maintained under a 12 h light-dark cycle. All mice were maintained in filter-topped cages and given autoclaved food and water.

Germ-free and gnotobiotic mice were housed in flexible film isolators (Class Biologically Clean [CBClean], WI, USA). Mouse housed at Hill Pavillion were fed autoclaved LabDiet 5021 (Cat# 0006540) ad libitum. Germ-free and gnotobiotic mice were caged on autoclaved Beta-chip hardwood bedding (Nepco, NY, USA). Sterility checks were regularly performed on isolators each month and additionally prior to any transfer of animals. Freshly collected pellets were cultured on brain heart infusion (BHI) (Oxoid, UK), NB1, and Sabouraud media for 65–70 h at 37oC under aerobic and anaerobic conditions with positive and negative control samples. Isolator sterility was confirmed externally every 3-4 months by Charles River Laboratories (NJ, USA). To generate the CMCom gnotobiotic line, whole cecal contents of a six-week-old female SPF NOD/Eα16 mouse was collected anaerobically in rPBS and 100 μl of the slurry was gavaged into six-to-eight-week-old germ-free C57BL/6J mice^41^.

No methods were used to predetermine sample sizes but, instead, sample sizes were determined by pilot experiments to assess effect sizes and variability. Genotyping of all experimental mice was conducted according to the standard protocol provided by the Jackson Laboratory.

### Immune cell preparation

To label CD45^hi^ immune cells in the vasculature of nonlymphoid organs, 3 ug of anti-CD45 (Clone:30-F11) in 300 ul of PBS was injected intravenously into mice^23^. Three minutes later, the mice were perfused with 30 ml of cold PBS under isoflurane anesthesia. After collecting the brain, dura mater was carefully collected from the skull cap84. For the dissection of the brain, olfactory bulb, striatum and thalamus, hypothalamus, hippocampus, cortex, midbrain, hindbrain (Pons and Medulla oblongata), cerebellum and choroid plexus were anatomically dissected under microscopy. At least four mouse brains were pooled for the brain dissection experiment.

The whole brain, and dissected brain parts were chopped with scissors and digested in digestion solution (10% FBS, 1 mg/ml Collagenase IV from Clostridium histolyticum (Sigma), and 50 ug/ml DNase I in RPMI 1640) with stirring with magnetic bar for 25 minutes at 37 °C. Samples were then filtered with 70um strainer and washed. Collected tissue was resuspended in 40% percoll and loaded on 75% percoll solution followed by centrifuge for 20 minutes at room temperature, without break. After removing the upper myelin layer, cells from the intermediate layer were collected and washed twice for further experiments.

Dura maters were transferred into a glass vial tube with digestion solution (10% FBS, 0.75 mg/ml Collagenase IV, and 50 ug/ml DNase I in RPMI 1640). The samples were stirred for 8 minutes at 37 °C with a magnetic bar, then washed twice for further experiments.

Cervical, inguinal, mesenteric lymph nodes and spleen were minced mechanically with slide glasses. Red blood cells were lysed using ACK lysis buffer for 2 minutes at room temperature and filtered with 40μm strainer. Blood cells were collected by heart puncture and red blood cells were lysed by treating with ACK lysis buffer for 3min at room temperature.

Single-suspended cells were stained using the LIVE/DEAD™ Fixable Near-IR Dead Cell Stain Kit (Thermo Fisher Scientific) to exclude dead cells. Surface antigens were stained for 30 minutes 4 °C with indicated antibodies after Fc blocking for 30 minutes at 4 °C with anti-mouse CD16/32 (BD) without washing (Table S1). After washing with PBS, samples were analyzed on BD FACSymphony™ A3 or BD® LSR II or Cytek Aurora. Data was collected through FACSDiva (BD Pharmingen) or SpectroFlo® (Cytek). Analysis was performed with FlowJo software (BD Biosciences).

### Intracellular cytokine staining

To measure cytokine expression in CD8^+^ T cells, single-suspended cells collected from the brain, blood, spleen, and dura were stimulated with 50 ng/ml of PMA and 500 ng/ml of ionomycin in the presence of 1x Brefeldin A in the culture medium (10% FBS, penicillin, streptomycin, and 2-Mercaptoethanol) for 4 hours in a 37°C CO_2_ incubator. After washing once, the cells were fixed with BD Cytofix/Cytoperm™ Fixation/Permeabilization Solution (BD Biosciences) for 20 minutes at 4°C. Following two washes, the cells were stained for 30 minutes at 4°C to measure IFN-γ and TNFα. To measure Granzyme B, and Perforin expression, surface-stained cells were fixed with the Foxp3 / Transcription Factor Staining Buffer (eBioscience™) for 30 minutes at 4°C, without prior stimulation. After two additional washes, intracellular antigens were stained for 1 hour at 4°C.

### CD8^+^ T cell depletion

For depleting CD8^+^ T cells in mice aged 3 wks (Juvenile Depletion) or 9 wks (Adult Depletion), littermate male mice were treated with rat IgG2b isotype control (BioXCells, Clone LTF-2, #BE0090) or anti-mouse CD8α antibody (BioXCells, Clone 2.43, #BE0061) via i.p. injection. The initial three injections were administered over six days, each at a dose of 200 ug, followed by more than four additional injections, spaced 4-5 days apart, each at a dose of 100 ug. Depletion efficiency was assessed one day after the behavior test.

For CD8^+^ T cell repopulation (brain-specific depletion), after juvenile depletion, mice were given 4 wks to allow CD8^+^ T cells to repopulate following the last injection.

### CD8^+^ T cell Adaptive Transfer

CD8^+^ T cells were isolated from spleen and lymph nodes of 6 to 8-wk-old C57BL/6 WT mice, Ifng KO or CD45.1 congenic mice. Total CD8+ T cells were sorted using flow cytometry (Symphony S6, BD) with purity 98%> after depleting B cells with magnetic beads using anti-B220 antibody. 3 x 10^6^ sorted CD8^+^ T cells were resuspended in sterile PBS and transferred intravenously into 2-wk-old littermate *CD8α* KO mice or 3-wk-old SCID mice. To check the migration time window, brain, dura mater immune cells were analyzed as described above 6weeks after transfer. For CD8aKO mice reconstituted with wildtype or IFNg KO CD8 T cells, behavioral tests were performed when the recipient mice reached 8 wks of age to assess the functional consequences of CD8^+^ T cell reconstitution. After the test, brain, dura mater, blood, spleen cells were analyzed as described above to evaluate reconstitution rate of transferred CD8 T cells. The specific behavioral tests used are described in detail in subsequent sections.

### Microglia depletion

Newborn C57BL/6 littermate pups were treated with DMSO (for control) or 30 mg/kg of CSFR1 inhibitor (Pexidartinib, PLX-3397; MedChemExpress) diluted in a 50:50 PBS-polyethylene glycol solution via S.C. injection starting on their post-natal day 3 (P3), with injections administered daily^85^. The mice were sacrificed for FACS analysis on P18.

### LCMV induction

LCMV Armstrong and cl13 were propagated and tittered as previously described^86,87^. For LCMV Armstrong infection, C57BL/6J mice (12-wk-old; male) were infected i.p. with 2×10^5 PFU of LCMV Armstrong (day 0). Tissues were harvested on day 8 p.i. For LCMV cl13 infection, C57BL/6N mice (6-wk-old; female) were adoptively transferred with 500 wildtype P14 cells (day −1), GK1.5-treated (day −1 and 1), and infected i.v. with 4×10^6 plaque-forming units (PFU) of LCMV cl13 (day 0). Tissues were harvested on day 40 p.i.

### In vitro CD8^+^ T cell culture with organ lysates

Brain, lung, and kidney were collected after transcranial perfusing mice with 30ml cold PBS. The tissues were finely chopped using scissors in culture medium (RPMI-1640 supplemented with 10% fetal bovine serum, 100 units/mL of penicillin, 100 μg/mL of streptomycin, 250 ng/mL of amphotericin B) supplemented with protease inhibitor cocktail. Tissue crude lysate was prepared by mixing chopped tissue with culture medium supplemented with protease inhibitor cocktail in 40ml per gram of tissue weight. The mixed lysate was centrifuged at 600 × g for 5 minutes at 4°C. The supernatant was carefully collected and filtered through a 0.45 µm filter. Naive CD8^+^ T cells were isolated from spleen and lymph nodes using the EasySep™ Mouse Naive CD8^+^ T Cell Isolation Kit (Stemcell Technologies, Cat. #19858) according to the manufacturer’s instructions. Purity of the isolated cells was confirmed by flow cytometry, achieving a purity of over 98%. A total of 100,000 isolated naive CD8^+^ T cells per one well were seeded onto the anti-CD3ε/CD28 antibody-coated 96 well plates and incubated in culture medium. 40ul of tissue lysate or protease inhibitor-supplemented culture media was added to each well. The cells were incubated for three days at 37°C in a 5% CO₂ before proceeding to further analyses.

### Interferon gamma (IFN-γ) Measurement in Brain Tissue

Mice were perfused with 30ml of cold PBS. The brain was collected, weighed, and homogenized in cold PBS at a ratio of 1.4ml per gram of tissue, using a disposable pestle in a 1.5ml tube. The homogenate was centrifuged at 6000xg for 5min at 4°C. The supernatant was collected and analyzed for measuring IFN-γ concentration using ProQuantum Immunoassay (Invitrogen), following the manufacturer’s instructions. qPCR was performed using QuantStudio 6 Real-Time PCR system (Applied Biosystems).

### Immunohistochemistry

Mice were exposed to either an empty pencil cup or a novel mouse under the pencil cup for 10 minutes. One hour later, mice were transcranial perfused with cold paraformaldehyde (PFA) (4% in PBS). Brains were kept in PFA overnight at 4°C and cryoprotected in a 30% sucrose solution. The sections were sliced at 18 mm using a cryostat (Leica). Slices were permeabilized with a blocking solution containing 0.1% Triton X-100 and 2% goat serum in PBS for 1 hour at room temperature. Subsequently, they were incubated overnight at 4°C with primary antibodies: rat anti-mouse CD8α (1:500, 14-0081-82, eBioscience), rabbit anti-c-Fos (1:1000, #2250, Cell Signaling), rabbit-anti-mouse CD31 (1:100, AB124432, Abcam), rat anti-mouse CD45 (1:500, 103101, Biolegend), goat anti-mouse IBA1 (1:1000, ab5076, Abcam), chicken anti-GFP (1:500, GFP-1020, Aveslabs), anti-mouse IFNg (1:100, MAB485, R&D), rabbit anti-mouse NeuN (1:1000, ab177487, Abcam), anti-Gephyrin (1:500, #147021, SYSY), and anti-vGAT (1:500, #131001, SYSY). The following day, slices were incubated with fluorescently conjugated secondary antibodies (Invitrogen) for 1 hour at room temperature, along with DAPI (1:2500, Thermo Fisher). Images of stained slices were acquired using a confocal microscope (LSM 810, Carl Zeiss) with a 10x, 20x, or 40x objective lens.

### Quantification and statistical analysis

Data were presented as mean ± standard error of the mean (SEM). No statistical methods were used to predetermine sample size. The quantification of cFos immunopositive cells in the BLA, hippocampus, mPFC, M1, S1, and ECT cortex was performed through drawing each ROI and using the function “Analyze Particles” in ImageJ. Specifically, the quantification of cFos^+^NeuN^+^ and c-Fos^-^NeuN^+^ cells in the mPFC was quantified in the rectangle ROI (415 μm width X 830 μm height). The number of NeuN-positive cells in the mPFC was quantified in the layer specific rectangle ROIs (L1: 120 μm, L2/3: 110 μm, L5a: 180 μm, L5b: 190 μm, L6: 150 μm width with 250 μm height). The percent of the image covered by GFAP- or Iba1-positive pixels in mPFC was calculated by dividing the total area of GFAP or Iba1 by the total area of the full photomicrograph.

For the analysis in microglial morphology, it was conducted based on the previous quantification studies^88,89^. Briefly, the microglial morphology was quantified at a single microglia level (cell body size, number/average length of branches, number of junctions, length of longest-shortest path, endpoints per microglia, circularity, fractal dimension, density, and span ratio) in ImageJ. After the immunohistochemistry and imaging of Iba1 marker, the photomicrographs were converted to binary, and the thresholds were adjusted. The selected microglia in the binarized image were isolated using multiple ROIs and were skeletonized for skeletal analysis and for fractal analysis.

### Mathematical modeling of tissue-resident CD8^+^ T cell population dynamics

To describe the population dynamics of naïve (N), central memory (C), and effector memory (E) CD8⁺ T cells, we formulated a system of ordinary differential equations (ODEs) capturing the migration, differentiation, and proliferation processes. We assumed that the differentiation rates of CD8⁺ T cells remain constant over time. The model is defined as:

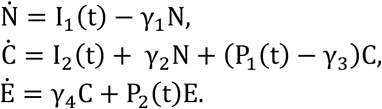

Where 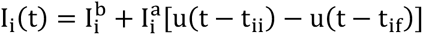, and 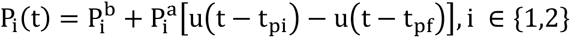. Here, I_i_(t) denotes the number of CD8**^+^** T cells migrating to the organ per time for naïve (i=1) and central memory (i=2) cells, while P_i_(t) represents the proliferation rate of central (i=1) and effector (i=2) memory CD8**^+^** T cells. Step function u(t) was employed to capture the rapid increase in migration and proliferation that occurs only during weaning period. 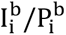 represents the baseline of migration/proliferation rate, and 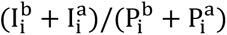 represents the maximum migration/proliferation rate between time t_ii_/t_pi_ and t_if_/t_pf_. The initial condition of N, C, and E is zero. This system has an exact analytical solution. For example, the solution of N(t) is:

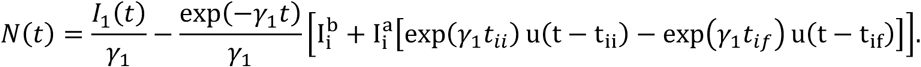

To obtain biologically plausible parameters consistent with experimental CD8⁺ T cell counts, we employed a genetic algorithm. The cost function was defined as the mean squared difference between model predictions and experimental data. We selected following parameters of the model: for brain, γ_1_ = 5.32, γ_2_ = 0.373, γ_3_ = 4.84, γ_4_ = 4.84, 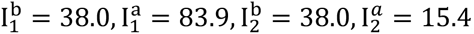, t_ii_ = 0.400, t_if_ = 1.40, 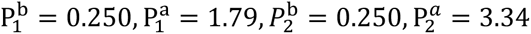, t_pi_ = 0.400, t_pf_ = 1.40. And for the meninges, γ_1_ = 5.32, γ_2_ = 0.373, γ_3_ = 4.84, γ_4_ = 4.84, 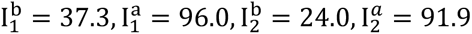, t_ii_ = 0.220, t_if_ = 0.820, 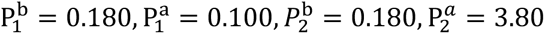, t_pi_ = 0.220, t_pf_ = 0.820.

### Deconvolution of public human brain bulk RNA-seq data for Immune cell composition analysis

To estimate the immune cell composition in bulk RNA-seq profiles of human brain samples, CIBERSORTx was applied with the LM22 signature matrix^78,79^. Batch correction was performed in B-mode, while quantile normalization and absolute mode were disabled, and 1,000 permutations were used for all analysis. For the analysis in Fig. S12K, we used data collected from PsychENCODE, specifically RPKM RNA-seq data^80^. We excluded embryos and infants whose cause of death was sudden infant death syndrome, resulting in a total of 17 samples ranging from 1 year to 40 years of age. For Fig. S12L, RPKM RNA-seq data were obtained from the recount2-brain database^81^, specifically from projects SRP035524 and SRP007483^82,83^.

### Single-cell RNA sequencing

#### Cell sorting and scRNA-Seq library construction

Three-month-old C57BL/6 mice (n = 5) were transcardially perfused with cold PBS after staining the vascular circulating immune cells as previously described^23^. Blood samples were collected via cardiac puncture before perfusion. The blood samples underwent red blood cell lysis for 4 minutes at room temperature. Brains were physically chopped and underwent digestion with Collagenase VI and DNase I, followed by Percoll gradient centrifugation as outlined in above. Single-cell suspensions from the brain and blood were stained as previously described. The stained cells were washed and FACS sorted by BD FACS Aria II instrument to isolate live blood CD45^hi^ and brain CD45^hi^ immune cells (purity > 98%). After one additional wash, the FACS-sorted cells were encapsulated using a Single Cell 3′ Reagent Kit (10X Genomics V3 Chemistry) following the manufacturer’s instructions. Libraries were subsequently sequenced on a NovaSeq 6000 platform.

#### scRNA-seq computational analysis

The initial processing of scRNA-seq data was performed using the Cell Ranger Pipeline (v.7.0.0, 10X Genomics^97^). Base call files were demultiplexed to FASTQ files using *cellranger mkfastq* with Illumina bcl2fastq2 software (v2.20.0.422). The resulting reads were aligned to the mm10 transcriptome (refdata-gex-mm10-2020-A) using *cellranger count*. Subsequent analysis were performed in R (4.4) using Seurat (v4.0.6).

We applied dataset-specific quality control thresholds: Unique genes per cell: > 300 (brain male), > 350 (blood male), >350 (brain female); < 7,300 (brain male), <6,000 (blood male), < 6,000 (brain female). Mitochondrial gene percentage: < 5% (brain male), < 7% (blood male), < 5% (brain female). Largest gene percentage: < 15% (brain male), < 10% (blood male), < 15% (brain female).

Doublets were removed using DoubletFinder (v2.0.3)^78^ with the following parameters: pN = 0.2 (brain male), 0.15 (blood male), 0.15 (brain female); pK = 0.11, 0.08, 0.005; and dp = 0.055, 0.075, 0.025, respectively.

Data were normalized and scaled using SCTransform (v2)^98^, identifying 3,000 variable features after excluding mitochondrial, ribosomal, and hemoglobin genes. Principal component analysis (PCA) was performed on all genes using *RunPCA*. The top principal components used for UMAP and Shared Nearest Neighbor (SNN) clustering were: 32 (brain male), 20 (blood male), and 27 (brain female). The Louvain algorithm was applied to the SNN graph via *FindClusters*. Microglia clusters were excluded based on *Siglech*, *Tmem119*, and *Aif1* expression.

Integration of datasets (brain male vs. blood male and brain male vs. brain female) was performed using Seurat’s *IntegrateData* function. Integration features and anchors were identified with *SelectIntegrationFeatures*, *PrepSCTIntegration*, and *FindIntegrationAnchors*, excluding mitochondrial, ribosomal, and hemoglobin genes. A total of 3,000 anchors were used. PCA was recomputed, and the top 31 components were used for UMAP and SNN clustering. Louvain clustering resolution was set to 1.0 (brain male vs. blood male) or 0.8 (brain male vs. brain female).

Differential expression was tested using the Wilcoxon rank-sum test with Bonferroni correction (*FindMarkers*). Genes were considered significant if they had a log2 fold change > 0.5 and were expressed in at least 20% of cells in the cluster.

### Trajectory analysis

To investigate dynamic transcriptomic changes in CD8⁺ T cells across the blood, dura mater, and brain, we employed the Monocle2 R package (v2.32.0)^99^. The Seurat objects containing raw RNA count data for CD8⁺ T cell subsets were converted into the Monocle2 objects via the *as.CellDataSet* function. The objects are processed with *estimateSizeFactors* and *estimateDispersions* function to normalize for differences in mRNA recovered across cells. Dimensionality reduction was performed using the *DDRTree* method implemented in *reduceDimension*, and cells were subsequently ordered along pseudotime using the *orderCells* function. The blood-derived CD8⁺ T cells were selected as the root state, representing the earliest transcriptional stage. To identify pseudotime-associated transcripts, the *differentialGeneTest* function was applied with the model formula *∼sm.ns(Pseudotime)* in the *fullModelFormulaStr* setting. Genes showing significant association with pseudotime (p < 0.05) were retained as key dynamic markers describing trajectory progression. Visualization of gene expression dynamics along pseudotime was achieved using heatmaps generated by the *plot_pseudotime_heatmap* function.

### Gene regulatory network analyses

To infer differential genetic regulatory interactions between TFs and target genes between in CD8^+^ T cells from the blood, dura mater, and brain, we employed SCENIC (v1.3.1) and pySCENIC (v0.12.1)^100,101^. To generate the candidate modules, we used the standard pySCENIC workflow with default parameters: 1. Co-expression module inference: GRNBoost2 was applied to the transcriptomic data to calculate co-expression modules, producing an adjacency matrix linking each TF to its target genes, with an associated importance score. 2. Motif enrichment analysis: Modules were refined by retaining target genes whose promoter regions contained the DNA motif specific to the corresponding TF. The v9 mm10 cisTarget databases (https://resources.aertslab.org/cistarget/) were used. 3. Regulon activity scoring: The activity of each regulon was quantified using AUCell. Regulon activity matrices were scaled using Seurat’s *ScaleData* function. Differential regulons among samples were identified using the Seurat *FindMarkers* function with default settings. The top 30 regulons ranked by variance across all samples are visualized in Figure 6E.

### Processing and analysis of public scRNA-seq datasets

Publicly available single-cell RNA sequencing datasets were reanalyzed to compare and validate transcriptional profiles of immune cells across physiological and infected conditions. Raw or processed count matrices were downloaded from the Gene Expression Omnibus (GEO) and processed using the Seurat package in R as described above.

Dura mater T cell dataset^45^: The scRNA-seq data (GSE144173) from mouse dura mater T cells were processed. We applied the filters: unique genes per cell: > 500, mitochondrial gene percentage: < 5%, largest gene percentage: < 10%. These cells were re-clustered using 25 principal components and a resolution of 0.7. Among these, CD8⁺ T cell clusters were selected based on Cd8a expression. For downstream comparison, this dataset is integrated with CD8⁺ T cell subsets from brain and blood datasets, by *SelectIntegrationFeatures*, *FindIntegrationAnchors*, and *IntegrateData* function. PCA and UMAP were performed using the top 30 principal components, Louvain clustering resolution was set to 1.0. LCMV dataset^54^: The processed CD8⁺ T cell scRNA-seq dataset from spleens of LCMV-infected mice was downloaded from the repository provided in the paper. The annotated Seurat object containing naïve, effector, memory, and exhausted CD8⁺ T cell states was used for comparative projection analysis with brain and blood CD8⁺ T cell subsets. Integration was performed using *SelectIntegrationFeatures*, *FindIntegrationAnchors*, and *IntegrateData*. PCA and UMAP were computed on the top 50 principal components, and Louvain clustering resolution was set to 0.9. Cluster annotations were reassigned across datasets.

Microglia dataset^36^: The processed single-cell transcriptomic dataset of the mouse brain (GSE121654) was reanalyzed to investigate hypothalamus cell’s H2-D1 expression along with age.

Brain cell dataset^37^: The processed single-cell transcriptomic dataset of the mouse hypothalamus (GSE132355) was reanalyzed to investigate hypothalamus cell’s H2-D1 expression along with age. All datasets were visualized using Seurat. Gene expression comparisons were performed using the Wilcoxon rank-sum test, and results were adjusted for multiple testing using the Bonferroni method.

### Bulk-RNA sequencing analysis

Adult male mice were transcardially perfused with ice-cold PBS right after social target stimulation, and the whole brain were collected. Brain immune cells were collected as described above. Collected cells were stained with Livedead and antibodies against CD11b (M1/70), CD45 (30-F11). CD11b^mid^, CD45^mid^ microglia cells were sorted using BD Symphony S6 (Purity >98%). Total mRNAs were collected using the RNeasy Mini Kit (# 74104, Qiagen). Raw data was generated using NovaSeq PE150 (Novogene). Raw reads underwent quality control checks using FastQC (version 0.11.9), and adapters were trimmed using Trim Galore (version 0.6.10). The reads were subsequently aligned to the mouse reference genome mm10 using the STAR aligner (version 2.7.2)^90^. Differential gene expression analysis was conducted using the R package DESeq2 (version 1.36.0) to identify genes that exhibited differential expression between the Iso and α -CD8 groups^91^. The threshold for significant differential expression was set at a FDR-adjusted p-value < 0.05 and absolute log2 fold change > 0.5.

For the enrichment analysis of gene sets, we used Metascape (www.metascape.org) for biological pathway analysis^92^. Specifically, we employed KEGG Pathway, WikiPathways, Canonical Pathways, BioCarta, Hallmark, and Reactome gene sets. The gene lists used to characterize microglia from the Iso and a-CD8 groups into HOM, M1, M2, and DAM categories is provided in Supplementary Table S4.

### TCR sequencing analysis

To compare TCR repertoire in CD8^+^ T cells between blood and the brain, CD8^+^ T cells from each organ were isolated as described above. Ten C57BL/6J male mice at 0.5 mo of age (n=1, pooled from 10 mice for 1 independent experiment) and ten C57BL/6J male mice at 2 mo of age (n=2, each pooled consisting of 10 mice, from 2 independent experiment) were used for sample preparation. DNAs were collected using the QIAamp DNA Micro Kit (#56304, Qiagen). Raw data were analyzed using ImmunoSEQ to identify productive clones^93^. The amino acid length of the complementary determining region 3 (CDR3) was analyzed and plotted using the immunarch (0.9.0) (https://doi.org/10.5281/zenodo.3367200). The clonal diversity of the T cell was calculated using the Chao1 estimator, which provides a non-parametric asymptotic estimate of species richness, indicating the number of species in a population. Clonality analysis was conducted to count the clonotypes using *repClonality* function with the method set to ‘rare’. To verify that mice are the non-infected condition, we compared our TRB repertoire to public database, VDJDB, which stores several pathogen-related repertoires from various studies^35^. We merged the TRB repertoire data of 2-mo-old mouse’s spleen, influenza-infected as the positive control and the non-infected as the negative control^94^. We calculated the Infection Similarity Index (ISI), quantitative metric designed to evaluate the proportion of T cell receptor (TCR) clonotypes in a given sample that are associated with infectious diseases, calculated as the proportion of sequences overlapping with VDJDB among the sample’s unique amino acid sequences. Mathematically, the ISI is expressed as:

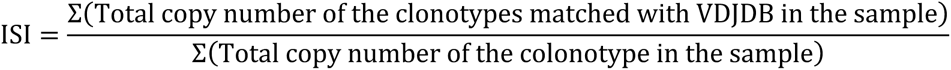

A higher ISI indicates a greater similarity between the sample’s TCR repertoire and known disease-associated clonotypes, suggesting a higher likelihood of infection.

To compare TCR repertoire in CD8^+^ T cells between the brain and peripheral organs, 2-mo-old C57BL/6J male mice (n=1, pooled from 2 mice) were. Immune cells from brain, mesenteric lymph node, lung, and liver were isolated as described above. Large intestine was flushed with PBS and cut longitudinally. Washed tissue was cut in 1cm pieces and incubated with stir bar in 37’C 10mM EDTA, 2% FBS suppled RPMI media for 20min. Incubated tissue was washed twice and chopped in fine pieces. Chopped tissue was incubated with digestion solution containing 10% FBS, 1 mg/ml Collagenase IV from Clostridium histolyticum (Sigma), and 50 ug/ml DNase I in RPMI 1640 media and stir bar for 40 minutes at 37 °C, then washed twice and centrifuged through a 40% percoll on 75% percoll solution for 20 minutes at room temperature, without break. Intermediate immune cells were collected and washed twice. Collected immune cells from each organ were stained with LiveDead and antibodies against CD45 (30-F11), CD8α (53-6.7), TCRβ (H57-597), CD11b (M1/70). CD8^+^ T cells were sorted using BD Symphony S6 (purity >98%). RNAs were extracted from the sorted cells using the RNeasy Mini Kit (# 74104, Qiagen). Raw data were analyzed by iRepertoire Inc. (Huntsville, AL, USA) to identify productive clones. Subsequent analysis is performed with Immunarch. Chao1 estimator was used for measuring the clonal diversity of the CD8+ T cell, and the function repClonality with the method set to ‘rare’ was used for measuring, with customed parameters.

### Developmental milestones in mouse pups

Each pup was removed from the cage and placed on a clean mat to assess the postnatal behavioral development. All examination was performed on all even days (P4 to P16) as previously described*95,96*. <u>Eye opening</u>: the day when both eyes were open was recorded. <u>Tactile startle reflex</u>: the presence of reflex to a gentle puff of air through syringe plunger was assessed. <u>Grasp reflex</u>: the grasping reflex to a home cage grid was observed. <u>Cliff avoidance</u>: the pup was placed on the edge of pipette tip box with its forepaws and nose hanging over the edge and the time the pup turned away from the edge was measured. <u>Negative geotaxis</u>: the pup was placed head down on a 45-degree incline and the time the pup turned to head up was measured. <u>Surface righting</u>: the pup was held on its back and the time the pup flipped over to prone position was assessed.

### Adult mouse behavioral analysis

Male mice were tested during the light cycle in a room with lighting maintained at 230 lux. Mice were transferred to the testing area at least one hour before the initiation of experiments. Tracking of mouse behavior was done using the EthoVision XT (Noldus) tracking system. Anxiety, social interaction, and novel objective cognition tests were conducted as previously described with minor modifications^43^.

#### Anxiety test-Open field test

Anxiety level was determined using an open field test. Mice were placed in the center of the open arena of the chamber (40 cm x 40 cm x 40 cm), and they were allowed to move freely for 10 minutes. All behavior was monitored with a digital camera, and the time spent in the center area (20 cm X 20 cm within the center) and the average velocity traveled were collected and analyzed by the software EthoVision XT (Noldus).

#### Anxiety test-Elevated Zero Maze

Exploration of the elevated zero maze is sensitive to anxiolytic drugs and some antidepressants; thus, it is used routinely to assess anxiety-related behavior. The maze consists of a 2.5-inch wide circular track with 2 walled and 2 open sections, elevated 12 inches above the floor. After a 30-minute habituation to the procedure room, a 5-minute trial started by placing a mouse in the middle of a walled section. Digital recordings fo the trials were processed for automated analysis by ANYmaze software (Stoelting Co) to generate the percent time spent in the open sections of the maze and distance traveled.

#### Social interaction

Sociability was measured using a three-chamber social approach test, which consisted of habituation and social interaction sessions (10 minutes). The arena was constructed of black acrylic (40 cm x 30 cm x 30 cm). Wire cups were placed in the back left and right corner. On day 0, mice were habituated to the arena for 10 minutes. Immediately after habituation, mice were singly housed. On day 1, mice were placed in the center of the arena and allowed to freely explore. Following 10 minutes, mice were confined to the center of the arena. An inanimate object or a male conspecific was placed beneath the wire cups. Mice were then allowed to freely explore the arena for 10 minutes. Sociability was defined as interaction time with the social target divided by total interaction time and expressed as a percentage.

#### Novel Objective Recognition

The mouse was placed into a clean rat cage with bedding to acclimatize for an hour. Two identical objects were placed in the cage for 10 minutes (process for making the familiar object). An hour after removing the familiar objects, one familiar object was reintroduced with a novel object at the same time. Count the interaction time separately between familiar and novel objects. The interaction time is defined as sniffing or direct nose contact, only in case the total interaction time is over 30 seconds. If the total interaction time did not reach 30 seconds, it was discarded. No innate preference within the object pairings was confirmed before the experiment.

### Brain slice recording

Mice were deeply anesthetized and transcardially perfused with ice-cold artificial CSF (aCSF) containing (in mM): 124 NaCl, 2.5 KCl, 1.2 HaH_2_PO_4_, 24 NaHCO_3_, 5 HEPES, 13 Glucose, 1.3MgSO_4_, 2.5 CaCl_2_. After perfusion, the brain was quickly removed, submerged and coronally sectioned on a vibratome (VT1200s, Leica) at 250 μm thickness in ice-cold aCSF. Slices were transferred to NMDG-based recovery solution at 32°C of the following composition (in mM): 92 NMDG, 2.5 KCl, 1.2 NaH_2_PO_4_, 30 NaHCO_3_, 20 HEPES, 25 Glucose, 5 Sodium ascorbate, 2 Thiourea, 3 Sodium pyruvate, 10 MgSO_4_, 0.5 CaCl_2_. After 12-15 minutes of recovery, slices were transferred to room temperature aCSF chamber (20-22°C) and left for at least 1 hour before recording. Following recovery, slices were placed in a recording chamber, fully submerged at a flow rate of 1.4-1.6 mL/min, and maintained at 29-30°C in oxygenated (95% O2, 5% CO_2_) aCSF.

In voltage-clamp recordings, recording pipettes were fabricated by pulling borosilicate glass (World Precision Instruments, TW150-3). These pipettes exhibited a tip resistance ranging from 3 to 5 MΩ when filled with an internal solution comprising the following concentrations (in mM): 130 CsMeSO_4_, 5 CsCl, 10 HEPES, 2.5 MgCl, 0.6 EGTA, 1 QX-314, 10 Na-Phosphocreatine, 4 NaATP, and 0.3 NaGTP, with a pH adjusted to 7.3-7.4 using CsOH. Pyramidal neurons were identified under visual control using IR-DIC optics (Olympus, BX51). During the experiments, neurons were voltage clamped at two specific membrane potentials: −56 mV, which corresponds to the chloride (Cl-) reversal potential, and +3 mV, representing the cation (cation reversal potential) reversal potential. These membrane potentials were determined in prior experiments. The stimulus electrode was positioned at the boundary between Layer VI and corpus callosum fibers. The stimulation intensity was maintained consistently throughout the entire excitatory/inhibitory (E/I) ratio experiment.

For current-clamp recordings, the recording pipette was filled with an internal solution containing (in mM) 140 K-gluconate, 5KCl, 0.2 EGTA, 2 MgCl_2_, 10 HEPES, 4 MgATP, 0.3 NaGTP, 10 2Na-Phosphocreatine (pH adjusted to 7.3-7.4 using KOH). To evaluate the passive membrane properties of the cells, they were maintained at their resting membrane potential. Resting membrane potential and input resistance were captured immediately after whole-cell configuration to minimize internal dialysis. A step-wise protocol was then applied for 500 ms, where each sweep involved an increase of +20 pA in the current injection. This step-wise protocol continued until a maximum current injection of 300 pA was reached. Each sweep in the step-wise protocol was separated by a duration of 15 seconds.

Recordings were performed using a MultiClamp 700B (Molecular Devices) and Igor7 (WaveMetrics; recording artist addon, developed by Richard C Gerkin, Github: https://github.com/rgerkin/recording-artist), filtered at 2.8kHz and digitized at 10 kHz. Axon terminals were stimulated with a brief (0.2 ms) pulses using isoflex isolator. Input and series resistance were monitored continuously, and experiments were discarded if either parameter changed by >20%. Data were analyzed using Igor7.

## Data availability

The scRNA sequencing have been deposited in the NCBI Gene Expression Omnibus (GEO) database under the accession number GSE244887 and the bulk RNA sequencing data have been deposited in the NCBI Gene Expression Omnibus (GEO) database under the accession number GSEXXXXX.

## Resource Availability

### Lead contact

Further information and requests for resources and reagents should be directed to and will be fulfilled by the lead contact, Yeong Shin Yim.

### Materials availability

This study did not generate new unique reagents.

### Data and code availability

scRNA-seq data have been deposited at GEO and are publicly available as of the date of publication. Any additional information required to reanalyze the data reported in this paper is available from Yeong Shin Yim upon request.

### Statistical analysis and reproducibility

Graphs and statistical analyses, as specified in figure legends, were obtained using GraphPad Prism or RStudio. All attempts at replication were successful and no data were excluded from the analyses.

## Acknowledgements

We thank members of the Yim laboratory and Dr. Mariko Bennets for comments on this manuscript. We would like to thank The Neurobehavior Testing Core at UPenn/ITMAT and IDDRC at CHOP/Penn U54 HD086984 for assistance with the behavior procedures. This work was supported by the Simons Foundation Autism Research Initiative (Y.S.Y.), Suh Kyungbae Foundation (Y.S.Y.), Chan Zuckerberg Initiative (Y.S.Y.) and National Institutes of Health grants 1DP2AG067492-01 (C.A.T.).

## Author Contribution

Designed the experiments and/or provided advice and technical expertise, K.P., J.J., S.J., F.C.B., J.E.W., M.A.S., C.A.T., M.V.F., and Y.S.Y. Immune cell profiling experiments, K.P., S.H., S.F.N, J.N.F., S.L. Bioinformatic analysis, J.J. Behavioral experiments, S.J., T.O.C., C.F.H. Electrophysiological experiments, K.C. Manuscript preparation, Y.S.Y, K.P., J.J., S.J. All authors edited the final manuscript.

## Competing Interests

The authors declare no competing interests.

---

**Figure S1. Gating strategy for multi-color flow cytometry analysis of each immune cell type in the brain, dura, and choroid plexus (ChP).**

(**A**) LiveDead-NearIR, Tissue CD45-PerCP-Cy5.5, Blood CD45-PE-Cy7, B220-BV650, γδTCR-FITC, ST2-BV605, NK1.1-PE-Cy5, Ly6G-BV785, CD11b-BUV496, CD11c-BUV737, Ly6C-BV711, CD117-PE, TCRβ-BV421, CD8α-BUV395, and CD4-PE/Dazzle594, CD19 BUV805 - conjugated antibodies were used for staining.

(**B**) Quantification of CD45^hi^ immune cells in the brain parenchyma of 3-month (mo)-old male and female mice.

(**C**) Proportion of each immune cells in the brain parenchyma of 3-month (mo)-old male and female mice. P value was calculated by a student t-test (**B**) and Two-way ANOVA with Sidak’s multiple comparisons test (**C**). Data are shown as the mean ±SEM.

**Figure S2. Immune cells in the spleen didn’t show immune activation signatures.**

(**A**) Gating strategy for flow cytometry analysis of immune cells from spleen.

(**B** and **C**) Representative FACS plots and quantification of Naive, Tcm, Tem subpopulation of CD4^+^ T cells (**B**) and CD8^+^ T cells (**C**) in spleen of WT mice.

(**D**) Relative distribution of immune cells in the spleen of WT mice used for this study.

**P<0.01, calculated by One-way ANOVA with Dunnett’s multiple comparisons test (B and C). Data are shown as the mean ±SEM.

**Figure S3. Tissue-resident CD45^hi^ immune cells were observed in the brain parenchyma, related to Figure 1**.

(**A**) Representative images for CD45^hi^ immune cell staining. Scale bar, 50um

(**B** and **C**) low (**B**) and high (**C**) magnitude images to show the localization of CD45^hi^ immune cells in the brain, including olfactory bulb (OB), cerebral cortex (Cbx), hippocampus (Hippo), striatum and thalamus (S+T), hypothalamus (Hypo), midbrain (MB), hindbrain (HB), and cerebellum (Cbll). Scale bar, 1mm (B) and 0.25mm (C).

**Figure S4. Brain CD45^hi^ immune cells show different proportional changes across ages and tissues, related to Figure 1**.

(**A-F**) Proportion (i) and number (ii) of each immune cell types across different ages, including 0.5 mo, 1 mo, and 3 mo, in the brain parenchyma (A), dura (C), ChP (D), blood (E), and spleen (F), as well as cells from circulating blood in the brain (B). All data were acquired by flow cytometry.

*P<0.05, **P<0.01, calculated by linear regression (A) and Two-way ANOVA with Dunnett’s multiple comparisons test (B). Data are shown as the mean ±SEM.

**Figure S5. Brain CD45^hi^ immune cells show different transcriptional profiling compared to the one in blood, related to Figure 1.**

(**A** and **F**) UMAP plots to show the cluster of each immune cell from blood-male and brain-male (**A**) and female-brain and male-brain (**F**).

(**B** and **G**) UMAP of the cell type marker expression, B cells (CD79a), CD4^+^ T cells (Cd4), CD8^+^ T cells (Cd8a), gd T cells (Trdc), NK cells (Klrb1c), Neutrophils (S100a8), DC (Mrc1) and Macrophages (Fn1), in the blood-male and brain-male (**B**) and the female-brain and male-brain (**G**).

(**C** and **H**) Bar graph showing the proportions of the cell types of CD45^hi^ immune cells in the blood-male and brain-male (**C**) and female-brain and male-brain (**H**).

(**D** and **I**) Strip plot showing the differentially expressed gene in each immune cell type between blood-male and brain-male (**D**) and female-brain and male-brain (I). For genes enriched in brain-male cells (D) and male-brain **(I**) are positive in fold change.

(**E** and **J**) Violin plots showing selected marker expressions for effector-like genes, including *Cd19, Cd27* for B cells, *Cd40, Sell, Cd44, Il2ra* for T cells, and *Itgax, Itgam, Cd68, Cd80, Cd83, Cd86, Cd209d, Mrc1, Cxcr2, Mertk* for myeloids, in the male-brain vs male-blood (**E**), the male-brain vs female-brain (**J**).

**Figure S6. Brain-specific CD8^+^ T cells migrate into the brain in a naive state during early life, related to Figure 2.**

(**A**) Relative distribution of CD8^+^ T cells in the blood at different ages (0.5 mo, 1 mo, 2 mo, and 3 mo). All data were acquired by using flow cytometry.

(**B**) CD4/CD8 ratio showing the expansion of CD8^+^ T cells specifically in the brain. The red dotted line indicates a 1:1 ratio between CD4^+^ T and CD8^+^ T cells. Values above the red line indicate a higher proportion of CD4^+^ T cells. All data were acquired using flow cytometry.

(**C**) Number of CD8^+^ T cells increased over time in both the brain and dura, but with different slopes.

(**D**) Relative distribution of CD8^+^ T cells across mouse brain regions at 0.5 mo, 1 mo, and 3 mo, as determined by flow cytometry. Relative CD8^+^ T cell distribution was calculated as the number of CD8^+^ T cells in each region of brain divided by the total number of CD8^+^ T cells in whole brain.

(**E**) Proportion of each subpopulation of CD8^+^ T cells in the brain, dura, and blood across different ages. Showing every data point of Fig 2.F.

(**F**) Gating strategy demonstrating an alternative method for analysing naive CD8^+^ T cells. CD8^+^ T cells were gated first and then further gated based on intravascular stained (prior to sacrifice) CD45 expression.

(**G**) Proportion of Tnaive cells within tissue CD8^+^ T cells using gating strategy #1 (standard) and #2.

(**H**) CD8^+^ T cells isolated from the spleen and lymph nodes (LNs) of CD45.1 mice were transferred to recipient mice aged 0.5-0.75 mo or 3 mo. After 1.5 months post-adoptive transfer, the repopulation efficiency was analyzed by flow cytometry.

(**I** and **J**) Proportion of CD45.1^+^ CD8^+^ T cells among CD45^hi^ immune cells from brain and dura were compared between recipient mice those got transfer on their age 0.5-0.75 mo and 3mo.

(**K**) Proportion of Ki-67^+^ CD8^+^ T cells were plotted. All data were collected through flow cytometry.

(**L** and **M**) Relationship between mouse age and the rate of change in CD8^+^ T cell number per unit age for Brain (**L**), and dura (**M**). The horizontal line shows the average change in cell number divided by the change in age. The x error bars indicate the range of ages over which the change in cell number was calculated. The y error bars represent the coefficient of variation, calculated as the sum of the variances of the cell number for both age points in each interval.

*P<0.05, **P<0.01, calculated by student t test (I and J) and One-way ANOVA with Dunnett’s multiple comparisons test (A and K), linear regression (B and E), and Two-way ANOVA with Dunnett’s multiple comparisons test (D). Data are shown as the mean ±SEM.

**Figure S7. Microglia- or astrocyte-specific β-2m depletion didn’t affect peripheral immunity, while PLX treatment seems to activate CD8^+^ T cells in spleen and blood, related to Figure 3.**

(**A**) ISI (Infection Similarity Index) of normal blood and brain CD8^+^ T cells from 0.5 mo and 2 mo old male mice with spleen CD8^+^ T cells from 0.5 mo old male non-infected and Influenza-infected mice^35^.

(**B**) Proportions of rare clonal CD8^+^ T cells determined from TCR sequencing of the brain, mesenteric lymph nodes (mLN), lung, liver, and large intestine (L.I.) from 2 mo-old male B6 mice (n=1, pooled from 2 mice).

(**C**) H2-D1 RNA expression profiles across age in brain cell types, including microglia^36^, astrocytes, oligoddendroytes, and endothelial cells^37^.

(**D-I**) Total number of CD8^+^ T cells (D), each number of naive, Tcm, Tem and etc of CD8^+^ T cells (E), each number of CD11b^+^ myeloid cells, NK cells, B cells and CD4^+^ T cells (F) in the brain from control (Veh) and PLX treated mice was shown. The proportion of CD8^+^ T cells among CD45^hi^ cells (G) and composition of naive, Tcm, Tem, etc among CD8^+^ T cells (H and I) in the spleen from control (Veh) and PLX treated mice was shown. All data were collected through flow cytometry.

(**J** - **O**) Histogram of MHCI expression of microglia from control and Tmem119 creERT2 –b2m conditional KO mouse (I). Each number of naive, Tcm, Tem and etc of CD8^+^ T cells (K), each number of CD11b^+^ myeloid cells, NK cells, B cells and CD4 T cells (L) in the brain from control (Cre^-^) and cKO mice (Cre^+^) was shown. The proportion of CD8^+^ T cells among CD45^hi^ cells (M) and composition of naive, Tcm, Tem, etc among CD8^+^ T cells (N and O) in the spleen from control (Cre^-^) and cKO mice (Cre^+^) was shown. All data were collected through flow cytometry.

(**P** - **U**) Histogram of MHCI expression of astrocytes from control and GFAP creERT2–b2m conditional KO mouse (P). Each number of naive, Tcm, Tem and etc of CD8^+^ T cells (Q), each number of CD11b^+^ myeloid cells, NK cells, B cells and CD4^+^ T cells (R) in the brain from control (Cre^-^) and cKO mice (Cre^+^) was shown. The proportion of CD8^+^ T cells among CD45^hi^ cells (S) and composition of naive, Tcm, Tem, etc among CD8^+^ T cells (T and U) in the spleen from control (Cre^-^) and cKO mice (Cre^+^) was shown. All data were collected through flow cytometry.

(**V** – **AA**) Histogram of MHCI expression of brain endothelial cells from control and Tek cre –b2m KO mouse (V). Each number of naive, Tcm, Tem and etc of CD8^+^ T cells (W), each number of CD11b^+^ myeloid cells, NK cells, B cells and CD4^+^ T cells (X) in the brain from control (Cre^-^) and KO mice (Cre^+^) was shown. The proportion of CD8^+^ T cells among CD45^hi^ cells (Y) and composition of naive, Tcm, Tem, etc among CD8^+^ T cells (Z and AA) in the spleen from control (Cre^-^) and KO mice (Cre^+^) was shown. All data were collected through flow cytometry.

**P<0.01, calculated by Wilcoxon rank-sum test with Bonferroni correction (C), student t test (D, G, M, S, and Y), and Two-way ANOVA with Sidak’s multiple comparisons tests (E, F, H, I, K, L, N, O, Q, R, T, U, W, X, Z, and AA). Data are shown as the mean ±SEM.

**Figure S8. The absence of brain-specific CD8+ T cells is associated with abnormal behaviors, related to Figure 5.**

(**A-D**) Relative and absolute distribution of brain CD8^+^ T cells (**A** and **B**) and CD4^+^ T cells (**C** and **D**) in 3-mo-old WT and CD8 KO mice.

(**E**) Relative distribution of each immune cell in the spleen of 3-mo-old WT and CD8 KO mice.

(**F**-**K**) Developmental milestones were examined in pups of both WT and CD8 KO from postnatal days 4 to 16. Behavioral assessments included eye-opening (**F**), tactile startle reflex (**G**), grasp reflex (**H**), cliff avoidance (**I**), negative geotaxis (**J**), and surface righting (**K**). No significant differences were observed between CD8 KO mice and WT mice.

(**L**-**O**) Adult cognitive behaviors, including anxiety-like behavior, social interaction, and recognition, were evaluated in adult mice of WT and CD8 KO strains. Anxiety-like behaviors were measured using an open field test (**L**) and Elevated Zero Maze test (**M**). Social interaction was assessed through a three-chamber test (**N**). Recognition was tested using a novel objective test (**O**).

(P and Q) Social interaction (P) and Anxiety-like (Q) behaviors with WT, TMEM119-creERT2 dependent β2m conditional knock-out mice (from Figure 3), and 4wk-Abx (from Figure 4) mice.

**P<0.01, calculated by student t-test (A-D, L-O, and R-T), two-way ANOVA with Sidak’s multiple comparisons test (E), two-way RM ANOVA with Sidak’s multiple comparisons test (F-K). one-way ANOVA with Dunnett (P and Q). Data are shown as the mean ±SEM.

**Figure S9 Depletion of brain-specific CD8^+^ T cells in adult mice is not associated with abnormal behaviors.**

(A-H) A CD8 depletion model was generated by i.p injecting an isotype control antibody or anti-CD8 antibody (clone 2.43) to wildtype C57BL/6 mice from 3 wks to 8 wks according to the injection protocol every 3-5 days. Proportions of dendritic cells (A), Macrophages (B), Monocytes (C), Neutrophils (D), NK cells (E), B cells (F), γδT cells (G) in the brain and spleen of control and CD8-depleted mice were shown. Total number difference of CD8^+^ T cells in the brain (**H**) according to the CD8^+^ T cell depletion are shown (Iso n=6; α-CD8, n=6 from 3 independent experiments).

(**I-P**) A CD8 repopulation model was generated by injecting depleting antibody for 5 wks from 3wks of mice age followed by 4 wks of antibody withdrawal (Iso-Re and α-CD8-Re). Proportions of dendritic cells (**I**), Macrophages (**J**), Monocytes (**K**), Neutrophils (**L**), NK cells (**M**), B cells (**N**), γδT cells (**O**) in the brain and spleen of control and CD8-depleted mice were shown. Total number CD8^+^ T cells in the brain (P) according to the withdrawal of CD8^+^ T cell-depletion are shown (Iso n=4; α-CD8, n=4 from 2 independent experiments).

(Q-U) Adult CD8 depletion model was generated by i.p injecting an isotype control antibody or anti-CD8 antibody (clone 2.43) to wildtype C57BL/6 mice from 9 wks to 13 wks according to the injection protocol every 3-5 days (Q). Representative FACS plots of T cells (Livedead^-^, CD45^hi^TCRβ^+^) were shown (R).

Proportions of CD8^+^ T cells (**C**) in the brain and spleen from control (Iso-AD) and CD8^+^ T cells depleted mice (α-CD8-AD) are shown (Iso-AD n=2; α-CD8-AD, n=2 from 1 independent experiments).

(**T** and **U**) Social interaction (**T**) and anxiety-like (**U**) tests were conducted with Iso-AD and α-CD8-AD mice (Iso-AD n=3; α-CD8-AD, n=4 from 2 independent experiments).

**Figure S10**

(**A**) Representative images (left) and quantification (right) of cFos-positive neurons in the BLA (Iso n=4, α-CD8 n=4), hippocampus (Iso n=4, α-CD8 n=4), M1 (Iso n=5, α-CD8 n=4), S1 (Iso n=4, α-CD8 n=4), and ECT (Iso n=3, α-CD8 n=4) of Iso and α-CD8 mice after 10 min of social interaction.

(**B)** Number of cFos-negative cells among NeuN-positive cells, related to Fig 5H and I.

(**C**-**L**) Microglial morphological analysis in Iso and α-CD8 mice after 10 min of social interaction. Schematics for the isolation of single microglia in a photomicrograph (C). The quantification of cell body size (D), number of branches, average length of branches (E), number of junctions (F), length of longest-shortest path (G), endpoints per microglia (H), circularity (I), fractal dimension (J), density (K), and span ratio (L) analyzed at a single cell level.

(**M)** Schematic illustration of the experimental workflow for bulk RNA sequencing. Microglia were isolated from mice injected with anti-CD8α antibody (α-CD8) or IgG2b isotype control (Iso).

(**N)** Volcano plot showing differentially expressed genes (false discovery rate (FDR) < 0.05 and log2 fold change > 0.5) between microglia from Iso- and α-CD8 mice. Genes upregulated in α- CD8 are highlighted in green.

(**O)** Box plots showing log2 fold changes of HOM, M1, M2, and DAM signature genes (gene list in Supplementary Table S5).

(**R**-**U**) (R) Representative confocal images of the mPFC of inhibitory synapses identified by the co-localization between vGAT and gephyrin (white arrows) in Iso and α -CD8 mice. Scale bars = 2um. (S) The number of inhibitory synapses between Iso and α -CD8 mice. The coverage ratio of Vgat signal (T) and Gephyrin signal (U) in the photomicrographs.

**Figure S11 Brain CD8^+^ T cells exhibit distinct characteristics, related to Figure 6**

(**A**) Volcano plots for differentially expressed between male-brain CD8^+^ T cells and female-brain CD8^+^ T cells. For genes enriched in male CD8^+^ T cells are highlighted in blue, and those enriched in female are highlighted in dark blue.

(**B-E**) Representative FACS plot and its quantification data of adult male CD8^+^ T cells from noted organs

(**B**). Brain and splenic CD8^+^ T cells from LCMV Armstrong infected, post-infection day 8 of adult male mice were used. GranzymeB-BV421 and Perforin-APC fluorophore-conjugated antibodies were used for staining.

(**F and G**) Representative histogram and its MFI value of CD107a expression of brain and spleen CD8^+^ T cells measured by flow cytometry (**F**). Quantification of MFI value of CD107a expression of brain and spleen CD8^+^ T cells(**G**).

(**H-K**) Representative FACS plot (**H**) and its quantification data (**I-K**) of intracellular IFN-γ and TNFα production of the adult male brain, blood, and spleen CD8^+^ T cells after 4 hr ex-vivo stimulation with PMA and Ionomycin in the presence of brefeldin A. Unstimulated splenic CD8^+^ T cells were used as negative control. TNFα-APC and IFN-γ-PE fluorophore-conjugated antibodies were used for the FACS plots.

(**L**) Proportion of IFNγ^+^ in CD8^+^T, CD4^+^T, gdT, NK, and CD45^hi^ Lineage^-^ cells from brain, dura, spleen, blood, kidney, and liver. Total CD45^hi^ cells were stimulated with PMA and Ionomycin in the presence of brefeldin A for 4 hours, followed by flow cytometry analysis.

(**M** and **N**) In vivo IFN-γ ^+^ CD8^+^ T cells were counted in the mouse brain. (IFNγ : Green, CD8α: Red) Scare bar: 10um.

(**O**) IFNγ protein amount per brain was determined by Immunoassay system (Invitrogen, ProQuantum Immunoassays). Brain lysate from wild-type mice, PBS (CD8 KO mice group), WT-CD8 (CD8 KO mice adoptively transferred with WT CD8^+^ T cells), and Ifng KO-CD8 (CD8KO mice adoptively transferred with Ifnγ-/- CD8^+^ T cells) described in Figure 7 was used for measuring IFNγ protein level.

(**P** and **Q**) Representative histogram of CD69 expression of brain and spleen CD8^+^ T cells (**P**). Quantification of MFI value of CD69 expression of brain and spleen CD8^+^ T cells(**Q**).

(**R and S**) Representative histogram of PD-1 expression and its proportion in brain, spleen CD8^+^ T cells and P14 brain CD8^+^ T cells (post infection day 40 of LCMV-clone 13-infected mice) (R). Quantification of the proportion of PD-1^+^ cell in brain, spleen and LCMV clone 13-infected brain P14 CD8 T cells (S). (**T and U**) Representative histogram showing the proportion of exhaustion-related inhibitory receptors (2B4, TIGIT, CD160, Tim3, Lag3) and Tox expression in the brain and spleen CD8^+^ T cells (**T**). Quantification of the proportion of inhibitory receptors and Tox expression of brain and spleen CD8^+^ T cells (**Q**).

(**V**) Schematic illustration of CD8^+^ T cell differentiation. Naïve CD8^+^ T cells differentiate into either memory precursor (MP) or effector cells. Memory and exhausted CD8^+^ T cells subsequently arise from the MP population.

(**W**) UMAP embeddings of combined CD8^+^ T cell subsets from brain and blood datasets with spleen-derived CD8^+^ T cells from LCMV-infected mice^54^. LeftL red dots represent blood, blue dots represent brain, and gry dots represent LCMV CD8^+^ T cells. Right: colors indicate cell type annotation of the CD8^+^ T cells.

(**X**) Violine plots showing expression levels of selected marker genes (*Gzma, Gzmb, Prf1, Ifng, Tnf,* and *Pdcd1*).

**Figure S12. Adaptive transferred CD8^+^ T cells eliminate the deficits in SCID mice, related to Figure 7.**

(**A**) Schematic diagram of CD8^+^ T cell transfer experiment to SCID mice. CD8^+^ T cells were isolated from spleen and LNs of WT donor mice. After 6 weeks from T cell transfer, SCID mice were used for behavioral examinations.

(**B**) Relative distribution of CD8^+^ T cells in the brain, dura, spleen, and blood of CD8^+^ T cells transferred SCID mice.

(**C**) Absolute number of CD8^+^ T cells in the brain and dura from SCID recipient mice.

(**D** and **E**) Social interaction (**D**) and anxiety-like (**E**) tests were conducted on PBS-SCID and CD8-SCID mice.

(**F**) Mating strategy to generate Tmem119-creERT2 dependent IFNγR1 conditional knock-out mice.

(**G and H**) Social interaction (**G**) and anxiety-like (**H**) tests were conducted on control and KO mice. (Cre-, n=5, Cre+, n=5, from 2 independent experiments).

(**I**) Mating strategy to generate GFAP-creERT2 dependent IFNγR1 conditional knock-out mice.

(**J and K**) Social interaction (**J**) and anxiety-like (**K**) tests were conducted on control and KO mice. (Cre-, n=3, Cre+, n=3, from 2 independent experiments).

(**L**) The proportion of CD8^+^ T cells among CD45^hi^ immune cells (CD8^+^ T /CD45^hi^ cells) was analyzed in the human brain using CIBERSORTx. The human brain exhibited a positive correlation between age and the proportion of CD8^+^ T cells. Raw data were obtained from PsychENCODE RNA-seq data.

(**M**) CD8^+^ T /CD45^hi^ cells were analyzed in the human ASD, schizophrenia, and bipolar disorder patient’s brain using CIBERSORTx (Voineagu, 2011, Cont (n=6), ASD (n=6); Xiao, 2014, Cont (n=11), Schizo (n=10), BD (n=14)).

*P<0.05, **P<0.01, calculated by an One-way ANOVA (B and M), student’s t test (C, D, E, G-H, J-K and M), and Pearson’s correlation test (L). Data are shown as the mean ±SEM.

## Notes

### Competing Interest Statement

The authors have declared no competing interest.

### Summary of Updates

we updated the funding information, so it can be searchable at NCBI.

