## supplementary figures for "Brain-Resident CD8^+^ T Cells Regulate Neuronal Activity and Behavior via Interferon-Gamma"

Figure S1

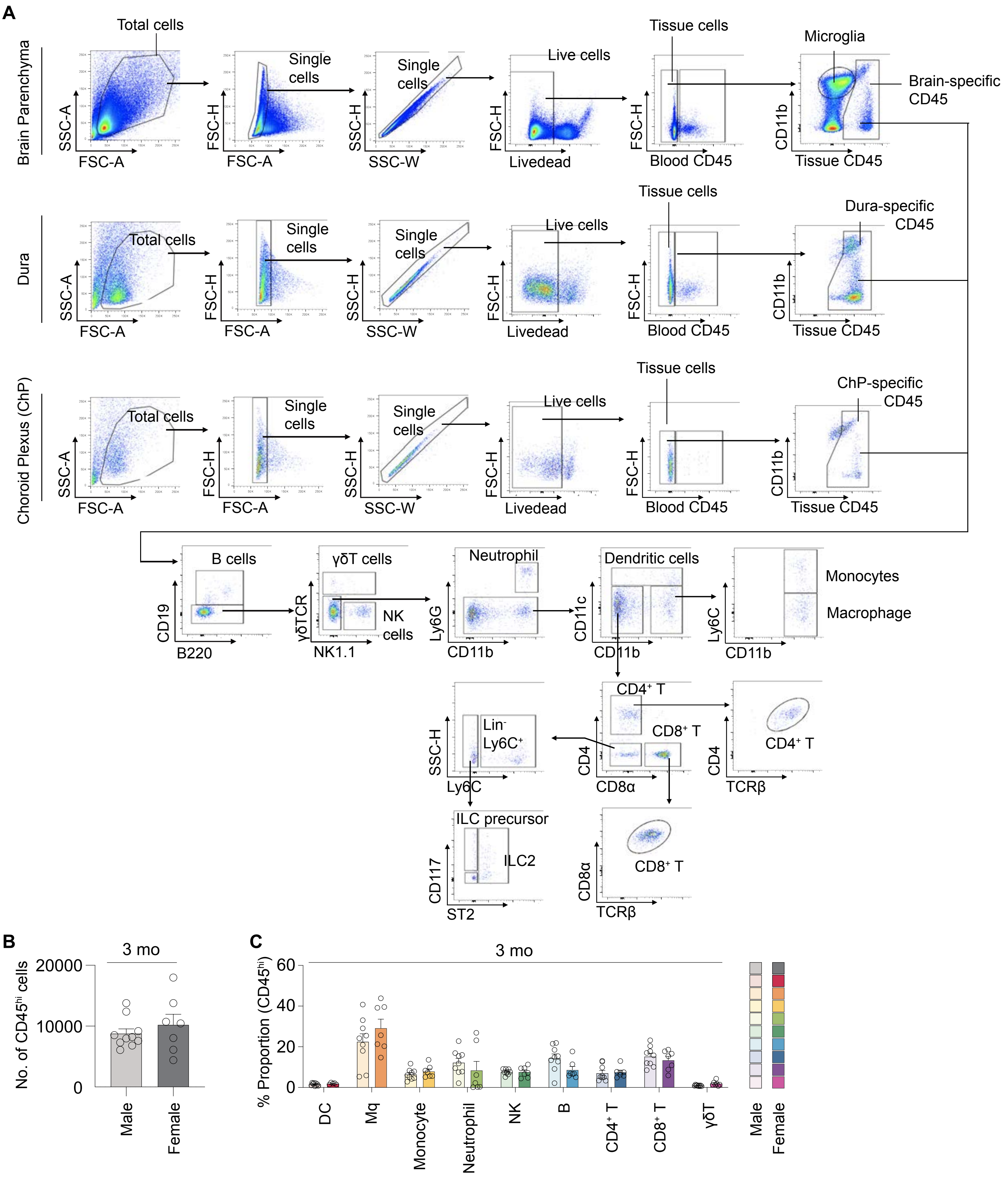
A
Brain Parenchyma
Total cells
SSC-A
FSC-A
Single cells
FSC-H
FSC-A
Single cells
SSC-H
SSC-W
Live cells
FSC-H
Livedead
Tissue cells
FSC-H
Blood CD45
Microglia
Brain-specific CD45
CD11b
Tissue CD45
Dura
Total cells
SSC-A
FSC-A
Single cells
FSC-H
FSC-A
Single cells
SSC-H
SSC-W
Live cells
FSC-H
Livedead
Tissue cells
FSC-H
Blood CD45
Dura-specific CD45
CD11b
Tissue CD45
Choroid Plexus (ChP)
Total cells
SSC-A
FSC-A
Single cells
FSC-H
FSC-A
Single cells
SSC-H
SSC-W
Live cells
FSC-H
Livedead
Tissue cells
FSC-H
Blood CD45
ChP-specific CD45
CD11b
Tissue CD45
B cells
CD19
B220
γδT cells
γδTCR
NK cells
NK1.1
Neutrophil
Ly6G
CD11b
Dendritic cells
CD11c
CD11b
Monocytes
Macrophage
Ly6C
CD11b
Lin⁻ Ly6C⁺
SSC-H
Ly6C
CD4⁺ T
CD8⁺ T
CD4
CD8α
CD4⁺ T
CD4
TCRβ
ILC precursor
ILC2
CD117
ST2
CD8⁺ T
CD8α
TCRβ
B
3 mo
No. of CD45hi cells
20000
10000
0
Male
Female
C
3 mo
% Proportion (CD45hi)
60
40
20
0
DC
Mq
Monocyte
Neutrophil
NK
B
CD4⁺ T
CD8⁺ T
γδT
Male
Female

### Figure S2

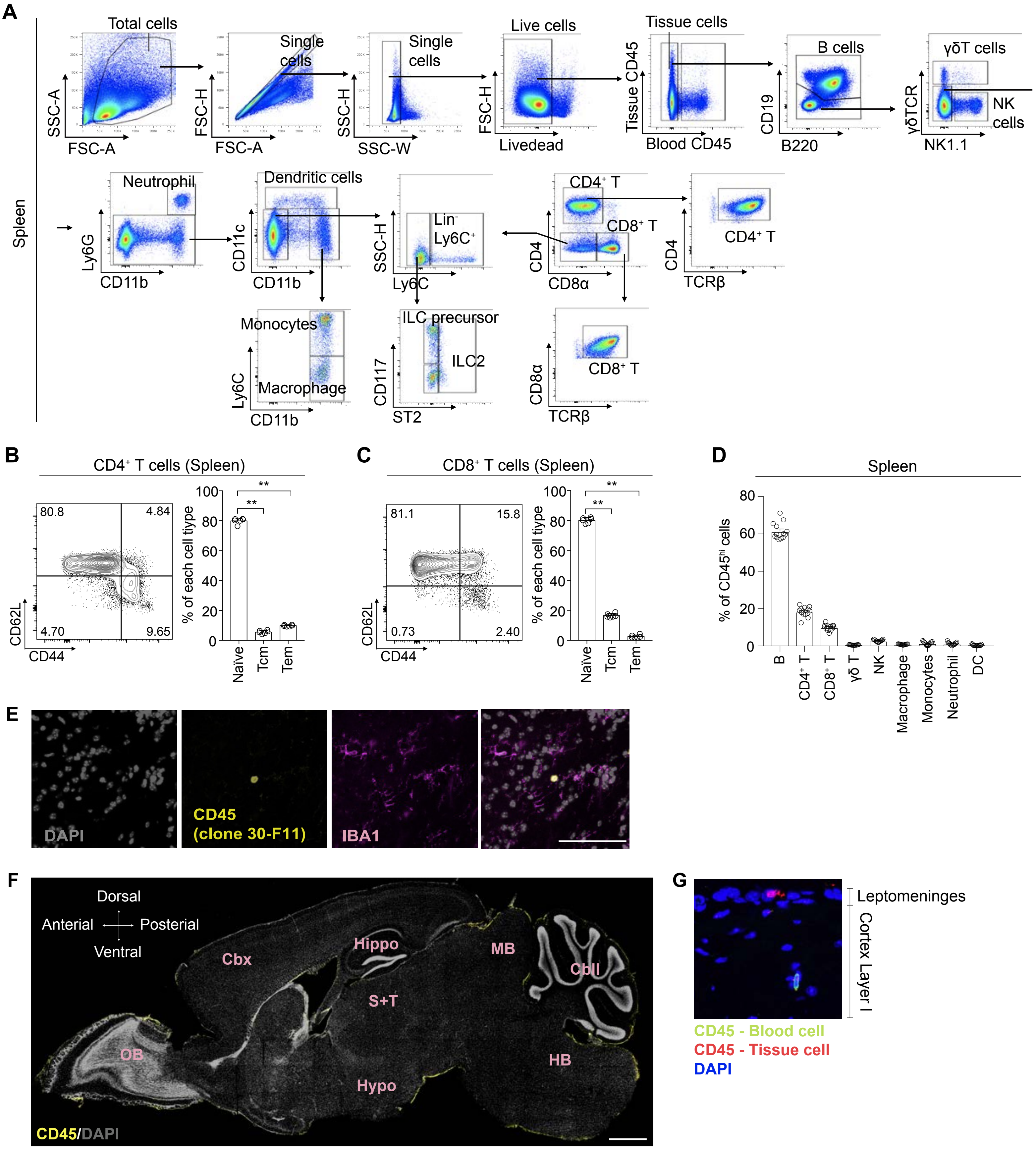
A
Spleen
Total cells
Single cells
Single cells
Live cells
Tissue cells
B cells
γδT cells
NK cells
SSC-A
FSC-A
FSC-H
FSC-A
SSC-H
SSC-W
FSC-H
Livedead
Tissue CD45
Blood CD45
CD19
B220
γδTCR
NK1.1
Neutrophil
Dendritic cells
Lin⁻ Ly6C⁺
CD4⁺ T
CD8⁺ T
CD4⁺ T
Ly6G
CD11b
CD11c
CD11b
SSC-H
Ly6C
CD4
CD8α
CD4
TCRβ
Monocytes
Macrophage
Ly6C
CD11b
ILC precursor
ILC2
CD117
ST2
CD8⁺ T
CD8α
TCRβ
B
CD4⁺ T cells (Spleen)
80.8
4.84
4.70
9.65
CD62L
CD44
% of each cell type
Naïve
Tcm
Tem
C
CD8⁺ T cells (Spleen)
81.1
15.8
0.73
2.40
CD62L
CD44
% of each cell type
Naïve
Tcm
Tem
D
Spleen
% of CD45hi cells
B
CD4⁺ T
CD8⁺ T
γδ T
NK
Macrophage
Monocytes
Neutrophil
DC
E
DAPI
CD45 (clone 30-F11)
IBA1
F
Dorsal
Anterial
Posterial
Ventral
Cbx
Hippo
MB
Cbll
S+T
OB
HB
Hypo
CD45/DAPI
G
Leptomeninges
Cortex Layer I
CD45 - Blood cell
CD45 - Tissue cell
DAPI

### Figure S3

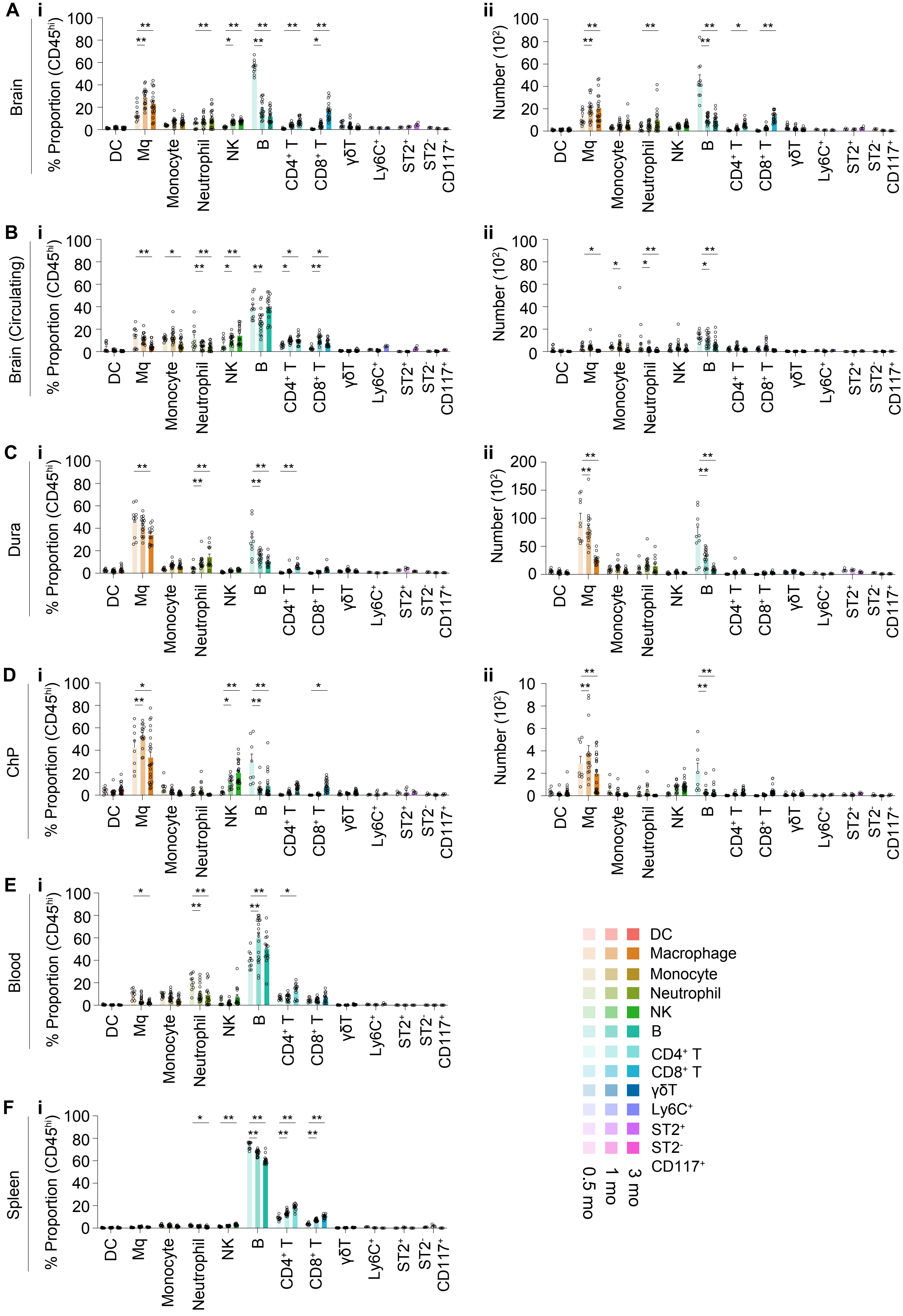

A
B
C
D
E
F
i
ii
Brain
Brain (Circulating)
Dura
ChP
Blood
Spleen
% Proportion (CD45hi)
Number (10²)
DC
Mq
Monocyte
Neutrophil
NK
B
CD4⁺ T
CD8⁺ T
γδT
Ly6C⁺
ST2⁺
ST2⁻ CD117⁺
DC
Macrophage
Monocyte
Neutrophil
NK
B
CD4⁺ T
CD8⁺ T
γδT
Ly6C⁺
ST2⁺
ST2⁻ CD117⁺
0.5 mo
1 mo
3 mo

### Figure S4

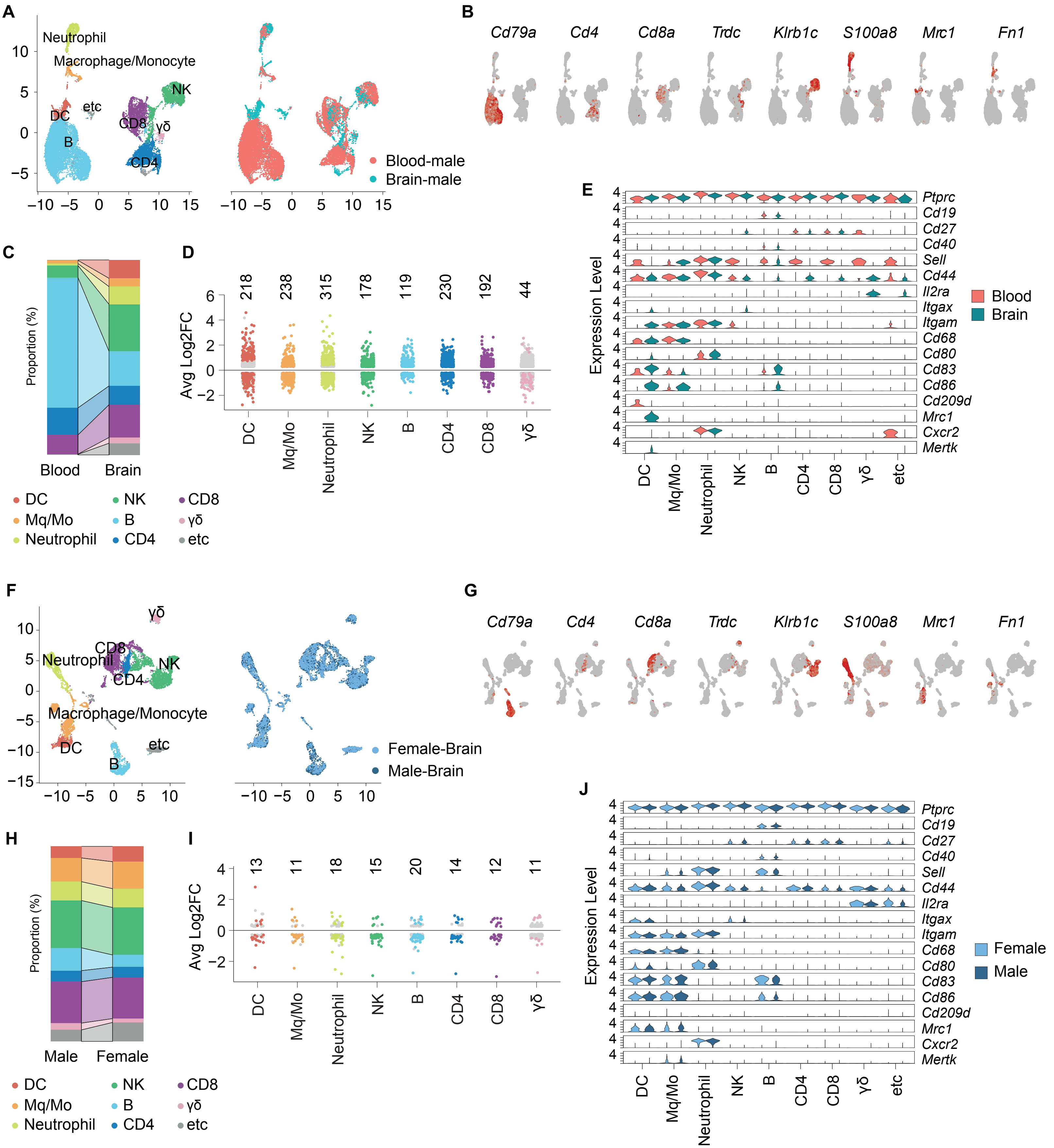

A
Neutrophil
Macrophage/Monocyte
NK
DC
etc
CD8
γδ
B
CD4
Blood-male
Brain-male
B
Cd79a
Cd4
Cd8a
Trdc
Klrb1c
S100a8
Mrc1
Fn1
C
Proportion (%)
Blood
Brain
DC
Mq/Mo
Neutrophil
NK
B
CD4
CD8
γδ
etc
D
Avg Log2FC
218
238
315
178
119
230
192
44
E
Expression Level
Ptprc
Cd19
Cd27
Cd40
Sell
Cd44
Il2ra
Itgax
Itgam
Cd68
Cd80
Cd83
Cd86
Cd209d
Mrc1
Cxcr2
Mertk
Blood
Brain
F
Female-Brain
Male-Brain
G
H
Male
Female
I
13
11
18
15
20
14
12
11
J
Female
Male

Figure S5

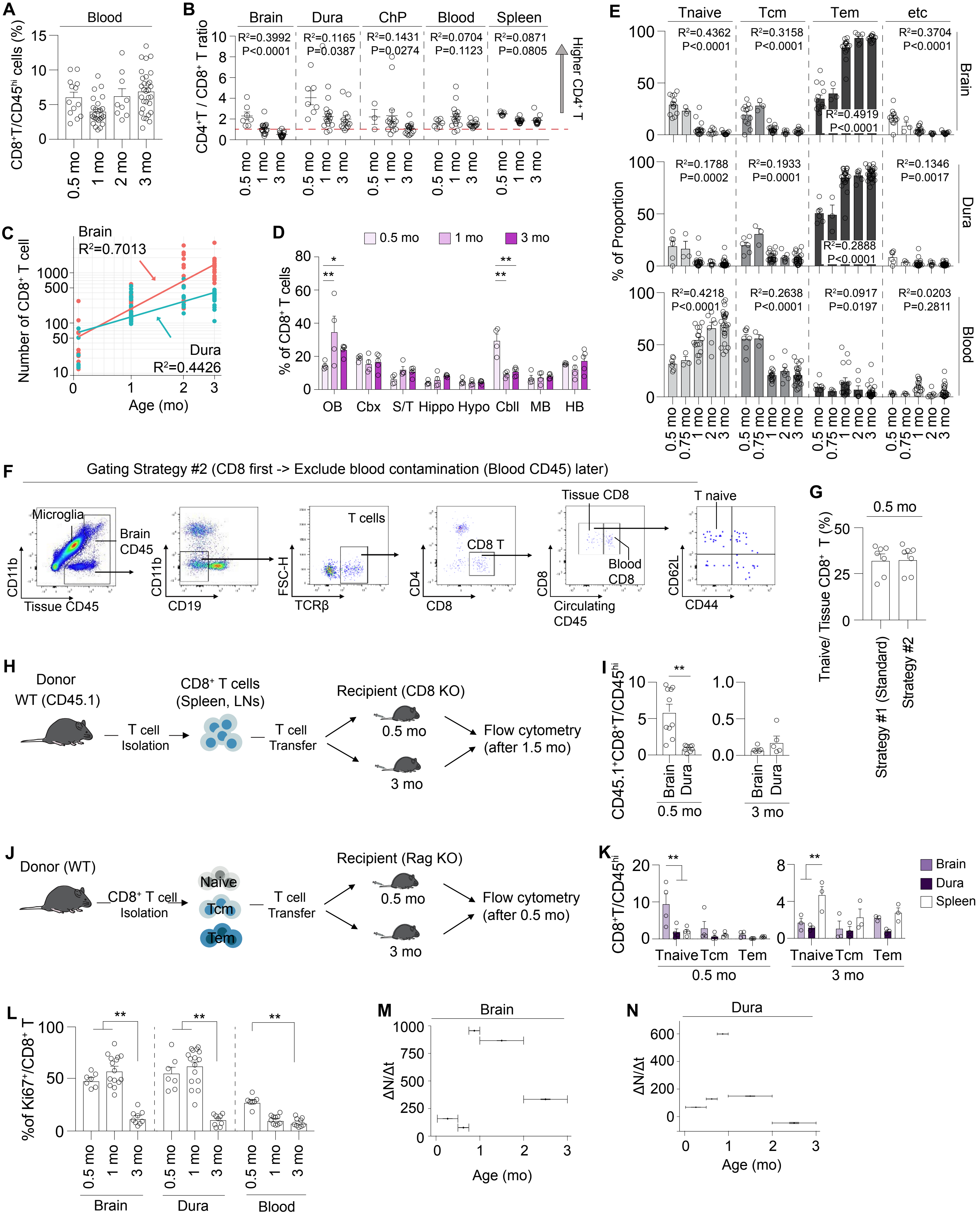

A
Blood
CD8+T/CD45hi cells (%)
0.5 mo
1 mo
2 mo
3 mo
B
Brain
Dura
ChP
Blood
Spleen
CD4+T / CD8+ T ratio
R²=0.3992 P<0.0001
R²=0.1165 P=0.0387
R²=0.1431 P=0.0274
R²=0.0704 P=0.1123
R²=0.0871 P=0.0805
Higher CD4+ T
C
Brain
R²=0.7013
Dura
R²=0.4426
Number of CD8+ T cell
Age (mo)
D
0.5 mo
1 mo
3 mo
% of CD8+ T cells
OB
Cbx
S/T
Hippo
Hypo
Cbll
MB
HB
E
Tnaive
Tcm
Tem
etc
% of Proportion
Brain
Dura
Blood
R²=0.4362 P<0.0001
R²=0.3158 P<0.0001
R²=0.4919 P<0.0001
R²=0.3704 P<0.0001
R²=0.1788 P=0.0002
R²=0.1933 P=0.0001
R²=0.2888 P<0.0001
R²=0.1346 P=0.0017
R²=0.4218 P<0.0001
R²=0.2638 P<0.0001
R²=0.0917 P=0.0197
R²=0.0203 P=0.2811
0.5 mo
0.75 mo
1 mo
2 mo
3 mo
F
Gating Strategy #2 (CD8 first -> Exclude blood contamination (Blood CD45) later)
Microglia
Brain CD45
CD11b
Tissue CD45
CD19
T cells
FSC-H
TCRβ
CD8 T
CD4
CD8
Tissue CD8
Blood CD8
Circulating CD45
T naive
CD62L
CD44
G
0.5 mo
Tnaive/ Tissue CD8+ T (%)
Strategy #1 (Standard)
Strategy #2
H
Donor WT (CD45.1)
T cell Isolation
CD8+ T cells (Spleen, LNs)
T cell Transfer
Recipient (CD8 KO)
0.5 mo
3 mo
Flow cytometry (after 1.5 mo)
I
CD45.1+CD8+T/CD45hi
Brain
Dura
0.5 mo
3 mo
J
Donor (WT)
CD8+ T cell Isolation
Naive
Tcm
Tem
T cell Transfer
Recipient (Rag KO)
0.5 mo
3 mo
Flow cytometry (after 0.5 mo)
K
CD8+T/CD45hi
Tnaive
Tcm
Tem
0.5 mo
3 mo
Brain
Dura
Spleen
L
%of Ki67+/CD8+ T
0.5 mo
1 mo
3 mo
Brain
Dura
Blood
M
Brain
ΔN/Δt
Age (mo)
N
Dura
ΔN/Δt
Age (mo)

### Figure S6

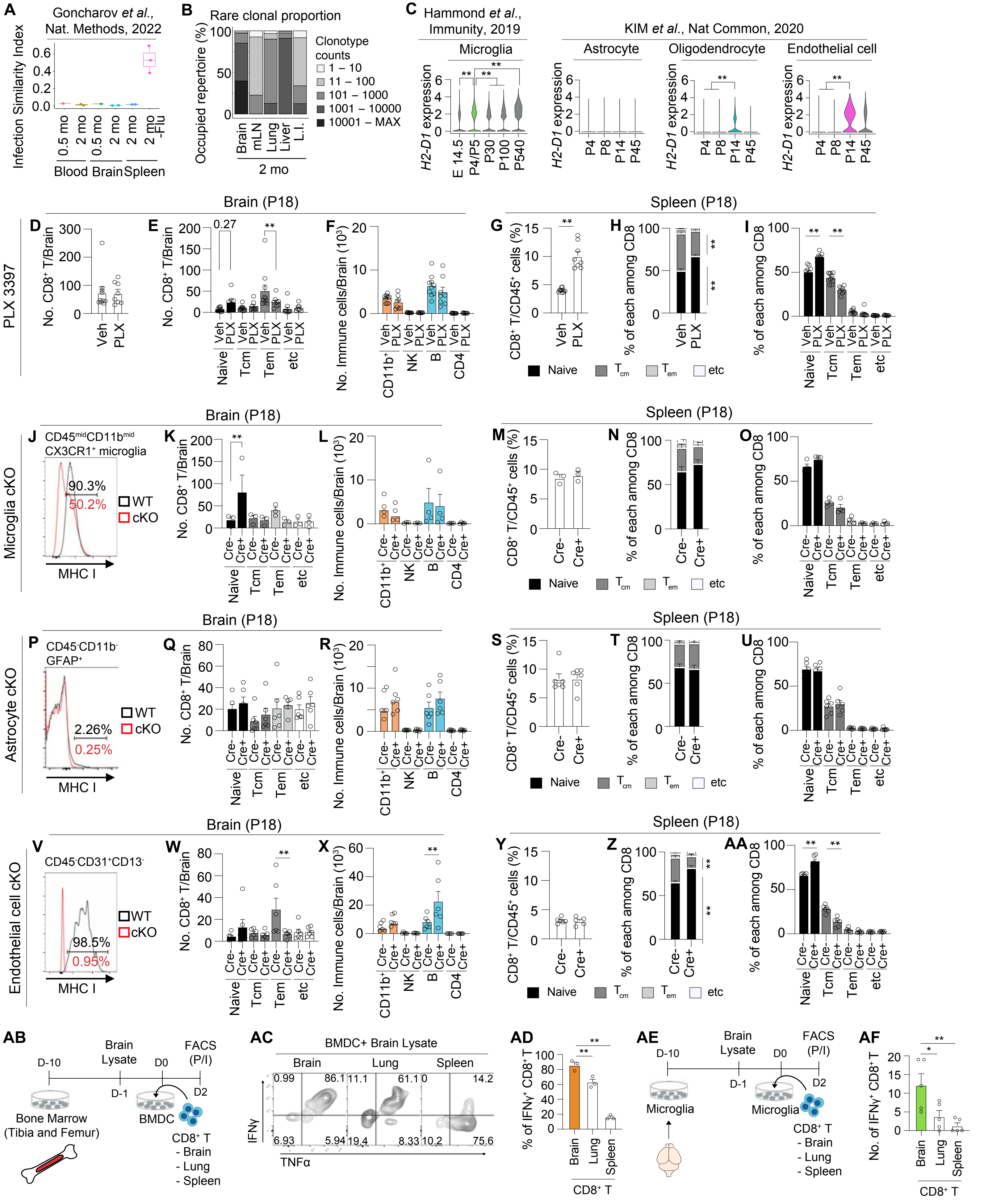
A
Goncharov et al., Nat. Methods, 2022
Infection Similarity Index
Blood Brain Spleen
B
Rare clonal proportion
Occupied repertoire (%)
Clonotype counts
1 – 10
11 – 100
101 – 1000
1001 – 10000
10001 – MAX
Brain mLN Lung Liver L.I.
2 mo
C
Hammond et al., Immunity, 2019
Microglia
KIM et al., Nat Commun, 2020
Astrocyte
Oligodendrocyte
Endothelial cell
H2-D1 expression
Brain (P18)
Spleen (P18)
PLX 3397
No. CD8+ T/Brain
No. Immune cells/Brain (10^3)
CD8+ T/CD45+ cells (%)
% of each among CD8
Naive Tcm Tem etc
Microglia cKO
CD45midCD11bmid CX3CR1+ microglia
90.3%
50.2%
WT
cKO
MHC I
Astrocyte cKO
CD45-CD11b- GFAP+
2.26%
0.25%
Endothelial cell cKO
CD45-CD31+CD13-
98.5%
0.95%
AB
D-10
Brain Lysate
D-1
D0
FACS (P/I)
D2
Bone Marrow (Tibia and Femur)
BMDC
CD8+ T
- Brain
- Lung
- Spleen
AC
BMDC+ Brain Lysate
Brain
Lung
Spleen
IFNγ
TNFα
AD
% of IFNγ+ CD8+ T
AE
Microglia
AF
No. of IFNγ+ CD8+ T

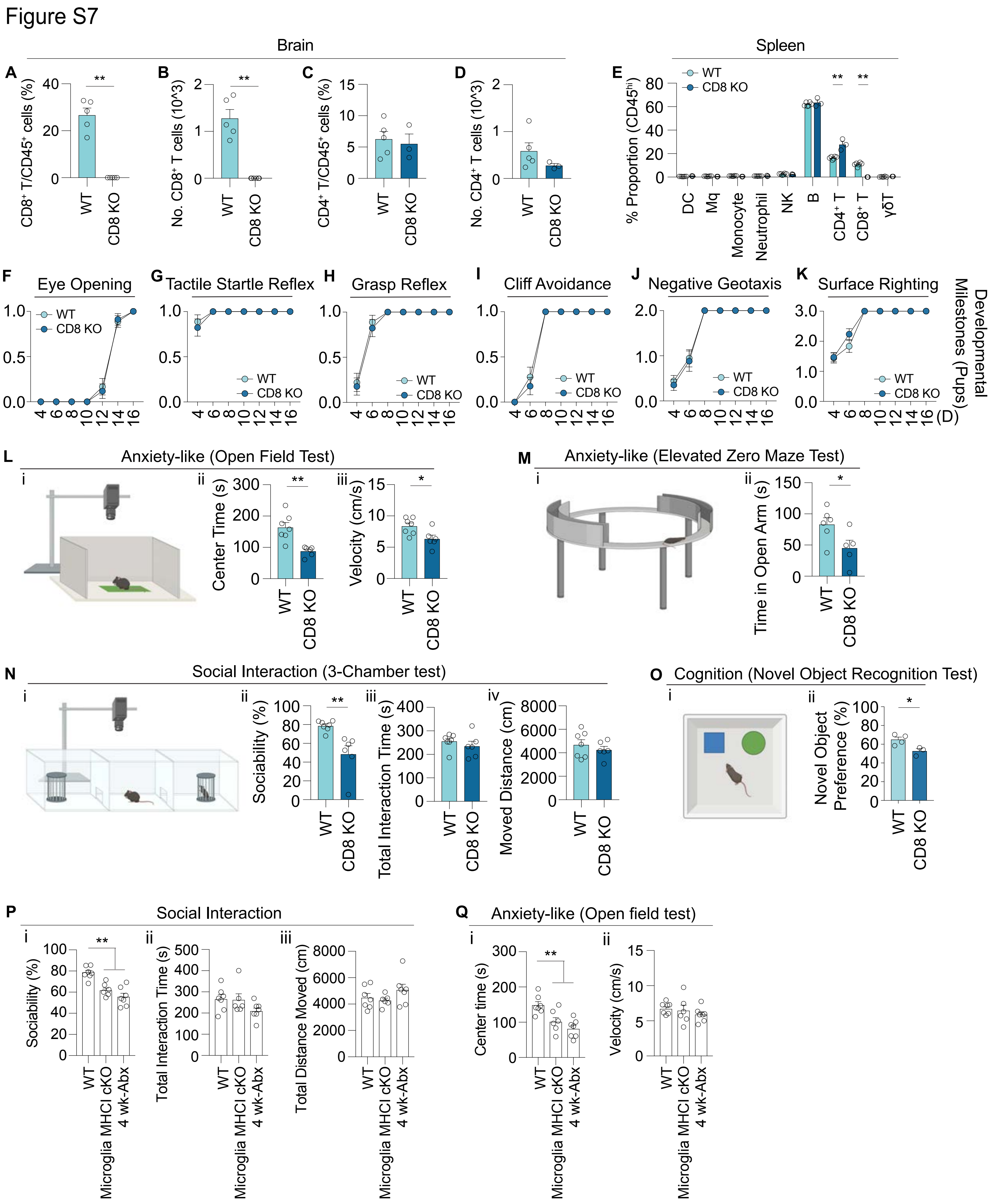
Figure S7
Brain
Spleen
A
CD8⁺ T/CD45⁺ cells (%)
B
No. CD8⁺ T cells (10^3)
C
CD4⁺ T/CD45⁺ cells (%)
D
No. CD4⁺ T cells (10^3)
E
% Proportion (CD45hi)
WT
CD8 KO
DC
Mq
Monocyte
Neutrophil
NK
B
CD4⁺ T
CD8⁺ T
γδT
F Eye Opening
G Tactile Startle Reflex
H Grasp Reflex
I Cliff Avoidance
J Negative Geotaxis
K Surface Righting
Developmental Milestones (Pups)
(D)
L Anxiety-like (Open Field Test)
Center Time (s)
Velocity (cm/s)
M Anxiety-like (Elevated Zero Maze Test)
Time in Open Arm (s)
N Social Interaction (3-Chamber test)
Sociability (%)
Total Interaction Time (s)
Moved Distance (cm)
O Cognition (Novel Object Recognition Test)
Novel Object Preference (%)
P Social Interaction
Sociability (%)
Total Interaction Time (s)
Total Distance Moved (cm)
Microglia MHCI cKO
4 wk-Abx
Q Anxiety-like (Open field test)
Center time (s)
Velocity (cm/s)

### Figure S8

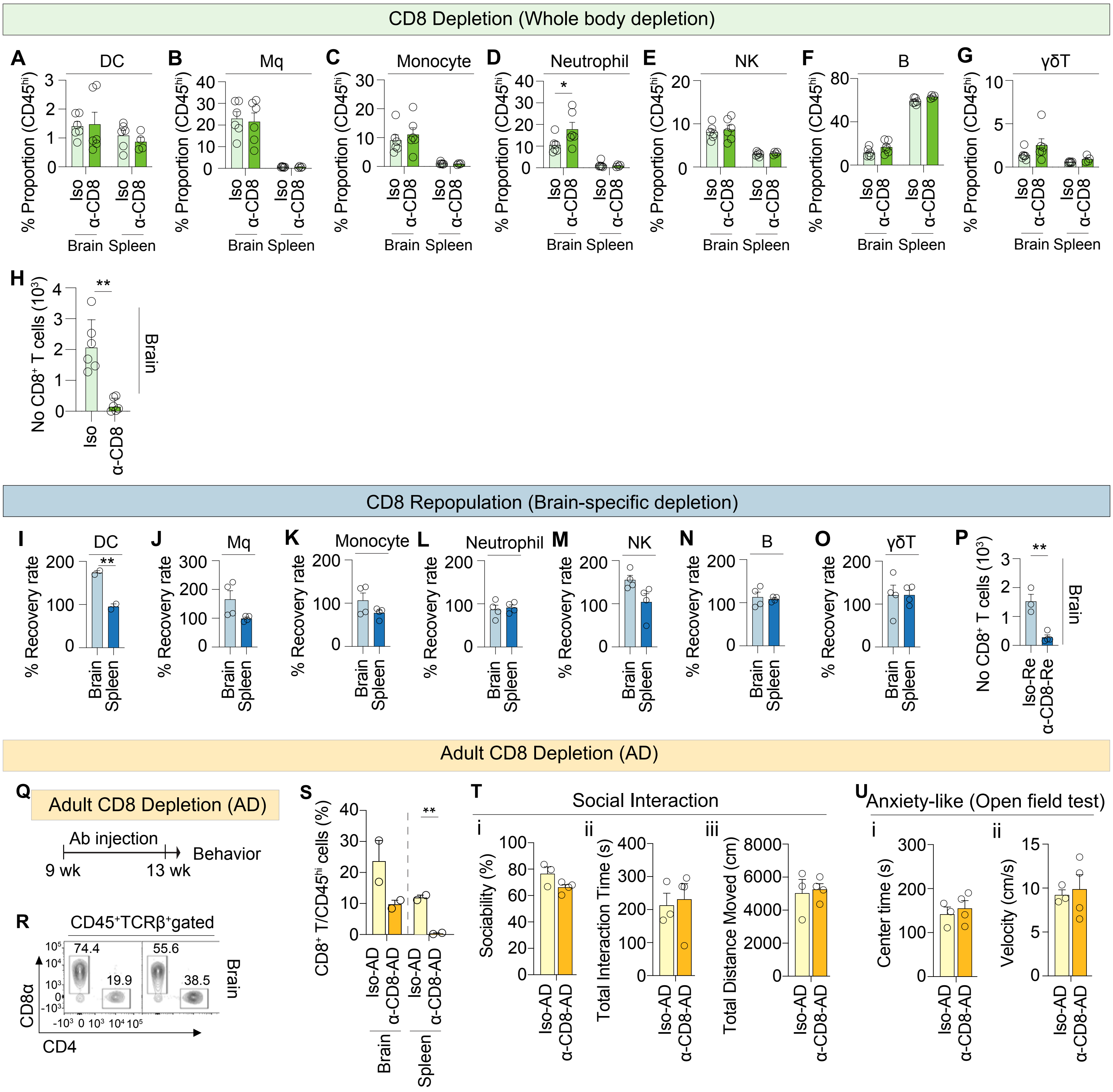

CD8 Depletion (Whole body depletion)
A DC
B Mq
C Monocyte
D Neutrophil
E NK
F B
G γδT
% Proportion (CD45hi)
Iso
α-CD8
Brain Spleen
H
No CD8+ T cells (10³)
Brain
CD8 Repopulation (Brain-specific depletion)
I DC
J Mq
K Monocyte
L Neutrophil
M NK
N B
O γδT
P
% Recovery rate
Brain
Spleen
Iso-Re
α-CD8-Re
Adult CD8 Depletion (AD)
Q
Adult CD8 Depletion (AD)
Ab injection
Behavior
9 wk
13 wk
R
CD45+TCRβ+gated
74.4
19.9
55.6
38.5
CD8α
CD4
S
CD8+ T/CD45hi cells (%)
Iso-AD
α-CD8-AD
T
Social Interaction
i
Sociability (%)
ii
Total Interaction Time (s)
iii
Total Distance Moved (cm)
U
Anxiety-like (Open field test)
i
Center time (s)
ii
Velocity (cm/s)

### Figure S9

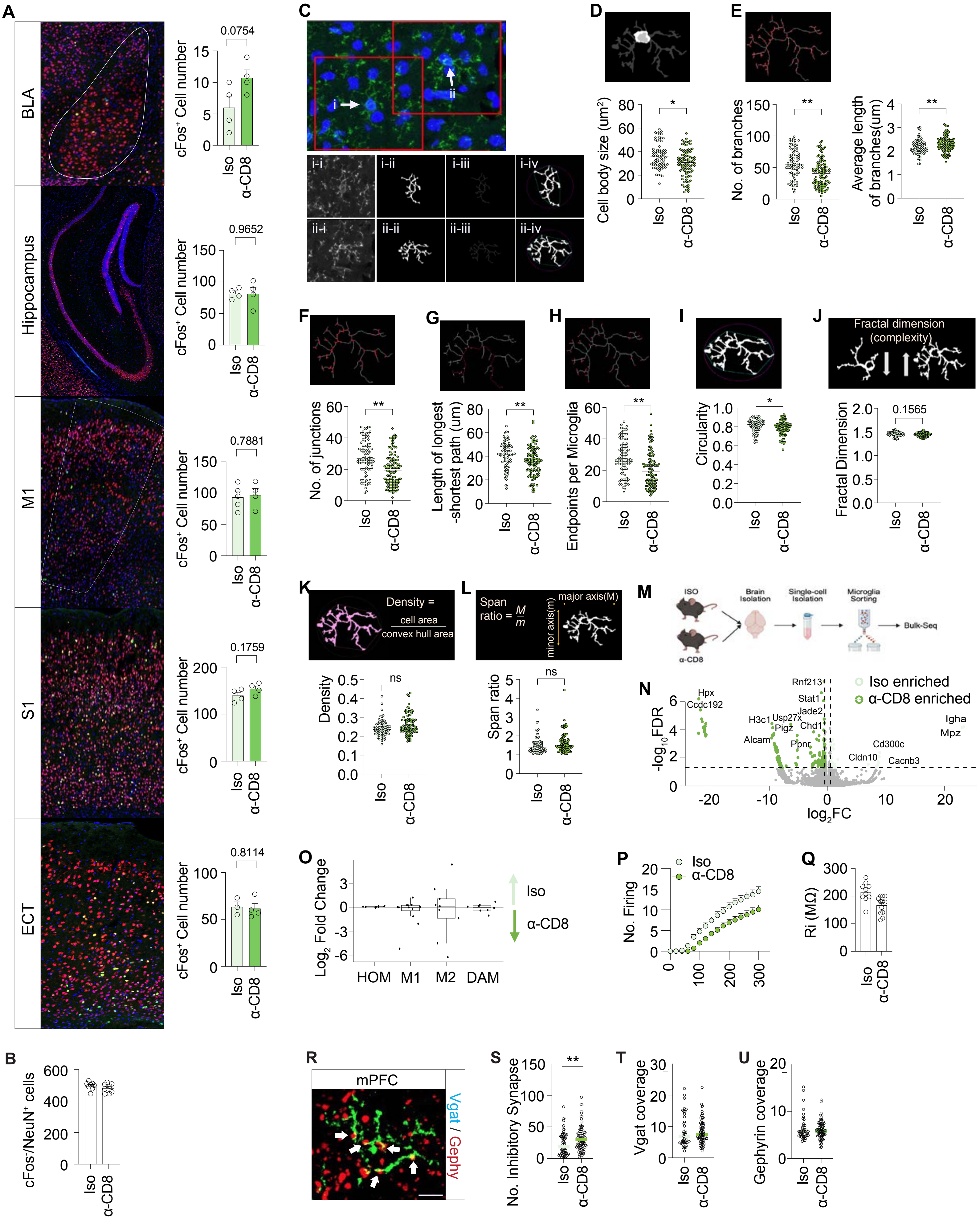
A
BLA
Hippocampus
M1
S1
ECT
cFos+ Cell number
0.0754
0.9652
0.7881
0.1759
0.8114
Iso
α-CD8
B
cFos+/NeuN+ cells
C
i
ii
i-i
i-ii
i-iii
i-iv
ii-i
ii-ii
ii-iii
ii-iv
D
Cell body size (um2)
E
No. of branches
Average length of branches(um)
F
No. of junctions
G
Length of longest -shortest path (um)
H
Endpoints per Microglia
I
Circularity
J
Fractal dimension (complexity)
Fractal Dimension
0.1565
K
Density = cell area / convex hull area
Density
ns
L
Span ratio = M / m
major axis(M)
minor axis(m)
Span ratio
ns
M
ISO
α-CD8
Brain Isolation
Single-cell Isolation
Microglia Sorting
Bulk-Seq
N
Iso enriched
α-CD8 enriched
-log10FDR
log2FC
Hpx
Ccdc192
Rnf213
Stat1
Jade2
H3c1
Usp27x
Pigz
Chd1
Alcam
Ppnr
Igha
Mpz
Cd300c
Cldn10
Cacnb3
O
Log2 Fold Change
HOM
M1
M2
DAM
Iso
α-CD8
P
No. Firing
Iso
α-CD8
Q
Ri (MΩ)
R
mPFC
Vgat / Gephy
S
No. Inhibitory Synapse
T
Vgat coverage
U
Gephyrin coverage

### Figure S10

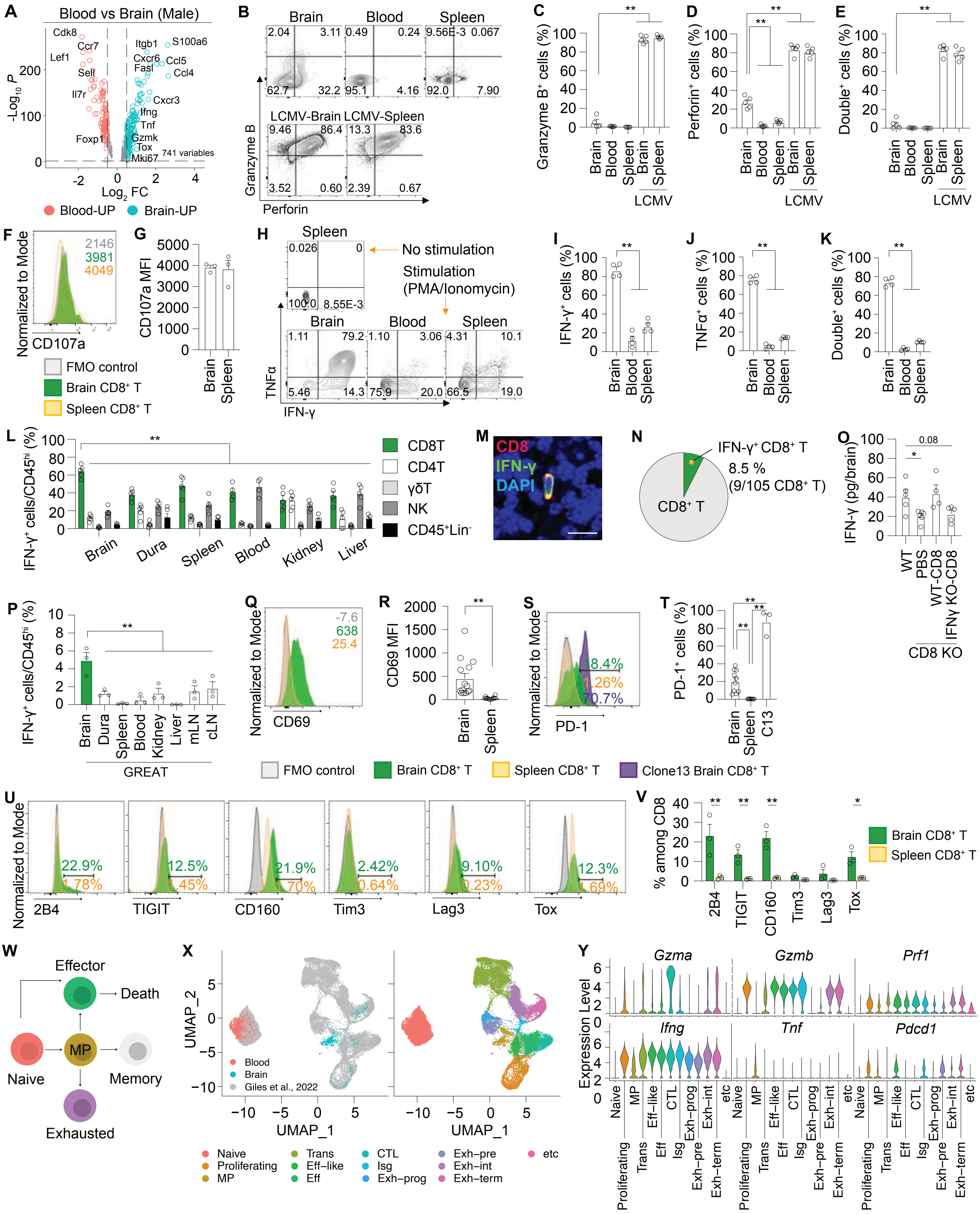

A
Blood vs Brain (Male)
Cdk8
Ccr7
Lef1
Sell
Il7r
Foxp1
Itgb1
S100a6
Cxcr6
Ccl5
Fasl
Ccl4
Cxcr3
Ifng
Tnf
Gzmk
Tox
Mki67
741 variables
-Log10 P
Log2 FC
Blood-UP
Brain-UP
B
Brain
Blood
Spleen
LCMV-Brain
LCMV-Spleen
Granzyme B
Perforin
C
Granzyme B+ cells (%)
D
Perforin+ cells (%)
E
Double+ cells (%)
Brain
Blood
Spleen
LCMV
F
Normalized to Mode
CD107a
2146
3981
4049
FMO control
Brain CD8+ T
Spleen CD8+ T
G
CD107a MFI
H
Spleen
No stimulation
Stimulation (PMA/Ionomycin)
TNFα
IFN-γ
I
IFN-γ+ cells (%)
J
TNFα+ cells (%)
K
Double+ cells (%)
L
IFN-γ+ cells/CD45hi (%)
CD8T
CD4T
γδT
NK
CD45+Lin-
Brain
Dura
Spleen
Blood
Kidney
Liver
M
CD8
IFN-γ
DAPI
N
IFN-γ+ CD8+ T
8.5 %
(9/105 CD8+ T)
CD8+ T
O
IFN-γ (pg/brain)
0.08
WT
PBS
WT-CD8
IFNγ KO-CD8
CD8 KO
P
IFN-γ+ cells/CD45hi (%)
Brain
Dura
Spleen
Blood
Kidney
Liver
mLN
cLN
GREAT
Q
Normalized to Mode
CD69
-7.6
638
25.4
R
CD69 MFI
S
Normalized to Mode
PD-1
18.4%
1.26%
70.7%
T
PD-1+ cells (%)
C13
FMO control
Brain CD8+ T
Spleen CD8+ T
Clone13 Brain CD8+ T
U
Normalized to Mode
2B4
TIGIT
CD160
Tim3
Lag3
Tox
V
% among CD8
W
Effector
Death
Naive
MP
Memory
Exhausted
X
UMAP_2
UMAP_1
Blood
Brain
Giles et al., 2022
Naive
Proliferating
MP
Trans
Eff-like
Eff
CTL
Isg
Exh-prog
Exh-pre
Exh-int
Exh-term
etc
Y
Gzma
Gzmb
Prf1
Ifng
Tnf
Pdcd1
Expression Level

### Figure S11

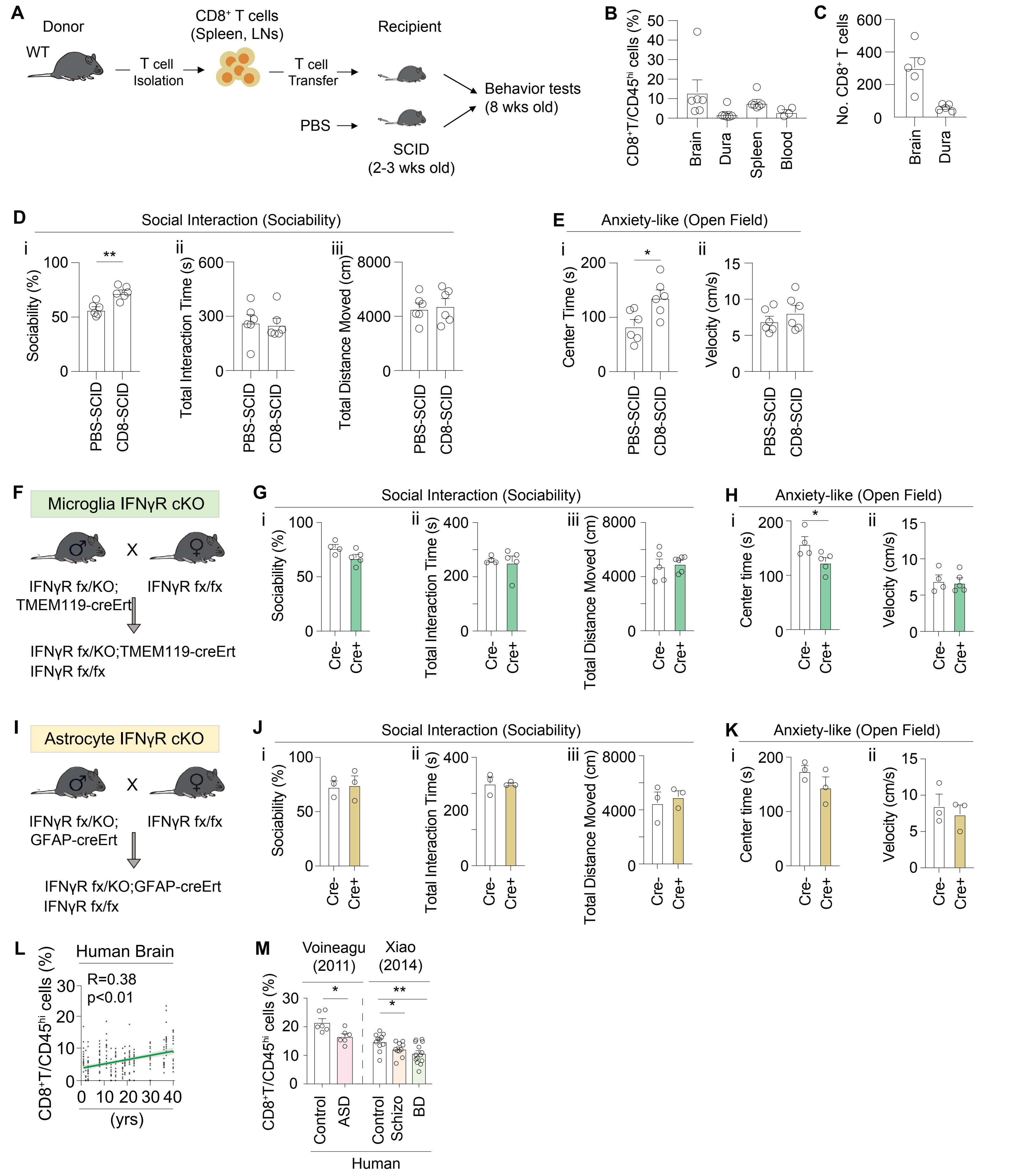

A
Donor
WT
T cell Isolation
CD8+ T cells (Spleen, LNs)
T cell Transfer
Recipient
PBS
SCID (2-3 wks old)
Behavior tests (8 wks old)
B
CD8+T/CD45hi cells (%)
Brain
Dura
Spleen
Blood
C
No. CD8+ T cells
Brain
Dura
D
Social Interaction (Sociability)
i
Sociability (%)
ii
Total Interaction Time (s)
iii
Total Distance Moved (cm)
PBS-SCID
CD8-SCID
E
Anxiety-like (Open Field)
i
Center Time (s)
ii
Velocity (cm/s)
F
Microglia IFNγR cKO
IFNγR fx/KO; TMEM119-creErt
IFNγR fx/fx
IFNγR fx/KO;TMEM119-creErt
IFNγR fx/fx
G
Social Interaction (Sociability)
Sociability (%)
Total Interaction Time (s)
Total Distance Moved (cm)
Cre-
Cre+
H
Anxiety-like (Open Field)
Center time (s)
Velocity (cm/s)
I
Astrocyte IFNγR cKO
IFNγR fx/KO; GFAP-creErt
IFNγR fx/fx
IFNγR fx/KO;GFAP-creErt
IFNγR fx/fx
J
Social Interaction (Sociability)
K
Anxiety-like (Open Field)
L
Human Brain
R=0.38
p<0.01
CD8+T/CD45hi cells (%)
(yrs)
M
Voineagu (2011)
Xiao (2014)
CD8+T/CD45hi cells (%)
Control
ASD
Control
Schizo
BD
Human
